# Cardiorespiratory physiological perturbations after acute smoke-induced lung injury and during extracorporeal membrane oxygenation support in sheep

**DOI:** 10.1101/058511

**Authors:** Saul Chemonges

## Abstract

**Background:** Numerous successful therapies developed for human medicine involve animal experimentation. Animal studies that are focused solely on translational potential, may not sufficiently document unexpected outcomes. Considerable amounts of data from such studies could be used to advance veterinary science. For example, sheep are increasingly being used as models of intensive care and therefore, data arising from such models must be published. In this study, the hypothesis is that there is little information describing cardiorespiratory physiological data from sheep models of intensive care and the author aimed to analyse such data to provide biological information that is currently not available for sheep that received extracorporeal life support (ECLS) following acute smoke-induced lung injury.

**Methods:** Nineteen mechanically ventilated adult ewes undergoing intensive care during evaluation of a form of ECLS (treatment) for acute lung injury were used to collate clinical observations. Eight sheep were injured by acute smoke inhalation prior to treatment (injured/treated), while another eight were not injured but treated (uninjured/treated). Two sheep were injured but not treated (injured/untreated), while one received room air instead of smoke as the injury and was not treated (placebo/untreated). The data were then analysed for 11 physiological categories and compared between the two treated groups.

**Results:** Compared with the baseline, treatment contributed to and exacerbated the deterioration of pulmonary pathology by reducing lung compliance and the arterial oxygen partial pressure to fractional inspired oxygen (PaO_2_/FiO_2_) ratio. The oxygen extraction index changes mirrored those of the PaO_2_/FiO_2_ ratio. Decreasing coronary perfusion pressure predicted the severity of cardiopulmonary injury.

**Conclusions:** These novel observations could help in understanding similar pathology such as that which occurs in animal victims of smoke inhalation from house or bush fires, aspiration pneumonia secondary to tick paralysis and in the management of the severe coronavirus disease 2019 (COVID-19) in humans.

## Introduction

During multifaceted experiments involving intensive care in large animal models in translational research, information related to animal monitoring is often collected with varying accuracy, scope and end-user applications. Data collection can be manual, electronic, or both^1–3^. Manually input data can include subjectively scored end points, like the plane of anaesthesia, and objective data such as heart rate or breaths per minute. Depending on the goals of the study, certain information may be used to validate or test novel therapies, or to understand and refine existing treatments. In certain cases, experimental information may be collected for scientific curiosity or for ‘classified’ use, and outcomes may never be publicly available, particularly if the results are negative.

The source of data for this study was from a sheep model^2–5^ in which sheep were treated for acute smoke-induced acute lung injury using veno-venous (VV) extracorporeal membrane oxygenation (ECMO) ^2^, a form of extracorporeal life support (ECLS) developed to complement the treatment of acute lung injury in humans^6–8^. During this type of ECLS, venous blood is carried from the patient to a gas exchange device where the blood is enriched with oxygen, has carbon dioxide is removed, and oxygenated blood is returned to the patient’s circulation in the right atrium. This method can be used for treatment, as respiratory support during lung transplantation, and in critically ill patients with potentially reversible respiratory failure^8, 9^. The multiple advanced cardiovascular^3^, respiratory, patient point-of-care procedures and instrumentation associated with ECLS even in animal experimentation is highly data- and equipment-intensive. This platform is useful for developing research and methodological skills for *in vivo* animal instrumentation and for the processing of large, real-time clinical data sets from multifaceted animal studies that can be applied to similar intensive care scenarios. An opportunity to develop these skills arose within a source study conducted at Queensland University of Technology and The University of Queensland, which was an ongoing publicly funded animal experimentation study. While the objectives of the primary study had a separate focus, there were considerable amounts of redundant raw data with potential use in veterinary science and other disciplines, once processed.

Although burn and smoke inhalation ECLS models in sheep have been around for many years, most studies have only focused on their translational potential for applications in human medicine and not for veterinary science applications or further refinement of the model. For example, it has been documented that prolonged exposure to smoke exacerbates lung injury after evaluating the manner various types of exposure to smoke chemicals cause injury^10^. Another sheep ECMO study investigated the pathophysiology of circulating leukocytes, oxygen free-radical activity, thromboxane release and respiration^11^. In the preceding study, animals treated with smoke injury followed by ECMO had significantly increased circulating thromboxane B2 levels and oxygen free-radical activity compared with controls and animals treated with smoke and mechanical ventilation but details of haemodynamics were not presented.

In the present study, the author hypothesised that there is little information describing physiological data from multifaceted sheep models of intensive care and the author aimed to analyse such data to provide cardiorespiratory biological information that is not currently available in sufficient detail for sheep that receive ECLS following acute smoke-induced acute lung injury. The overall goal was to provide useful information relevant to the sheep model itself as well as to those interested in broad animal experimentation and veterinary medicine in general. The specific objective was to utilise the raw data from the sheep ECMO model study and analyse that data to provide biological information that is not currently available for sheep that receive ECLS following acute smoke-induced lung injury to further understand the physiology of cardiorespiratory support.

## Materials and methods

### Ethics statement

Animals were obtained and treated in accordance with the Australian Code of Practice for the Care and Use of Animals for Scientific Purposes^12^, with strict adherence to published and currently acceptable guidelines on using experimental animals and reporting findings^13, 14^. All studies were registered with institutional animal welfare and ethics departments; moreover, the Queensland University of Technology Animal Ethics Approval No. 110000053 was obtained and it was ratified by The University of Queensland.

### Experimental animals

Batches of approximately 2-year old healthy adult merino ewes *(Ovis aries)* were obtained and carted to Brisbane from the Australian Commonwealth Scientific and Industrial Research Organisation (CSIRO) breeding facility in Armidale, New South Wales. The sheep were agisted as a flock in an open pasture farm to be used as part of, and a continuation of previously described studies^15^. The farm had improved pastures and the sheep had natural shade from trees with free access to water before being transferred within two weeks of experiments to a purpose-built animal experimental laboratory at QUT-MERF^1, 15^. The sheep were handled as per standard humane operating procedures that have been described in detail elsewhere^1^. The experiments were conducted at the Biological Research Facility (BRF) housed within the Medical Engineering Research Facility of Queensland University of Technology (QUT-MERF) in Brisbane. Animal selection and experimental procedures for this, and also other multifaceted studies that used similar methods have previously been described^15–17^. Briefly, the sheep were fed proprietary sheep pellets, lucerne and had free access to water. Shelter was provided in built concrete-floored sheds in which the sheep had free access. Shade was also provided by large trees in the paddocks and the sheep interacted freely with each other. Animals were fasted for approximately 24 h with free access to drinking water until two hours before the commencement of the experimental procedure at 8:00 – 9:00 a.m. on the day of the experiment. Local guidelines mandated the presence of a team comprising of a veterinarian and other trained personnel to monitor the sheep at all times for the duration of the experiment for animal welfare purposes and for the mitigating staff fatigue. If at any time the animal became physiologically distressed to such an extent that it could not be managed or reversed, mechanisms were in place to have the sheep be immediately euthanased and documented accordingly ^1^.

A sample size calculation was performed for the entire original sheep ECMO project which comprised 72 sheep (9 groups of 8 sheep each), as previously reported^1^. There was a subsequent addition of two control groups (two groups of eight sheep each), bring the total of sheep to 88. The other experimental groups were out of scope, as they investigated other aspects of ECLS such as blood transfusion studies, therefore, this study focuses on the cardiorespiratory physiology (4 groups) only. In this study, 19 sheep were pseudo-randomised into four experimental groups (Table 1). The experimental groups were classified based on the following two aspects: the duration of treatment (24 hours of ECMO only—E24H); treatment after smoke inhalation (injury) (24 hours of ECMO after smoke inhalation—SE24H). Two additional groups included one group that received smoke inhalation injury but no treatment (24 hours of monitoring only after smoke inhalation and no ECMO; SC24H), and another group that inhaled room air only as the injury (placebo) and no treatment (24 hours of monitoring, no smoke inhalation and no ECMO; C24H). Robust data was acquired from 16 sheep (E24H and SE24H) and included fully in the study; however, data of three sheep from groups SC24H and C24H was considered only as early observational data – within the timeline required to satisfy the requirements of a research degree at The University of Queensland back then. A systematic approach was developed for processing the data. (All raw and processed data can be downloaded at https://doi.org/10.5061/dryad.3r2280gd5.)

**Table 1.** Characteristics of treated and untreated groups (control experiments) of sheep

| **Experiment Group** | **Date of experiment** | **Sheep No.** | **Age (Y)** | **Weight (kg)** | **Length of Sheep (cm)** | **BSA** |
| --- | --- | --- | --- | --- | --- | --- |
| E24H | 06/10/2011 | E24H-01/390 | 2 | 50 | 110 | 1.29 |
|  | 20/10/2011 | E24H-02 | 2 | 47.6 | 110 | 1.25 |
|  | 17/11/2011 | E24H-03 | 2 | 51 | 110 | 1.31 |
|  | 01/03/2012 | E24H/4616 | 2 | 50 | 110 | 1.29 |
|  | 29/03/2012 | E24H-05/4627 | 2 | 47 | 110 | 1.24 |
|  | 04/04/2012 | E24H-06/4146 | 2 | 40 | 110 | 1.11 |
|  | 12/04/2012 | E24H-07/4032 | 2 | 52.5 | 110 | 1.34 |
|  | 03/05/2012 | E24H-08/4630 | 2 | 53 | 110 | 1.34 |
| SE24H | 02/02/2012 | SE24H-01/4139 | 2 | 44 | 110 | 1.19 |
|  | 09/02/2012 | SE24H-02/4542 | 2 | 53 | 110 | 1.34 |
|  | 16/02/2012 | SE24H-03/4280 | 2 | 45.5 | 110 | 1.21 |
|  | 23/02/2012 | SE24H-04/4624 | 2 | 50 | 110 | 1.29 |
|  | 17/05/2012 | SE24H-05/4458 | 2 | 55 | 140 | 1.38 |
|  | 24/05/2012 | SE24H-06/8461 | 2 | 46 | 140 | 1.22 |
|  | 24/01/2013 | SE24H-07/09C8032 | 3 | 52 | 130 | 1.33 |
|  | 21/02/2013 | SE24H-09A0142 | 2 | 50 | 140 | 1.29 |
| SC24H | 18/06/2013 | SC24H-01 | 2 | 51 | 140 | 1.31 |
|  | 27/06/2013 | SC24H-02 | 2 | 57 | 140 | 1.41 |
| C24H | 08/08/2013 | C24H-01 | 2 | 53 | 140 | 1.34 |

(placebo/untreated).

### Experimental procedures

#### Critical care of animals, VV ECMO setup and physiological data acquisition

The details of animal selection, care, and pre-anaesthetic processes; anaesthesia technique; airway access and ventilation; instrumentation for VV ECMO; haemodynamic monitoring; respiratory monitoring; temperature, fluids, vasoactive drug administration, and electrolyte management; blood collection; physiological data acquisition; and the technique for euthanasia of the sheep after the experiments have previously been described in a detailed protocol^1^. In brief, the sheep was restrained in a sling cage and the ventral neck region was aseptically prepared to enable intravenous access. For VV ECMO implementation, venous blood was accessed from the right jugular vein of the animal and then oxygenated and returned to the right atrium of the heart after it 6 was made to pass through an oxygenator. For the combined purpose of blood sampling, administration of medications, and fluid administration, a multi-lumen central venous catheter was inserted into the left jugular vein of the animal under local anaesthesia. The left jugular vein was also cannulated with an 8G sheath for the insertion of a pulmonary artery catheter for haemodynamic monitoring. In addition, an 11G sheath catheter was then inserted proximally into the left jugular vein for intra-cardiac echocardiography catheter insertion. The right jugular vein was cannulated both proximally and distally with single lumen central lines to aid insertion of return and access ECMO cannulas, respectively. All animals were intubated and received mechanical ventilation as previously described^1^. Briefly, the initial ventilator tidal volume was set to approximately 10 mL/kg with a respiratory rate of 15 breaths/min, positive end expiratory pressure (PEEP) of 5 cm H_2_O, and an initial F_i_O_2_ (fraction of inspired oxygen) of 1.0. These settings were then titrated based on arterial blood gas results. A low tidal volume—high PEEP strategy was used to minimise ventilator-induced lung injury.

In order to obtain high-quality cardiorespiratory monitoring data, the instrumentation of the sheep was undertaken to acquire and derive the following physiological parameters using established standard methods at defined timepoints: core body temperature (T), pO_2_, SpO_2_, alveolar–arterial oxygen gradient P(A-a)O_2_, PaO_2_/FiO_2_ ratio, end-tidal carbon dioxide concentration (etCO_2_), heart rate (HR), arterial blood pressure (BP) (systolic and diastolic), mean arterial BP (MAP), pulmonary artery pressure (PA) (systolic and diastolic), mean pulmonary artery pressure (MPAP), central venous pressure (CVP), pulmonary artery occlusion pressure (PAOP), mixed venous oxygen saturation (SvO_2_), stroke volume (SV), continuous cardiac output (CCO), cardiac index (CI), systemic vascular resistance (SV), systemic vascular resistance index (SVRI), pulmonary vascular resistance index (PVRI), left ventricular stroke work index (LVSWI), right ventricular stroke work index (RVSWI), coronary perfusion pressure (CPP), arterial oxygen content (CaO_2_), and oxygen delivery index (O_2_EI). (These methods are detailed in data files that can be downloaded at https://doi.org/10.5061/dryad.3r2280gd5.)

#### Smoke inhalation injury

In the original study of the ECMO model^4^, sheep inhaled standardised cotton smoke generated by a device that combusts material in an oxygen-deficient environment as previously described^18^. In brief, 8 g of cotton towelling was combusted in a chamber with transparent walls and 400 ml tidal volume. One tidal volume breath (approximately 10–12 ml/kg) of the smoke was delivered to the sheep via plastic tubing that was 1 m long connected to a tracheostomy tube. A fixed number (12) of breaths were given with each load of cotton over a period of approximately one minute. Serial arterial blood gas samples were taken to assess the effect of smoke inhalation, beginning at a predetermined time point after the smoke breath cycles.

### Physiological data management

Raw data were obtained from critical care monitoring of sheep undergoing treatment for acute smoke-induced lung injury that involved several separate previous projects. Data analysed were collected prior to 23 August 2013 and were kindly obtained from two of the scientists (Diab, S. and Dunster, K.R., who developed the base model – see reference^4^), as part of a research higher-degree project of the author at The University of Queensland^19, 20^. All data files were stored in Microsoft^®^ Excel 97–2003 (Microsoft Corporation, Redmond, WA, USA) format and were grouped per sheep and date of the experiment. Data comprised separate files of real-time physiological data recorded in the hard drives of the monitoring devices (electronically acquired data) and parameters manually recorded by those monitoring the sheep under anaesthesia (manually acquired data)— which included data from the electronic monitoring equipment—as back-up if the electronic monitors malfunctioned.

### Manually acquired physiological data workflow

A clone of the master manual data entry Excel spreadsheet was created by excluding the formatting and formulas. Several members of the sheep ECMO research team repeatedly inspected the data for errors to in order to ensure that all columns, rows, time points, and data points had been copied correctly, including number formats. Redundant columns were omitted from the spreadsheet and data were curated and aligned to predetermined experimental time points. While maintaining the same experimental time point headers on the spreadsheet, data were grouped into the following categories: ventilator settings, blood pressure and haemodynamics, fluids and urine output, arterial blood gas values, activated clotting time, anaesthetics, anticoagulants, and ECLS circuit observations.

### Electronically acquired physiological data workflow

Electronically acquired physiologic monitoring raw data were inspected for completeness. The data comprised 36 time points: ECLS pump time (min); time of day (h); electrocardiograph (heart rate); arterial blood pressure (mean, systolic, diastolic, heart rate); central venous pressure (mean); pulmonary artery pressure (mean, systolic, diastolic); oxygenator pressure (pre- and post-); capnography (etCO_2_, respiratory rate); pulse oximetry (SpO_2_, heart rate); ECLS pump (flow rate, speed); ventilator (mode, frequency, oxygen, pressure control, inspiratory volume, expiratory volume, expiratory minute volume, pressure maximum, mean pressure, positive end-expiratory pressure, plateau pressure, inspiratory resistance, expiratory pressure, pulmonary compliance, inspiratory flow); mixed venous oxygen saturation (SvO_2_); and continuous cardiac output (CCO). A baseline time point was established after instrumentation and an injury time point corresponding to the smoke inhalation time point was determined thereafter. It is important to note that there may or may not have been any data at any given point in time. The electronically acquired physiological monitoring data were inspected for errors and cleaned to provide data for downstream analysis.

### Respiratory efficiency and haemodynamic monitoring

Manually acquired observations at specifically designated timepoints were recorded and extracted as detailed in the data filed at doi:10.5061/dryad.3r2280gd5. These timepoints were at baseline (soon after instrumentation of the sheep), smoke injury, 5 min post smoke injury, 1 hour post smoke injury. This was then followed by ECMO treatment, which was recorded in the following manner: 0, 0.25, 1, 1.5, 2, 4, 6, 6.5, 7, 8, 10, 12, 14, 16, 18, 20, 22 and 24 hours of ECMO.

### Pre-data analysis checks

Thereafter, data were subjected to further integrity checks. An important step was to make a plot of data versus time together with descriptive statistics for all data points in the grouped data. After artefact elimination and integrity checks, data for individual sheep were assigned to six categories: activated clotting time; anaesthetics + inotropes and anticoagulants; arterial blood gas values; blood pressure + ventilation and haemodynamic data; calculated respiratory + haemodynamic variables; and fluids and urine production. Using specially written macros, data were extracted from each experiment and grouped by parameters corresponding to experimental time points. All sheep treatment data were then filed according to parameter.

Data from the 19 sheep from groups E24H, SE24H, C24H and SC24H were processed further. Data integrity checks were performed again and repeated by several sheep ECMO research team members. The treatment timeline comprised 22 time points for all experiments in which sheep received acute smoke-induced lung injury (SE24H). A trend plot and descriptive statistics panel in Excel were used for data quality control processes for suitability for downstream data analysis and end-user applications.

### Statistical methods

In order to meet the specific objective of the study, data from the groups, uninjured/treated and injured/treated groups were analysed after testing for normality using D’Agostino–Pearson omnibus normality test. The means, medians and standard deviations of the weights of the sheep, where applicable, were tabulated and graphically compared. The physiological parameters of the groups were charted and compared with each other using one-way analysis of variance (ANOVA), where appropriate and significance was reported based on Brown-Forsythe test. Further, parameters between groups were compared using a paired two-tailed t-test. All p-values were two-sided and p <0.05 was considered statistically significant. All statistical calculations were performed using GraphPad PRISM 6 software (GraphPad Software, La Jolla, CA, USA). An earlier version of this article can be found on bioRxiv (doi: https://doi.org/10.1101/058511).

### Results

The biodata of the sheep that were used in the current analysis are presented in the *Methods* section (see Table 1). The weights of the uninjured/treated sheep, unlike the injured/treated group, did not pass the D’Agostino–Pearson omnibus normality test; however, there was no significant difference in the weights of the sheep between the groups (Figure 1).

**Figure 1.**
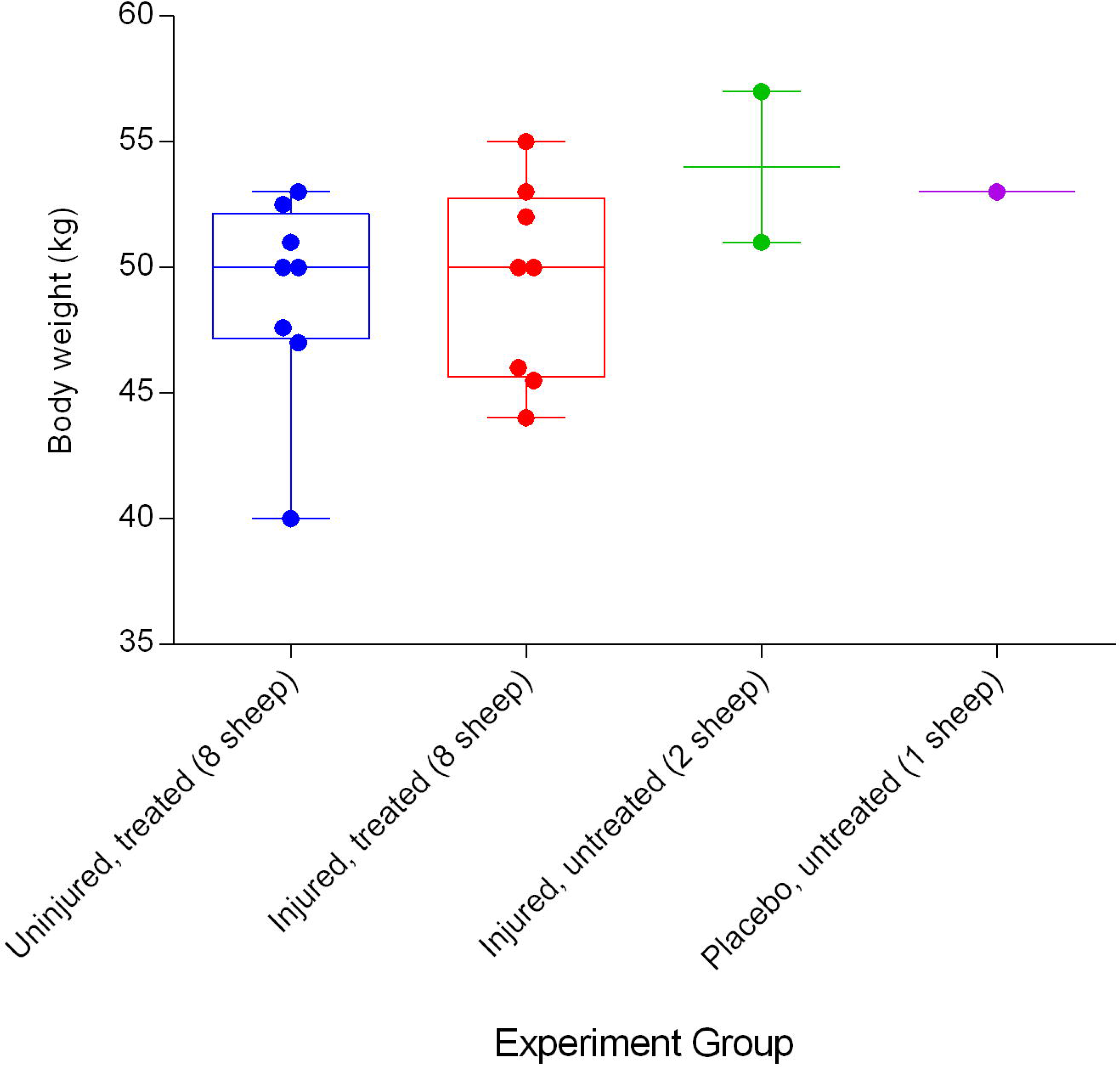
Body weights (Mean ± SD) of smoke and non-smoke injured sheep that received extracorporeal life support (ECLS) as compared to those of untreated controls.

### Mechanical ventilation

A decrease in pulmonary compliance was found in all of the sheep during the course of the experiments, with the injured/treated (SE24H) animals having the most severe and drastic decrease followed by the uninjured/treated (E24H), injured/untreated (SC24), and placebo/untreated (C24) sheep in that order (Figure 2). There was a significant difference (*p* = 0.0013) in pulmonary compliance between uninjured/treated and injured/treated groups. The injured/treated sheep had consistently lower SpO_2_ compared with the other groups, but there was no significant difference in SpO_2_ readings between the groups (Figure 3). Further, there was an initial increase in etCO_2_ followed by a rapid decrease that reduced 15 minutes after the treatment was began. The etCO_2_ of the injured sheep continued to trend downward and plateaued in the uninjured groups (Figure 4). There was a significant difference (*p* = 0.0147) in the etCO_2_ between the uninjured/treated and injured/treated groups.

**Figure 2.**
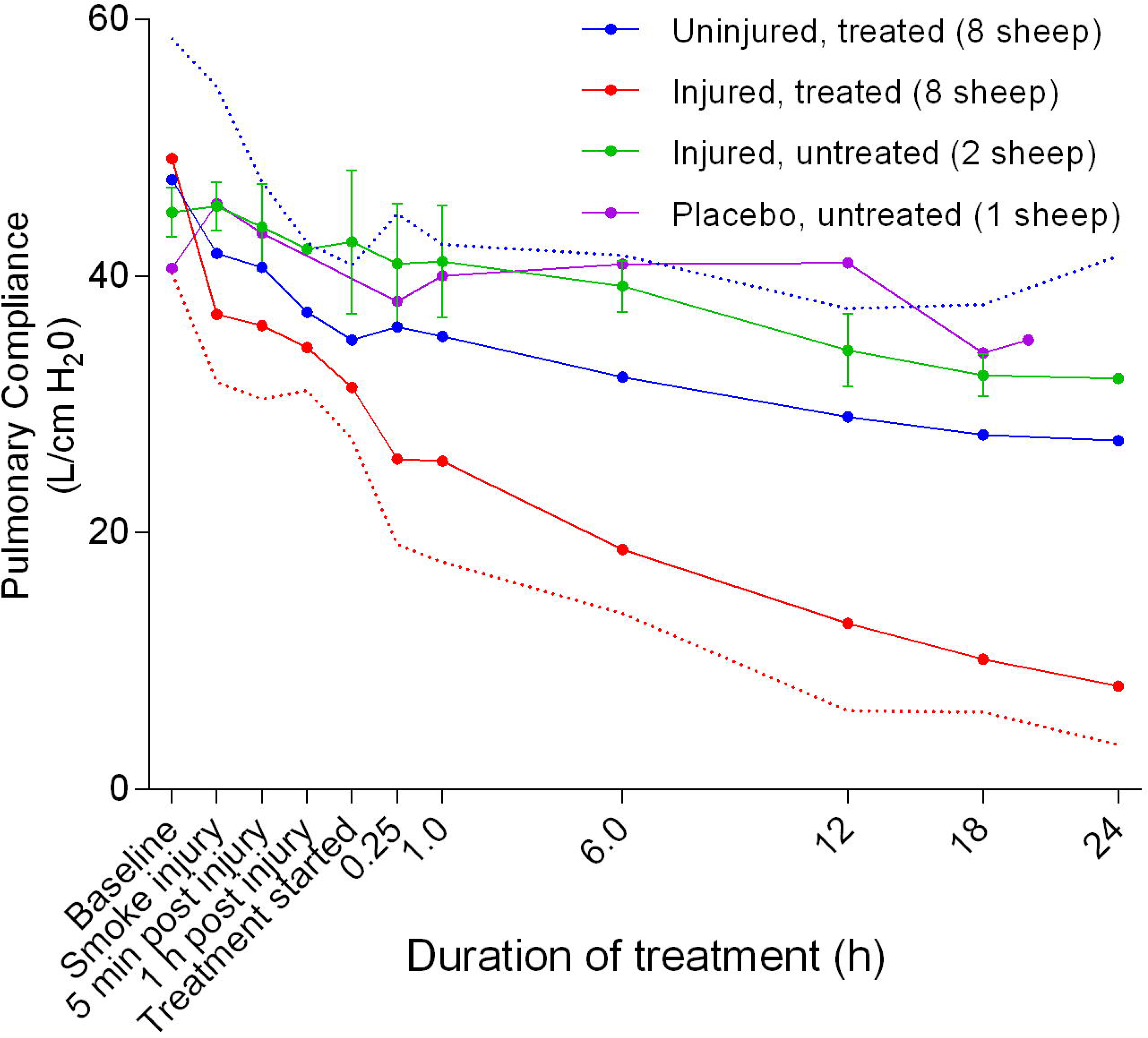
Pulmonary compliance (Mean ± SD) in smoke and non-smoke injured sheep that received extracorporeal life support as compared to that in untreated controls. Dotted lines represent error bar margins.

**Figure 3.**
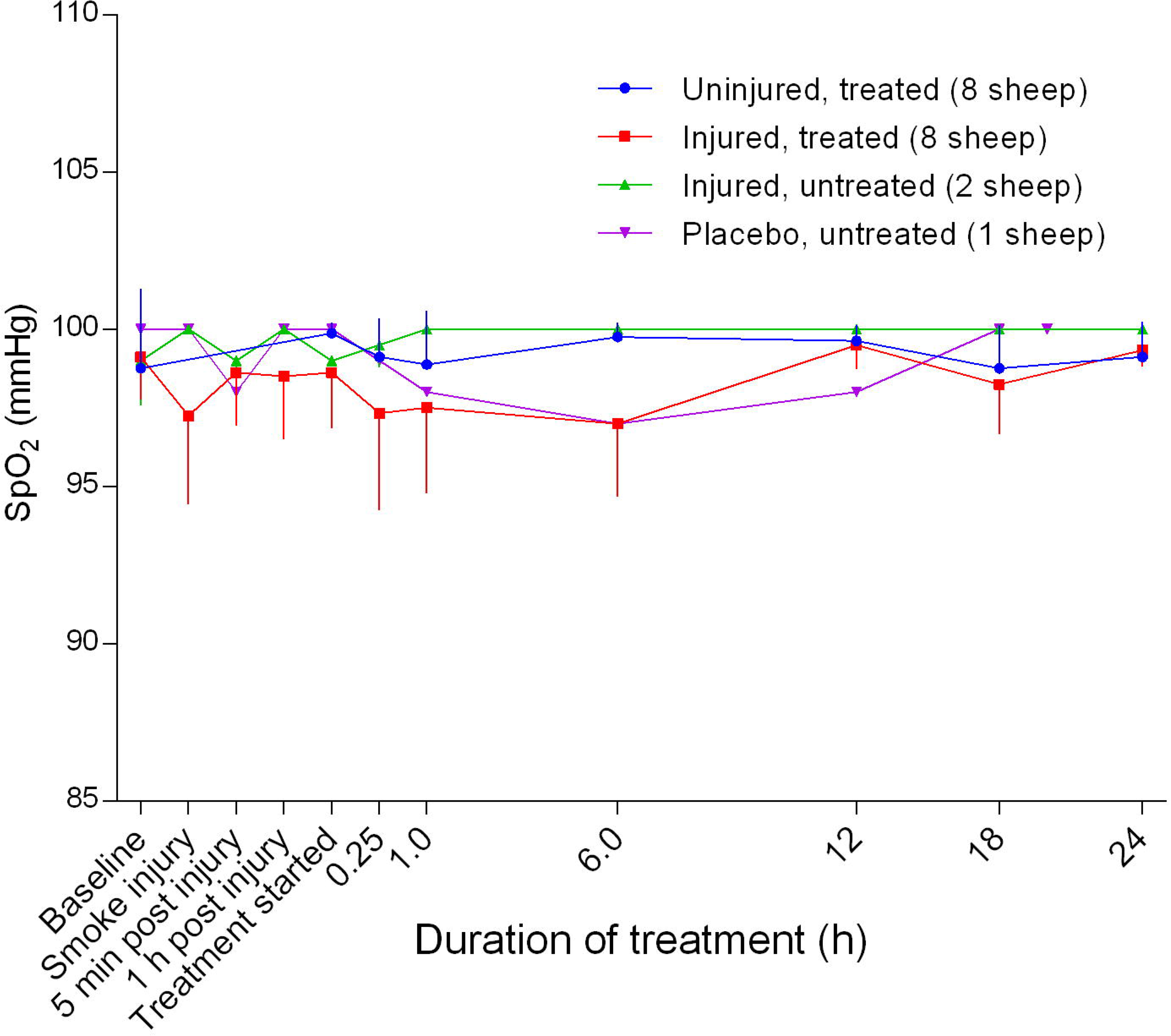
Arterial oxyhaemoglobin saturation (SpO_2_) (Mean ± SD) in smoke and non-smoke injured sheep that received extracorporeal life support as compared to that in untreated controls.

**Figure 4.**
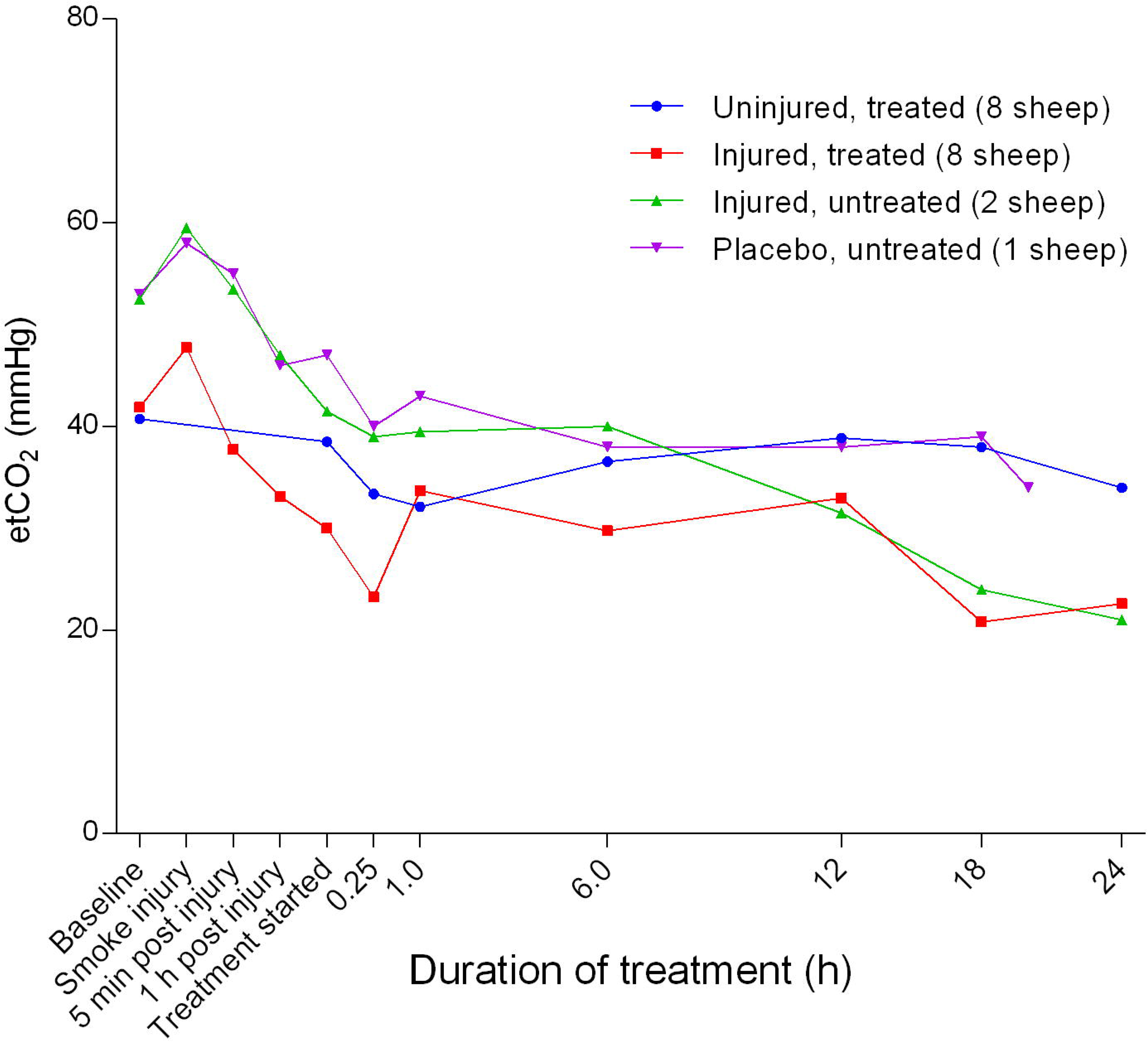
End-tidal carbon dioxide tension (etCO_2_) (Mean ± SD) in smoke and non-smoke injured sheep that received extracorporeal life support as compared to that in untreated controls.

### Arterial blood gas evaluation

Blood pH varied between the groups (Figure 5). The placebo/untreated sheep had the highest pH while the injured/treated group had the lowest. There was a significant difference in pH between the uninjured/treated and injured/treated groups (*p* = 0.0343). The pCO_2_ in all but the uninjured/treated sheep increased initially before plummeting sharply, thereby forming a shallow trough corresponding to 1 hour after the treatment, followed by a slight increase before stabilising in all sheep (Figure 6). There was a gradual decrease in pO_2_ in the treated groups of sheep from baseline before decreasing dramatically at the start of treatment with the injured sheep having the most profound decrease (Figure 7). However, there was no significant difference in pO_2_ between the uninjured/treated and injured/treated groups.

**Figure 5.**
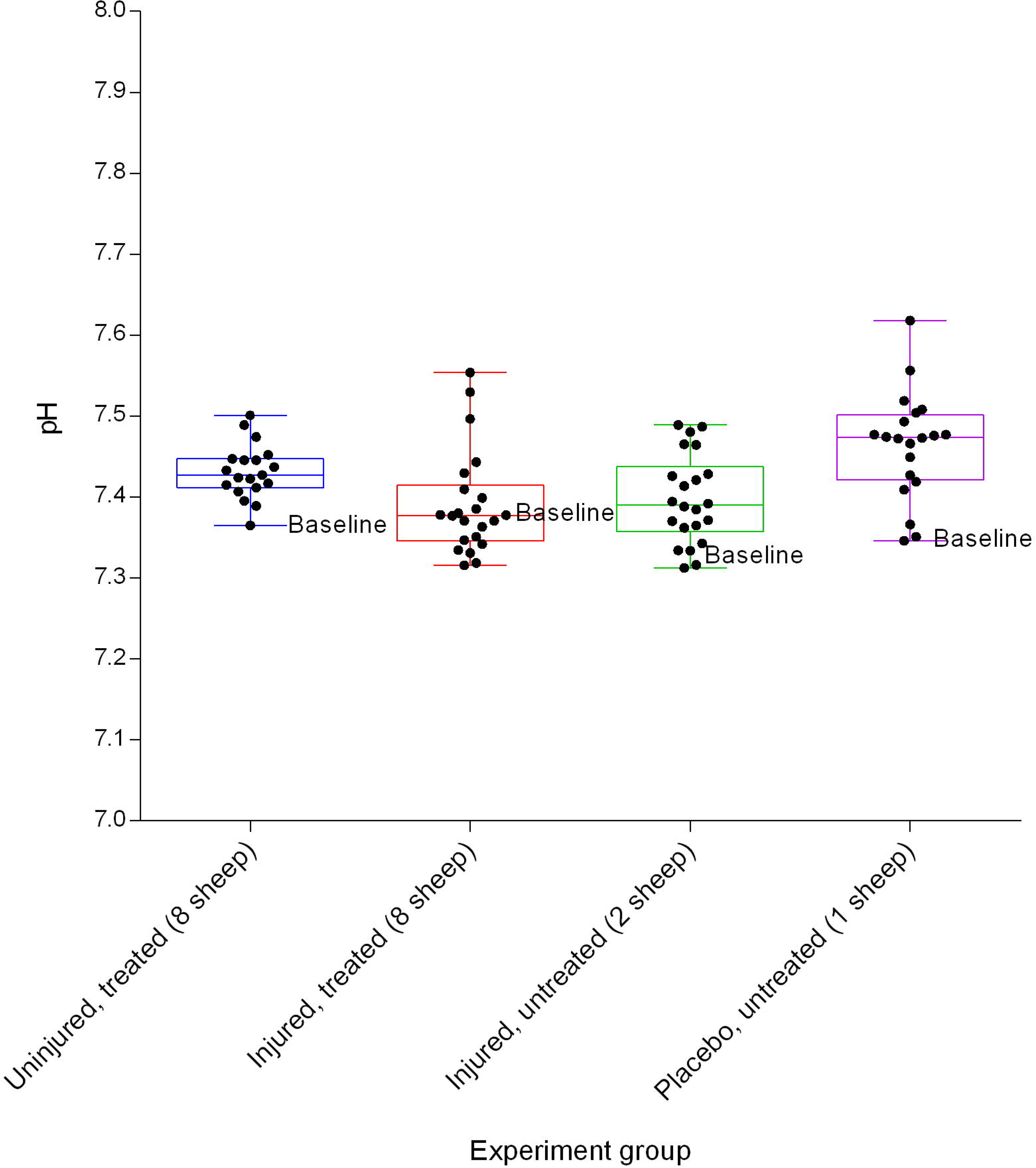
Arterial blood pH (Mean ± SD) over a 24h period in smoke and non-smoke injured sheep that received extracorporeal life support as compared to that in untreated controls. The dots represent hourly time-points.

**Figure 6.**
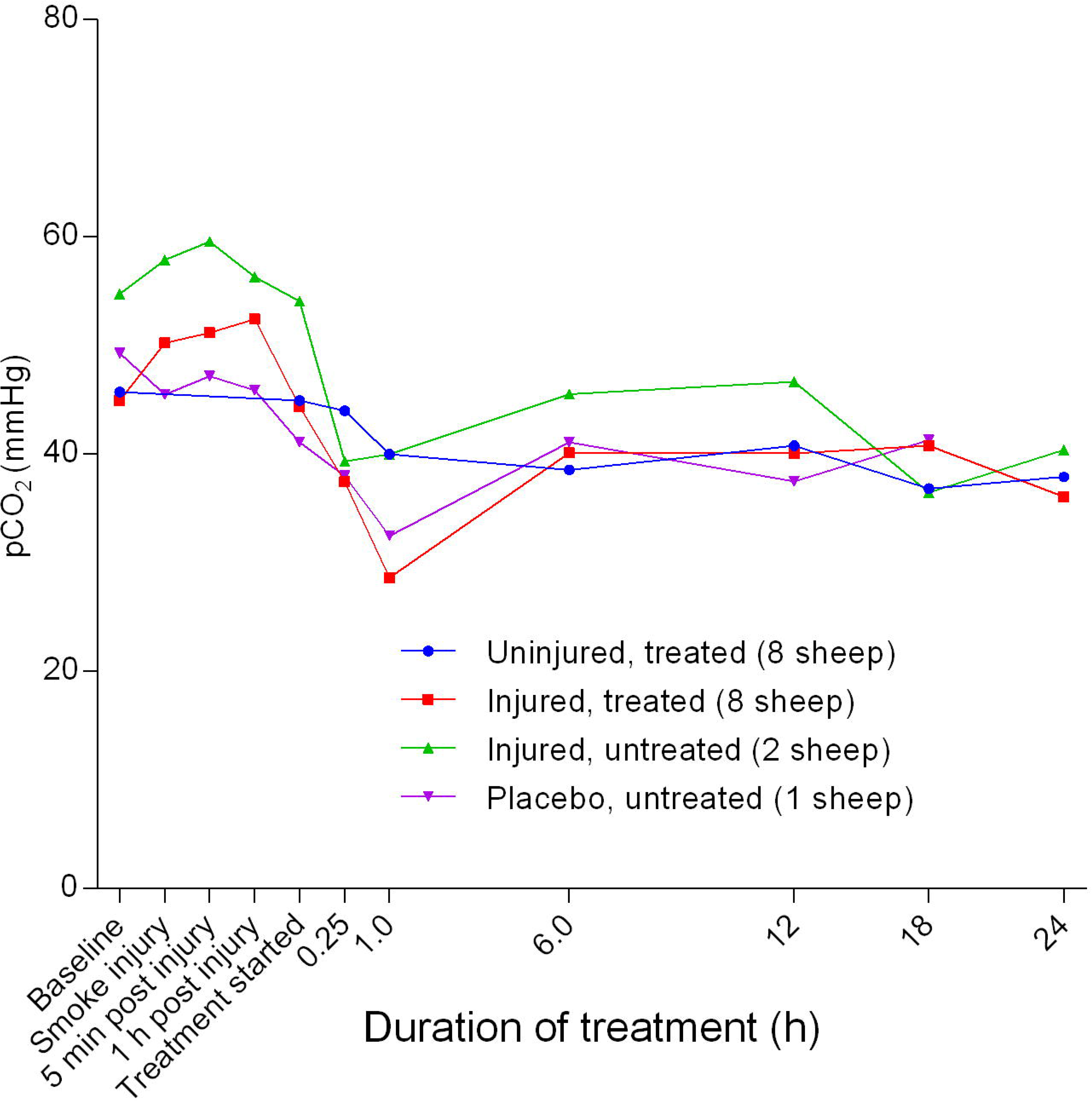
Mean arterial carbon dioxide partial pressure (pCO_2_) (Mean ± SD) in smoke and non-smoke injured sheep that received extracorporeal life support as compared to that in untreated controls.

**Figure 7.**
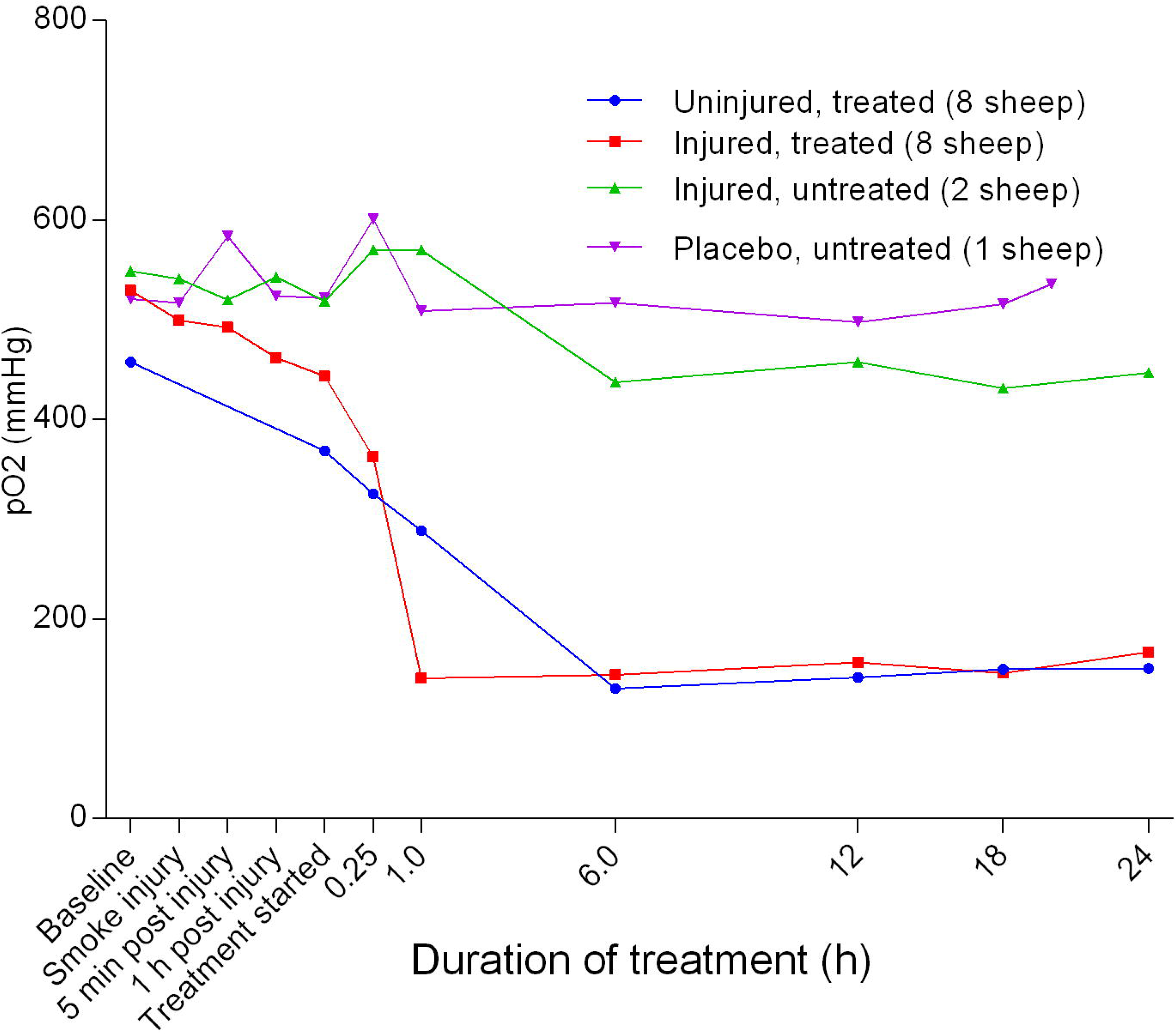
Arterial oxygen tension (pO_2_) (Mean ± SD with no error bars shown) in smoke and non-smoke injured sheep that received extracorporeal life support as compared to that in untreated controls.

### Haemoglobin dynamics

The concentration of haemoglobin [Hb] was found to decrease slightly from baseline before gradually increasing in the injured sheep and remained relatively constant over time in the uninjured sheep. There was a significant difference in [Hb] between the uninjured/treated and injured/treated (p= 0.0131) groups (Figure 8). The fraction of oxyhaemoglobin (FO_2_Hb) decreased sharply with the lowest reading at five minutes post-injury before returning to near baseline levels within 1 hour of the treatment (Figure 9). The injured/treated sheep had a considerably deeper trough in FO_2_Hb level and there was a significant difference (*p* = 0.046) between troughs. There was no change in FO_2_Hb for the uninjured sheep. Further, the fraction of carboxyhaemoglobin (FCOHb) increased sharply from baseline, peaking at approximately five minutes post-injury and decreased sharply thereafter to the beginning of treatment before gradually returning to near-baseline levels at approximately 6 hours after the treatment was begun in the injured sheep (Figure 10). The injured/treated sheep had a higher peak FCOHb than the injured/untreated sheep, although the difference was not significant. There was no change in FCOHb for the uninjured sheep. The fraction of methaemoglobin (MetHb) increased gradually from baseline, peaking at approximately five minutes post-injury and then gradually decreased when the treatment was begun (Figure 11). This was followed by a gradual 12 return to near-baseline levels at approximately 6 hours after the treatment was begun in the injured/treated sheep. There was no change in MetHb for the uninjured sheep. There was an initial subtle decrease in calculated haematocrit (Hct) before a steady increase in the injured sheep and relatively flat slopes for the uninjured sheep (Figure 12).

**Figure 8.**
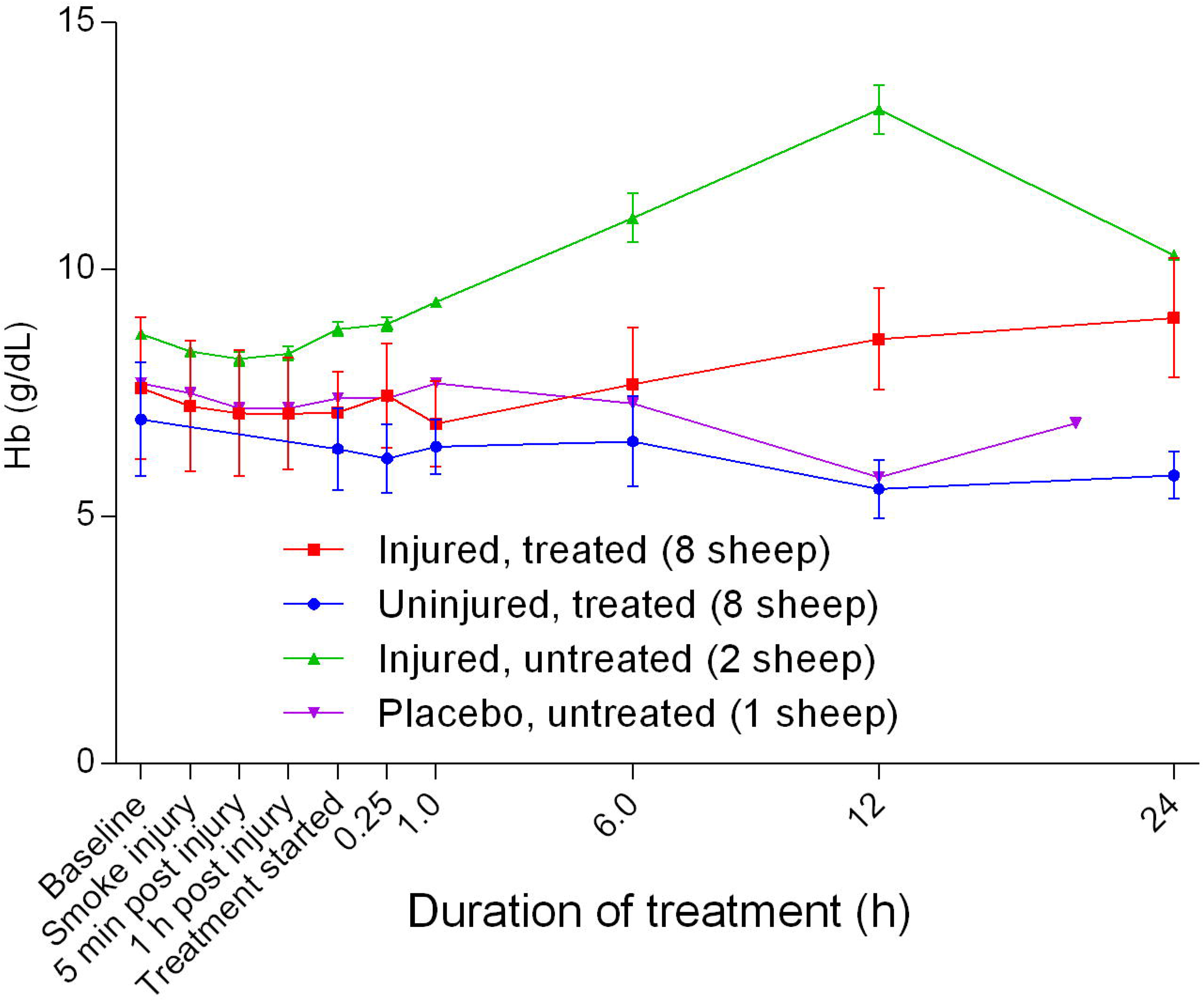
Haemoglobin concentration [Hb] (Mean ± SD) in smoke and non-smoke injured sheep that received extracorporeal life support as compared to that in treated controls.

**Figure 9.**
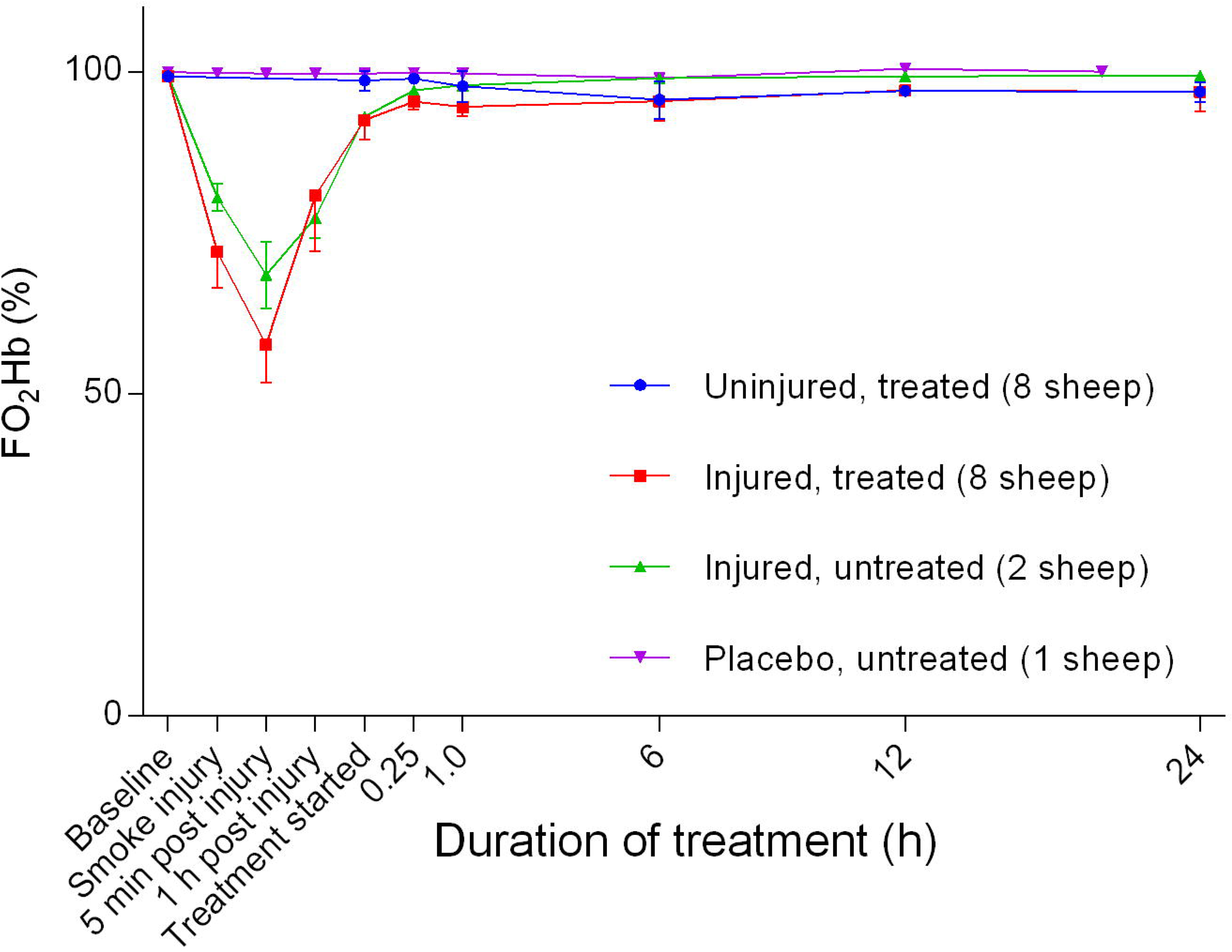
Fraction of oxyhaemoglobin (FO_2_Hb) (Mean ± SD) in smoke and non-smoke injured sheep that received extracorporeal life support as compared to that in untreated controls

**Figure 10.**
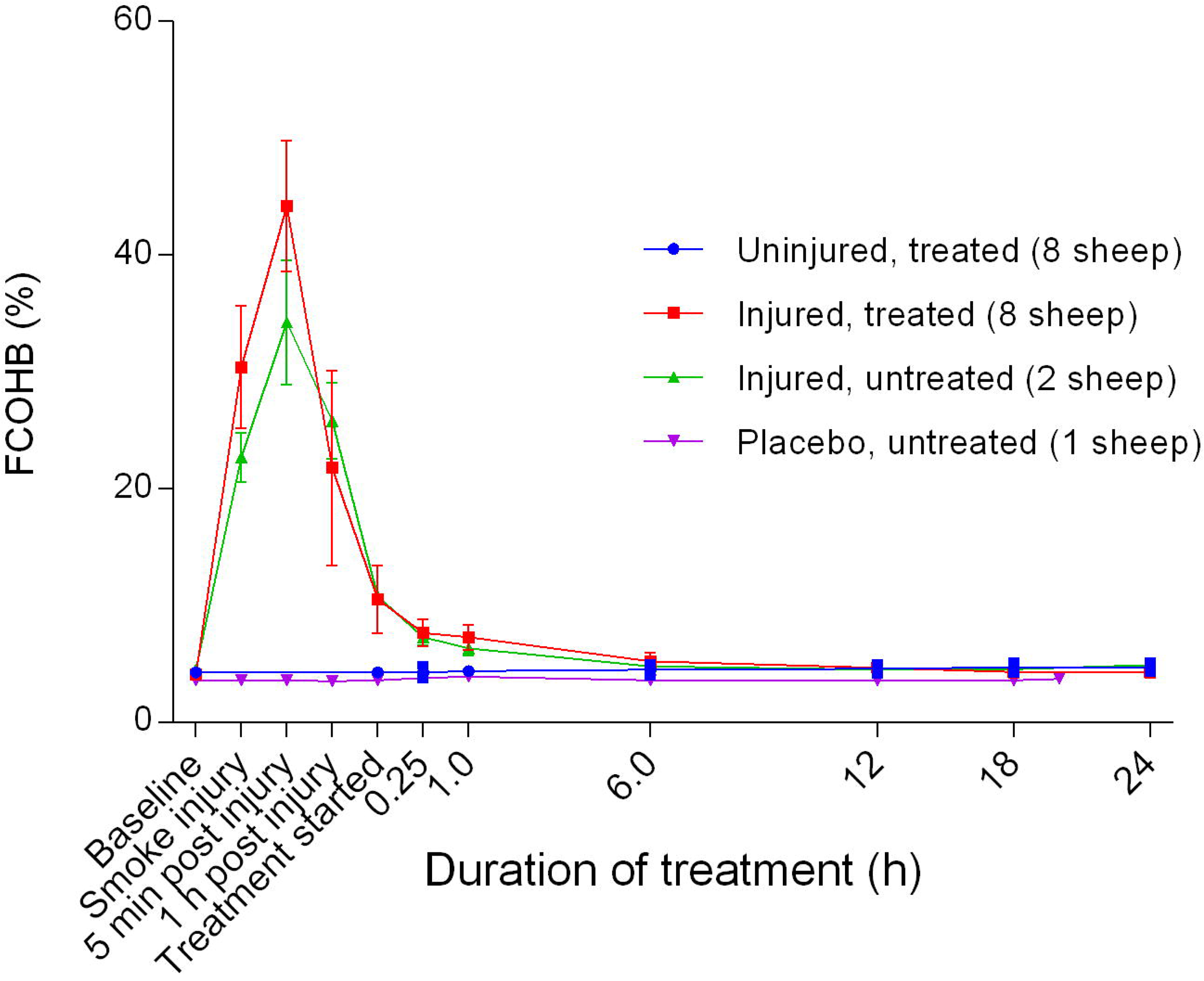
Fraction of carboxyhaemoglobin (FCOHb) (Mean ± SD) in smoke and non-smoke injured sheep that received extracorporeal life support as compared to that in untreated controls.

**Figure 11.**
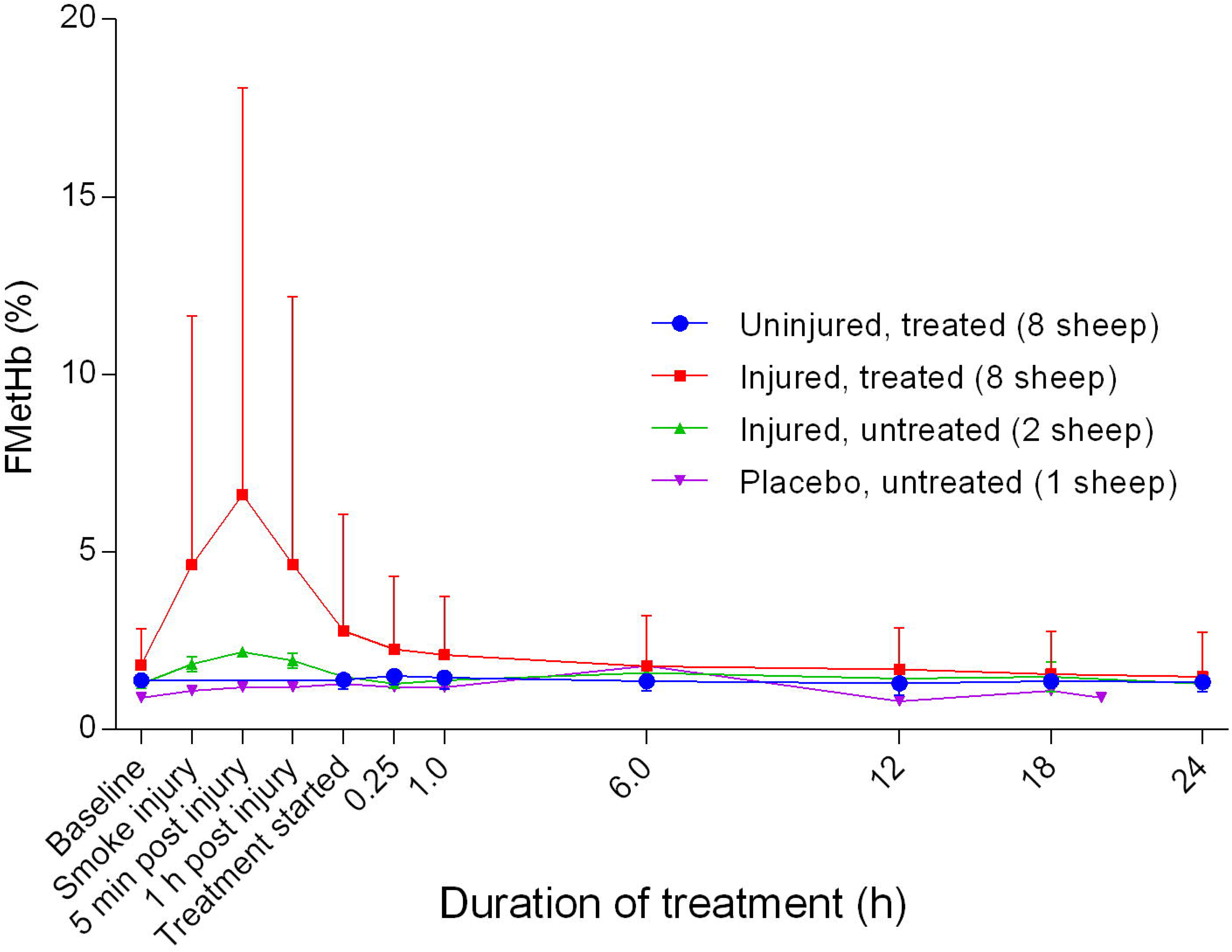
Fraction of methaemoglobin (FMetHb) (Mean ± SD) in smoke and non-smoke injured sheep that received extracorporeal life support as compared to that in untreated controls.

**Figure 12.**
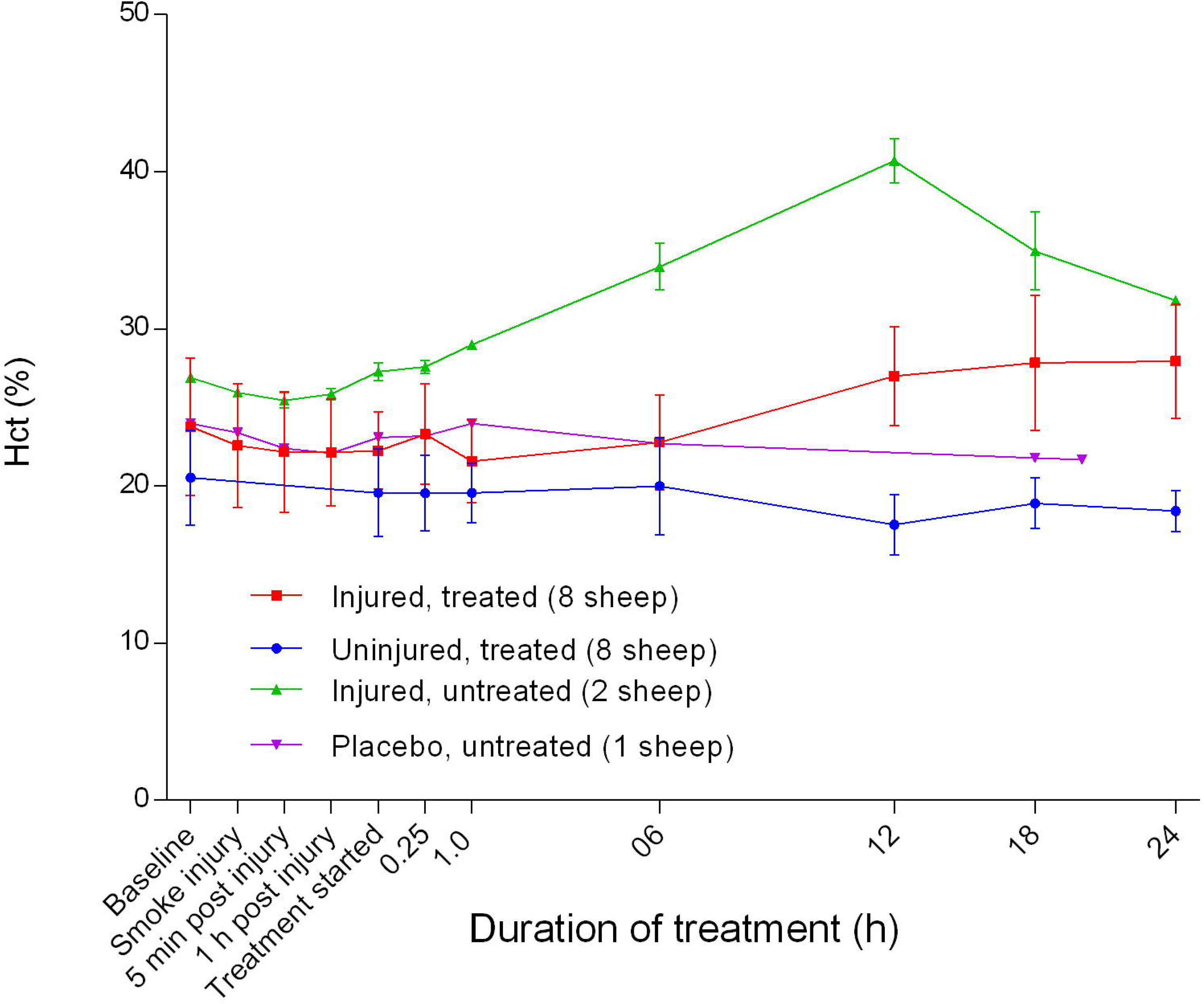
Calculated haematocrit (Hct) (%) (Mean ± SD) in smoke and non-smoke injured sheep that received extracorporeal life support as compared to that in untreated controls.

### Electrolytes

The blood sodium concentration [Na^+^] was relatively stable and there were no significant differences between groups (Figure 13). There was an initial decrease in the blood calcium [Ca^2+^] level, with the lowest point at approximately 1 hour after the treatment was begun, before it levelled out thereafter in all groups (Figure 14). Further, there was a significant difference in [Ca^2+^] between the uninjured/treated and the injured/treated groups (*p* = 0.0001). The placebo/untreated and injured/treated groups maintained the highest and lowest levels of [Ca^2+^], respectively, throughout the experiments. Blood chloride [Cl^-^] levels remained stable compared with baseline levels during the initial stages and then increased gradually thereafter (Figure 15). The blood potassium concentration [K^+^] initially decreased as compared with baseline levels, reaching a minimum concentration 1 hour after the treatment was begun and then gradually increased with a peak at approximately 12 hours after treatment was begun in all experimental groups (Figure 16). Although the injured/untreated and injured/treated sheep had higher [K^+^] than the uninjured sheep, the differences were not significant. Overall, the anion gap decreased gradually, achieving a relatively gentle slope at approximately 6 hours after the treatment was begun and did not change significantly, thereafter (Figure 17). There was a gradual decrease in anion gap from baseline in the course of the experiments and there was no significant difference in anion gap between the uninjured/treated and injured/treated groups.

**Figure 13.**
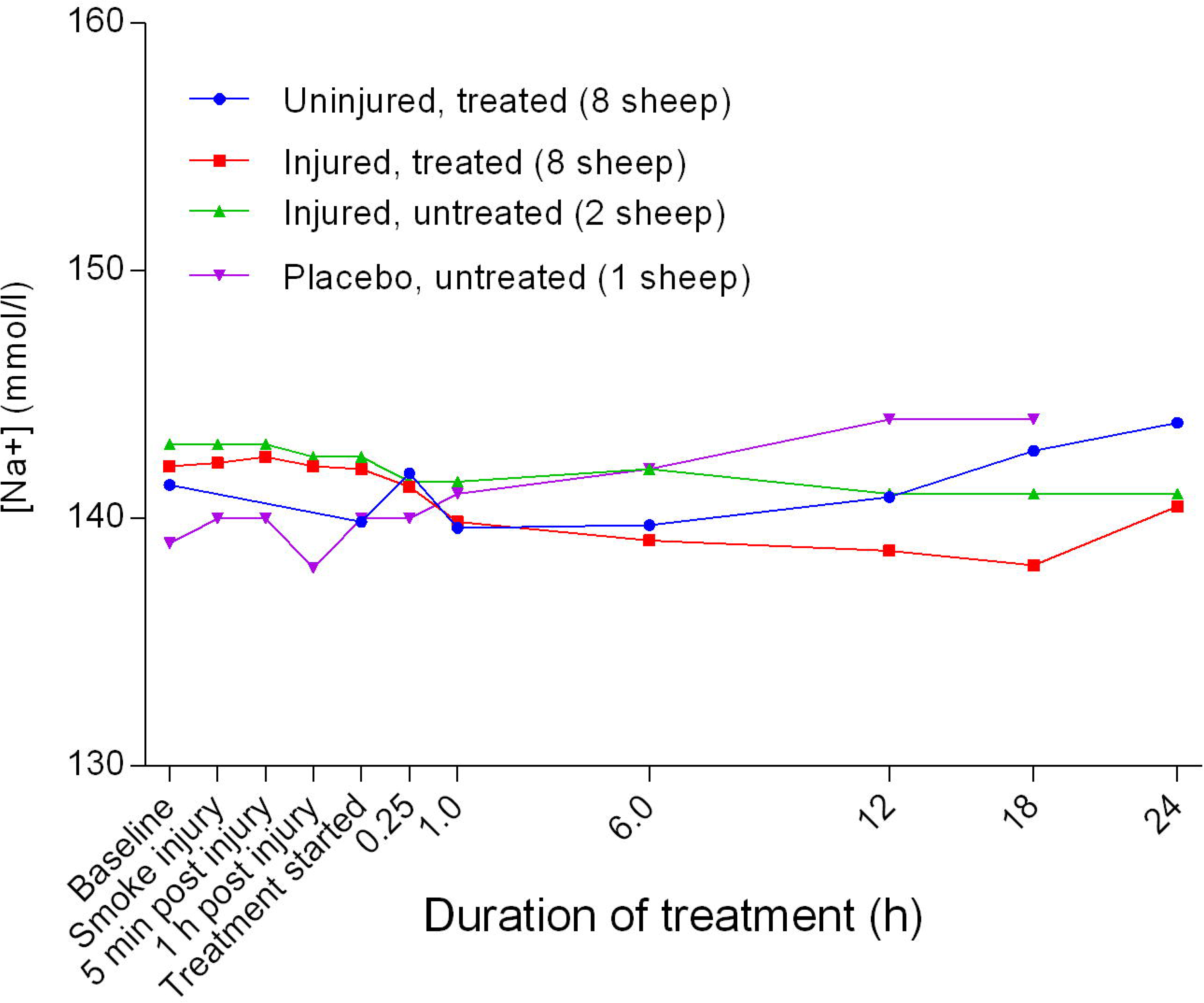
Concentration of sodium ions in blood (Mean ± SD) in smoke and non-smoke injured sheep that received extracorporeal life support as compared to that in untreated controls.

**Figure 14.**
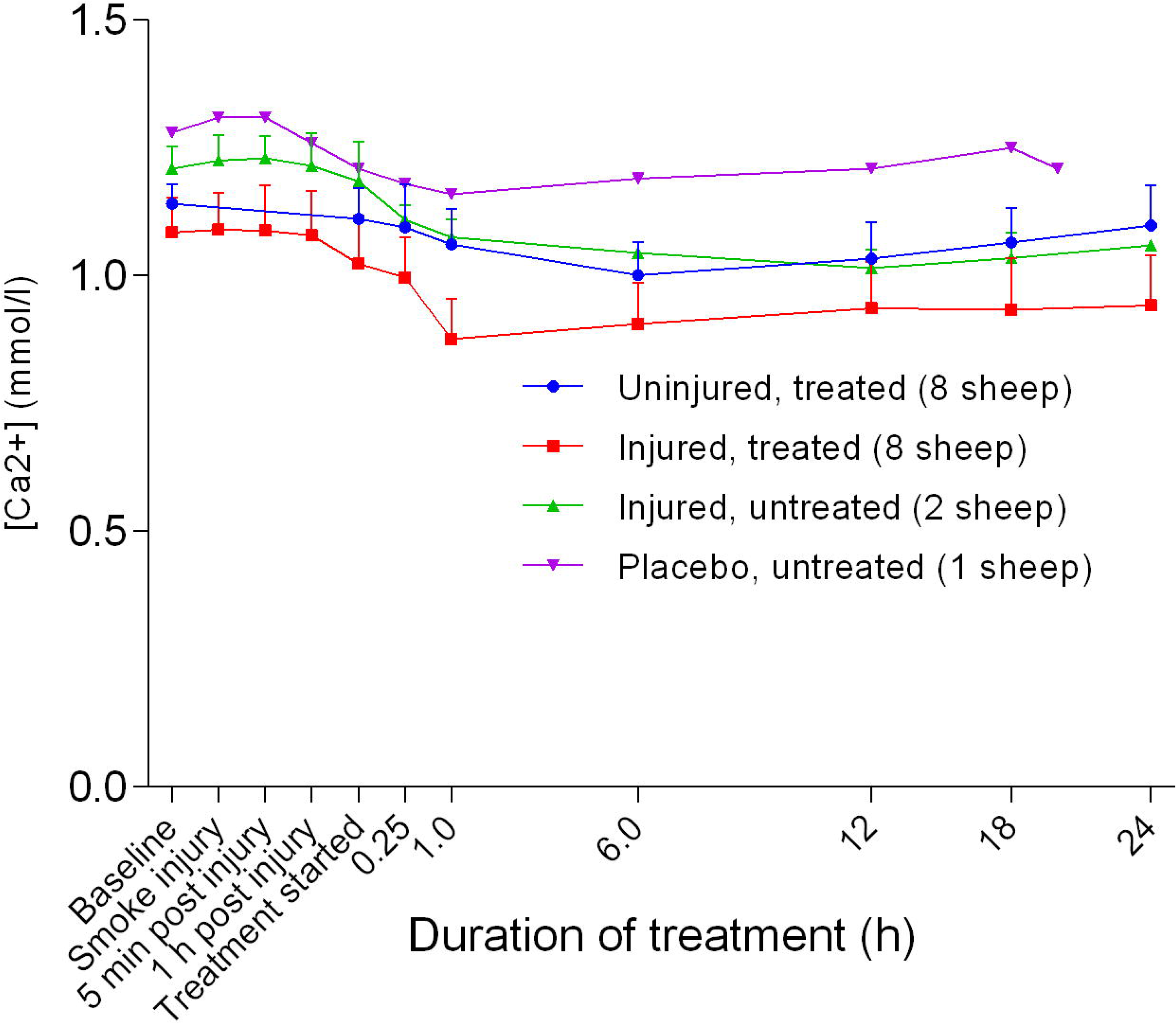
Concentration of calcium ions in blood (Mean ± SD) in smoke and non-smoke injured sheep that received extracorporeal life support as compared to that in untreated controls.

**Figure 15.**
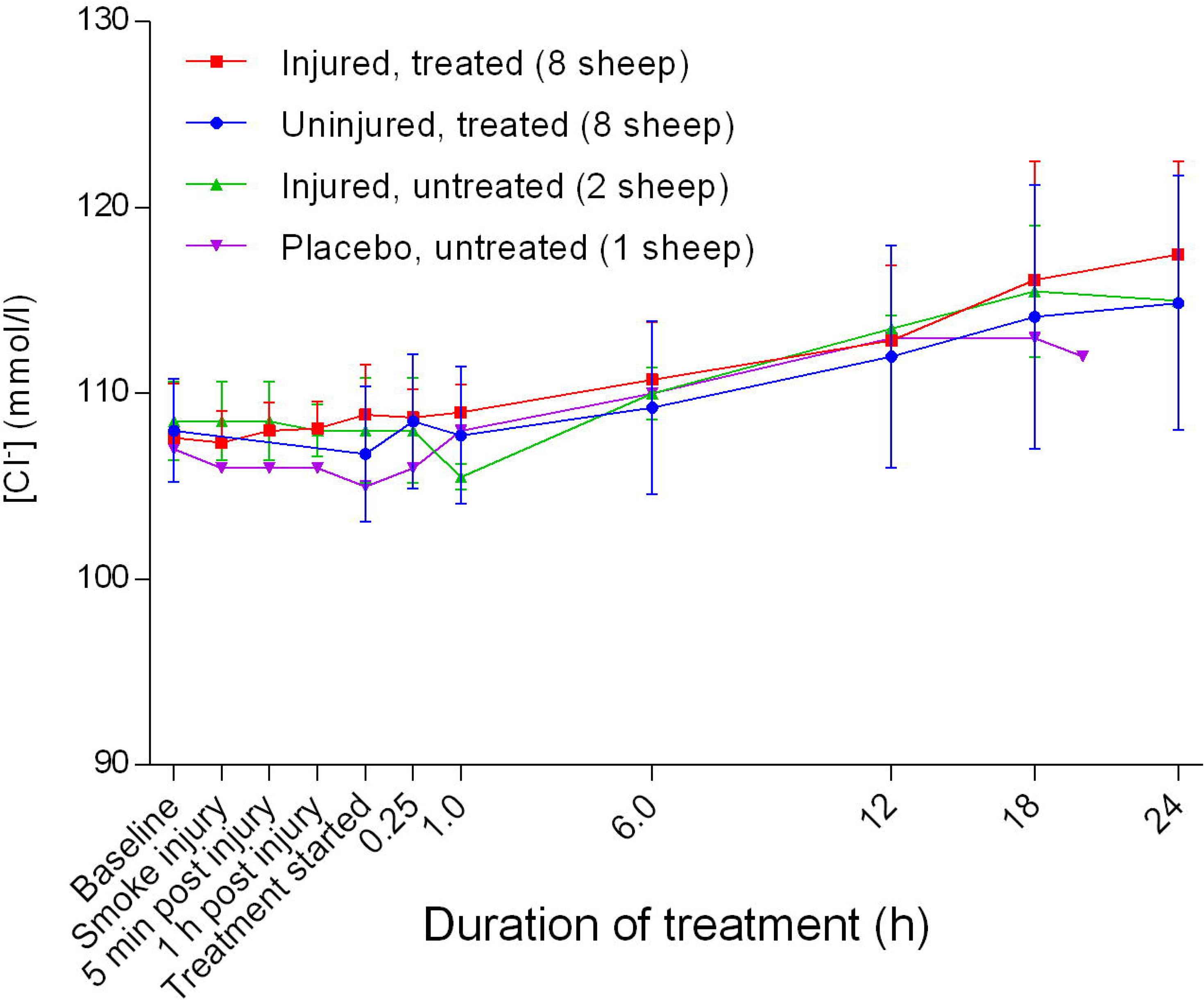
Concentration of chloride ions in blood (Mean ± SD) in smoke and non-smoke injured sheep that received extracorporeal life support as compared to that in untreated controls.

**Figure 16.**
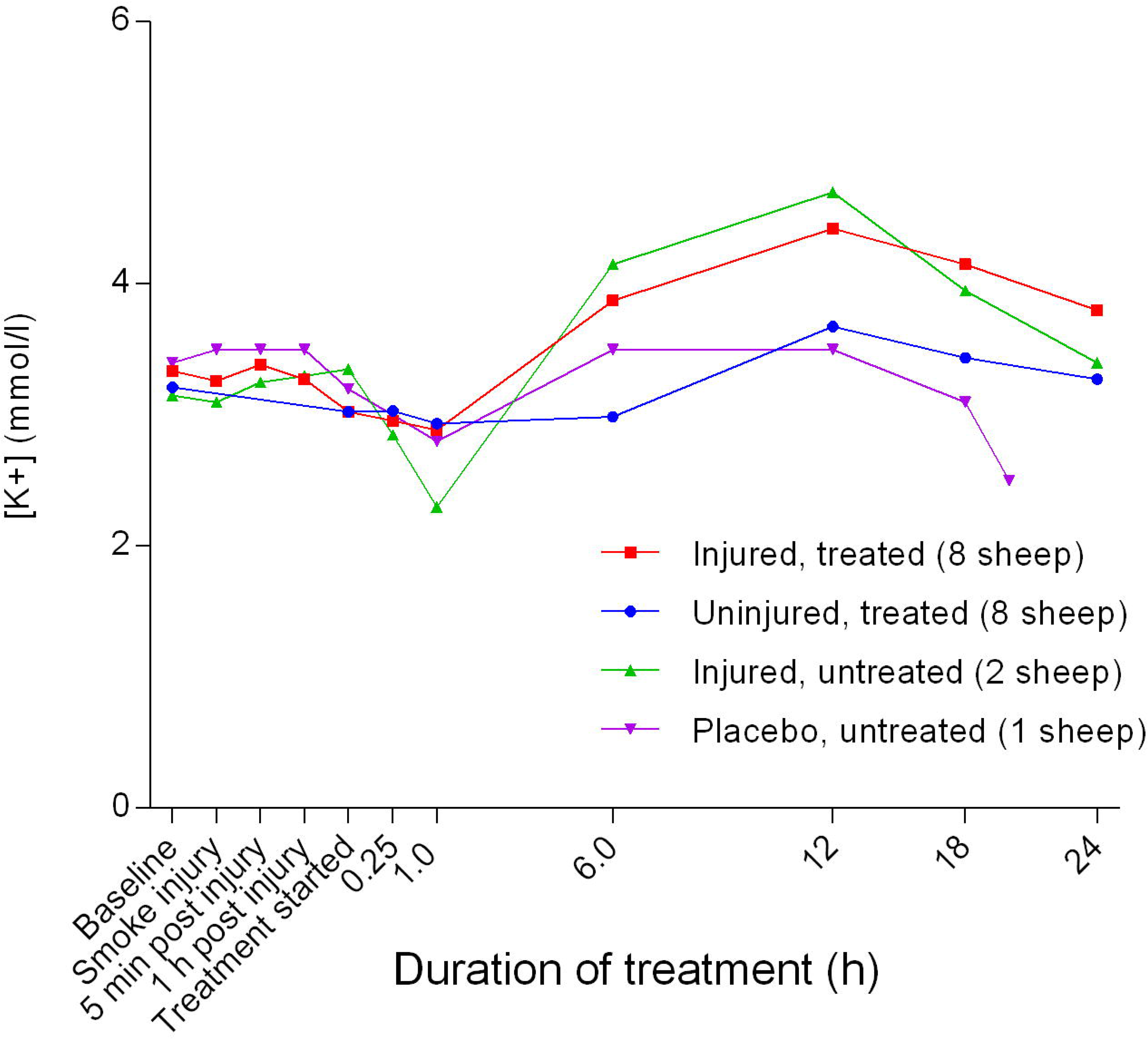
Concentration of potassium ions in blood (Mean ± SD, error bars not shown) in smoke and non-smoke injured sheep that received extracorporeal life support as compared to that in untreated controls.

**Figure 17.**
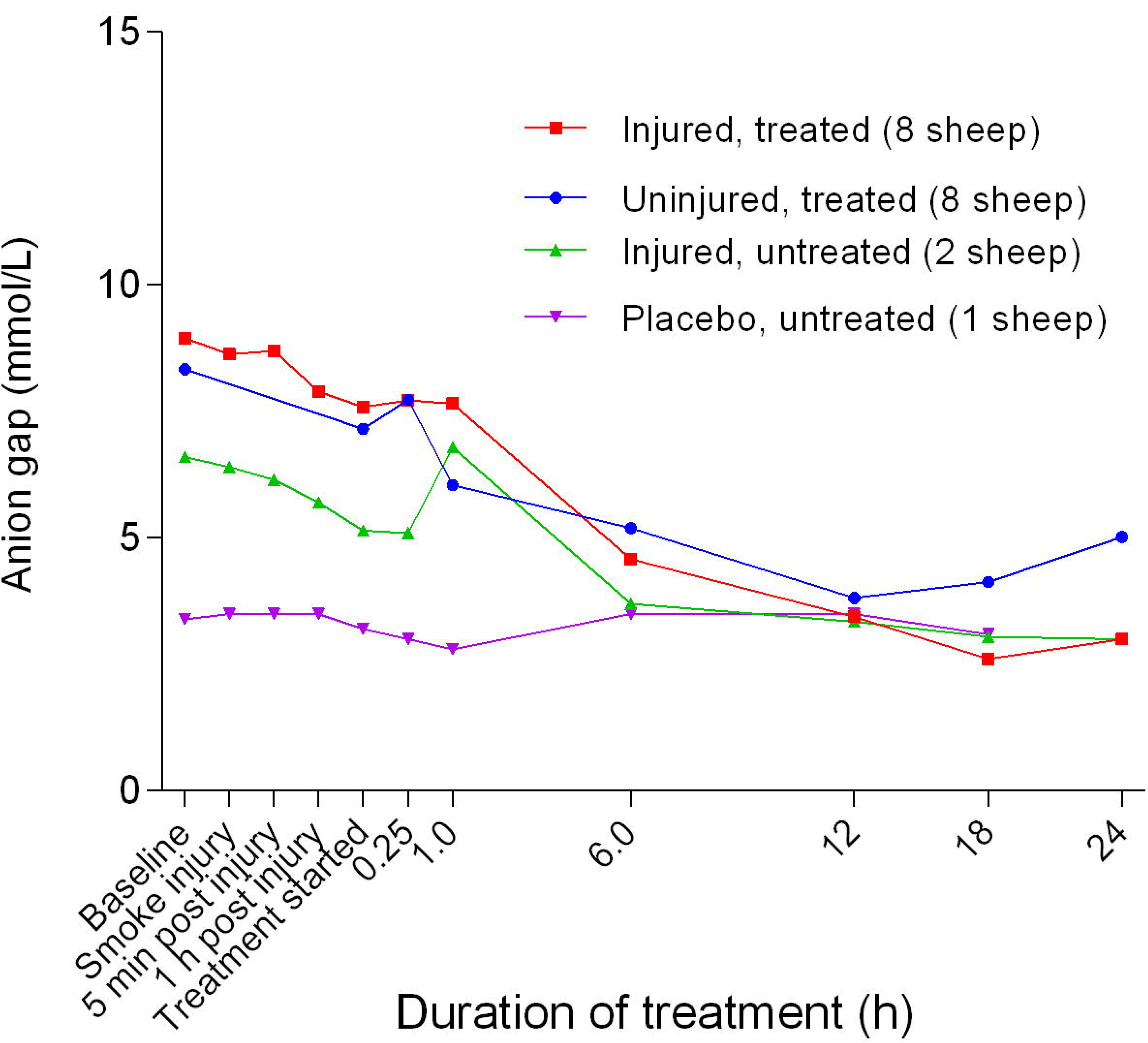
Anion gap (Mean ± SD, error bars not shown) in smoke and non-smoke injured sheep that received extracorporeal life support as compared to that in untreated controls.

**Figure 18.**
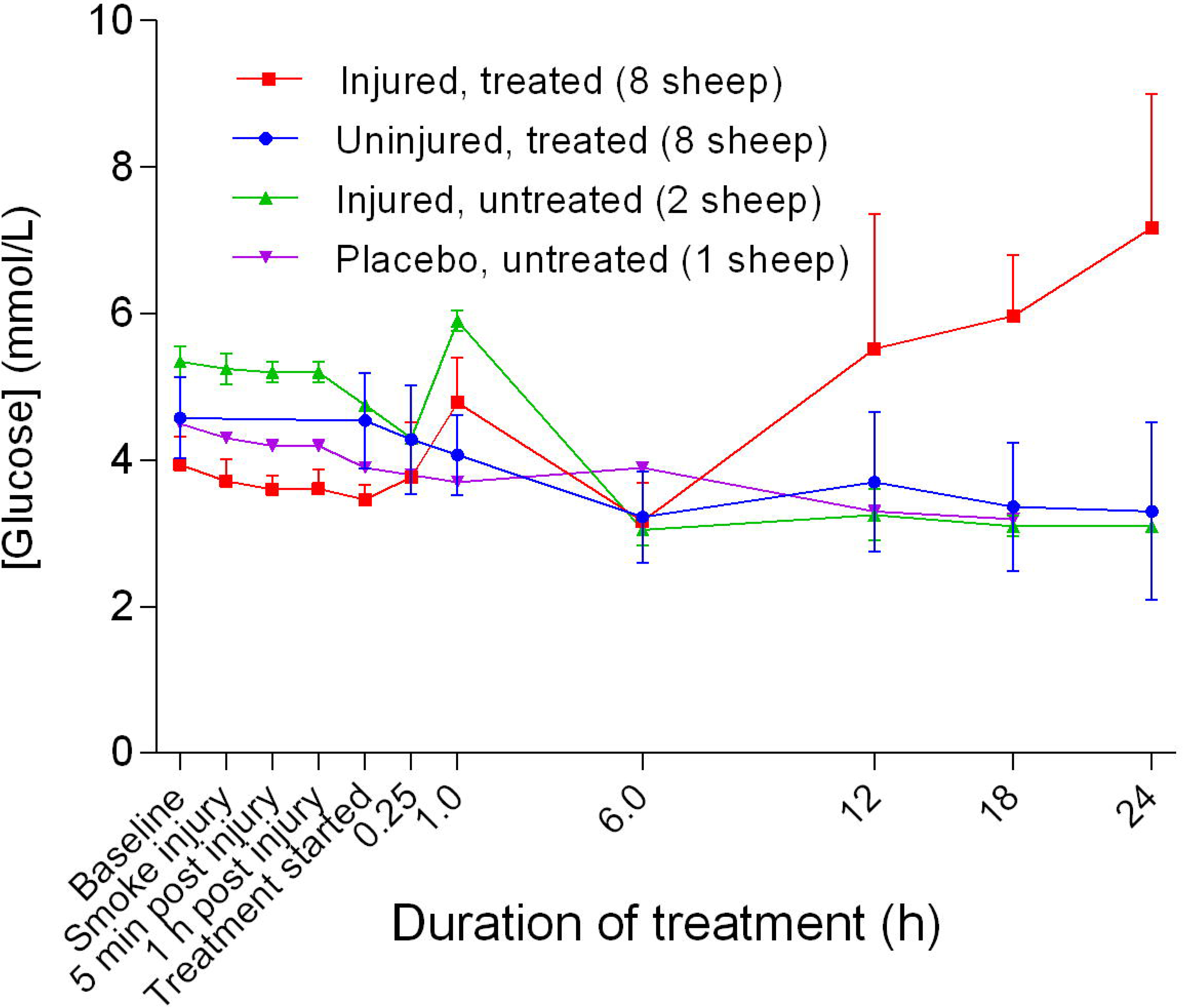
Blood glucose concentration (Mean ± SD) in smoke and non-smoke injured sheep that received extracorporeal life support as compared to that in untreated controls.

**Figure 19.**
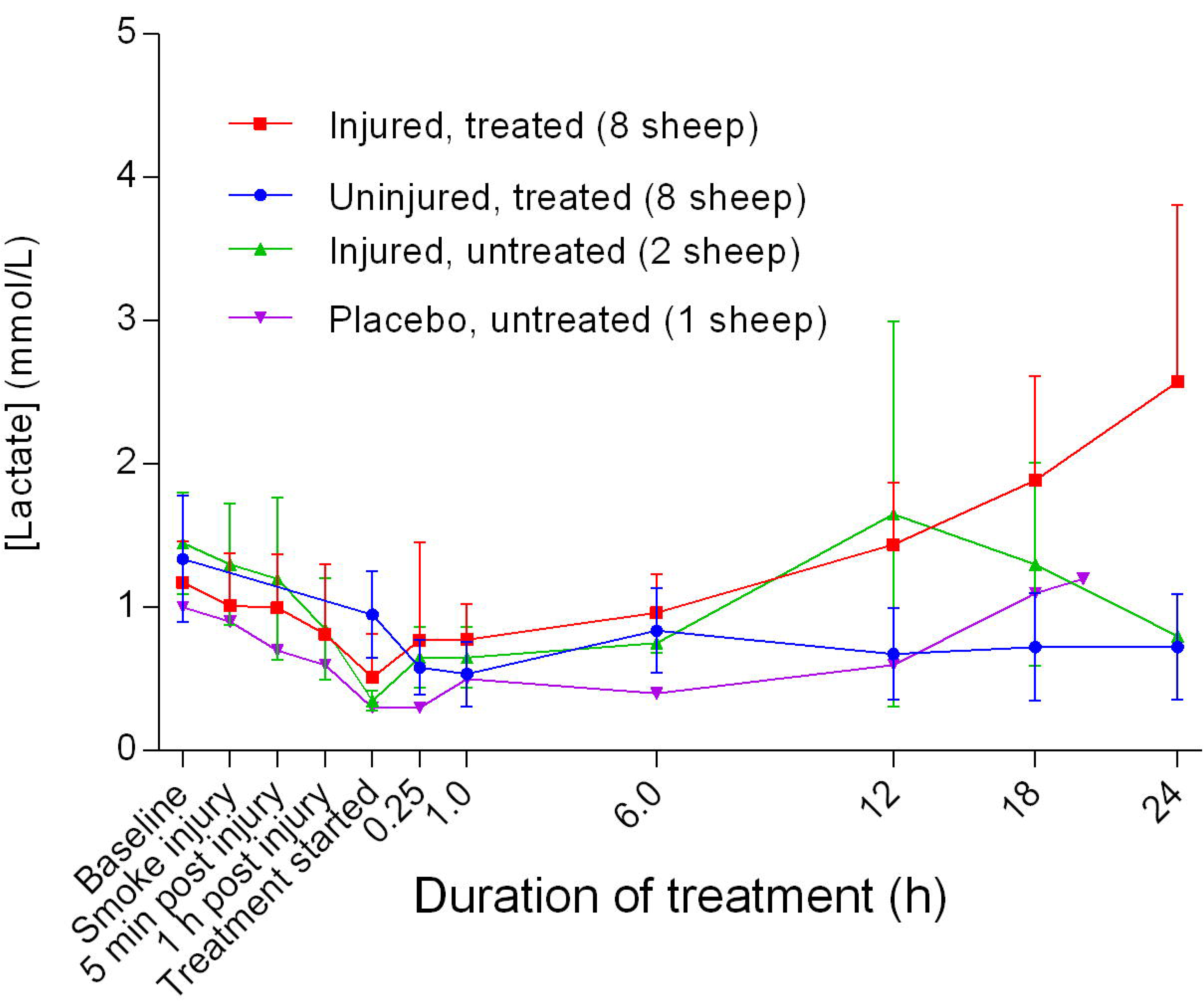
Blood lactate concentration (Mean ± SD) in smoke and non-smoke injured sheep that received extracorporeal life support as compared to that in untreated controls.

### Metabolites

Although there was an increase in blood glucose level [Glu] for the injured/treated sheep after 6 hours of treatment, the change was not significant. There was an initial decrease in lactate levels [Lac] 6 hours after the treatment was begun, followed by a gradual increase for the injured sheep, particularly for the injured/treated group. There was no significant difference in [Lac] between the treated groups.

### Acid-base balance

There was an increase in the blood base levels [Base (ecf)] that peaked 1 hour post-treatment, followed by a gradual decrease in the untreated group. [Base (ecf)] in the treated groups remained at baseline levels to 1 hour after the treatment begun, before decreasing markedly in the injured/treated sheep (Figure 20). There was a significant difference (*p* = 0.0257) in base (ecf) between the uninjured/treated and injured/treated groups. Further, blood bicarbonate concentrations [HCO_3_^-^] increased initially in the untreated groups before decreasing gradually; however, levels remained higher compared with the treated sheep (Figure 21).

**Figure 20.**
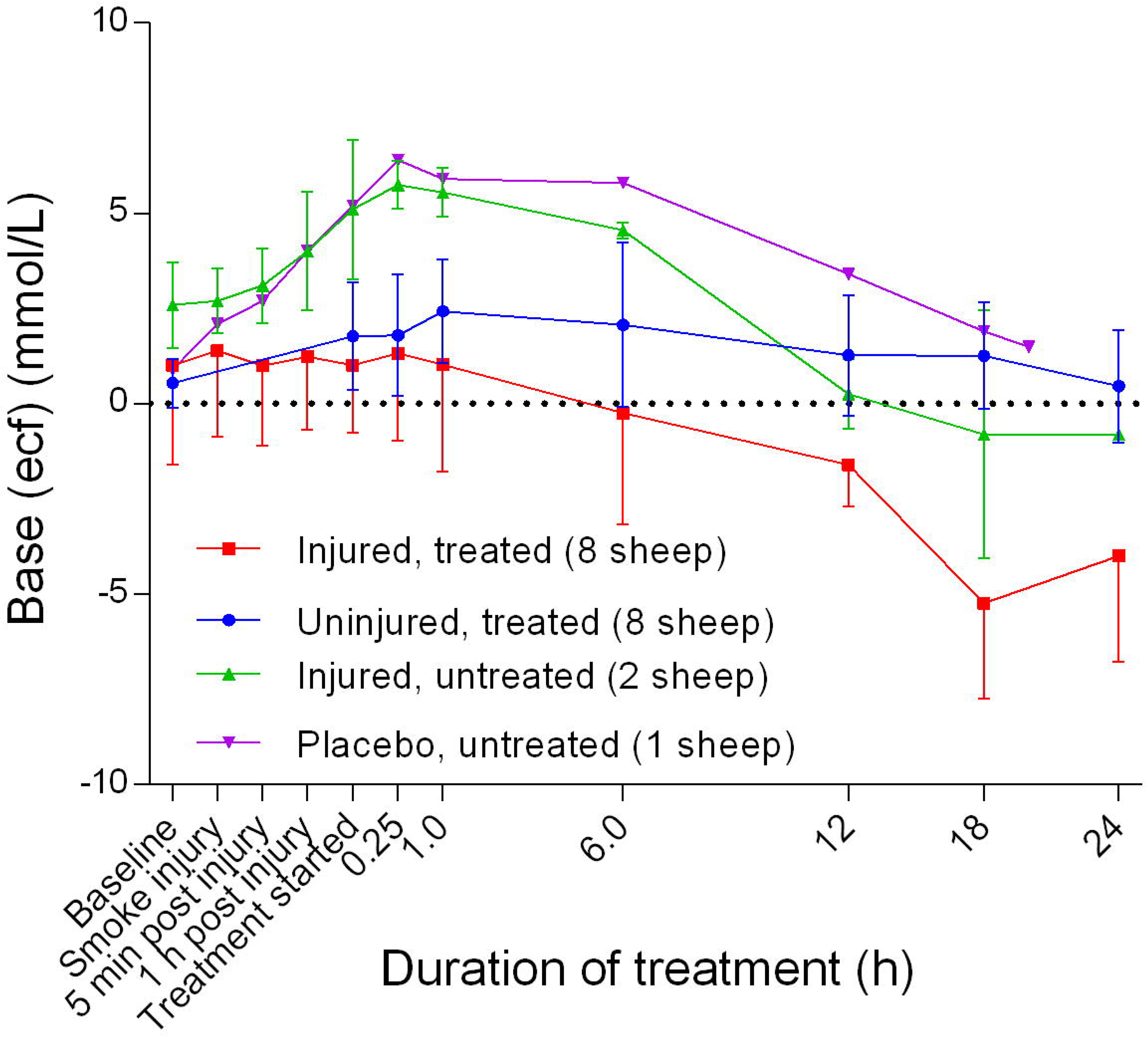
Concentration of base (ecf) in blood (Mean ± SD) for smoke and non-smoke injured sheep receiving extracorporeal life support alongside untreated controls.

**Figure 21.**
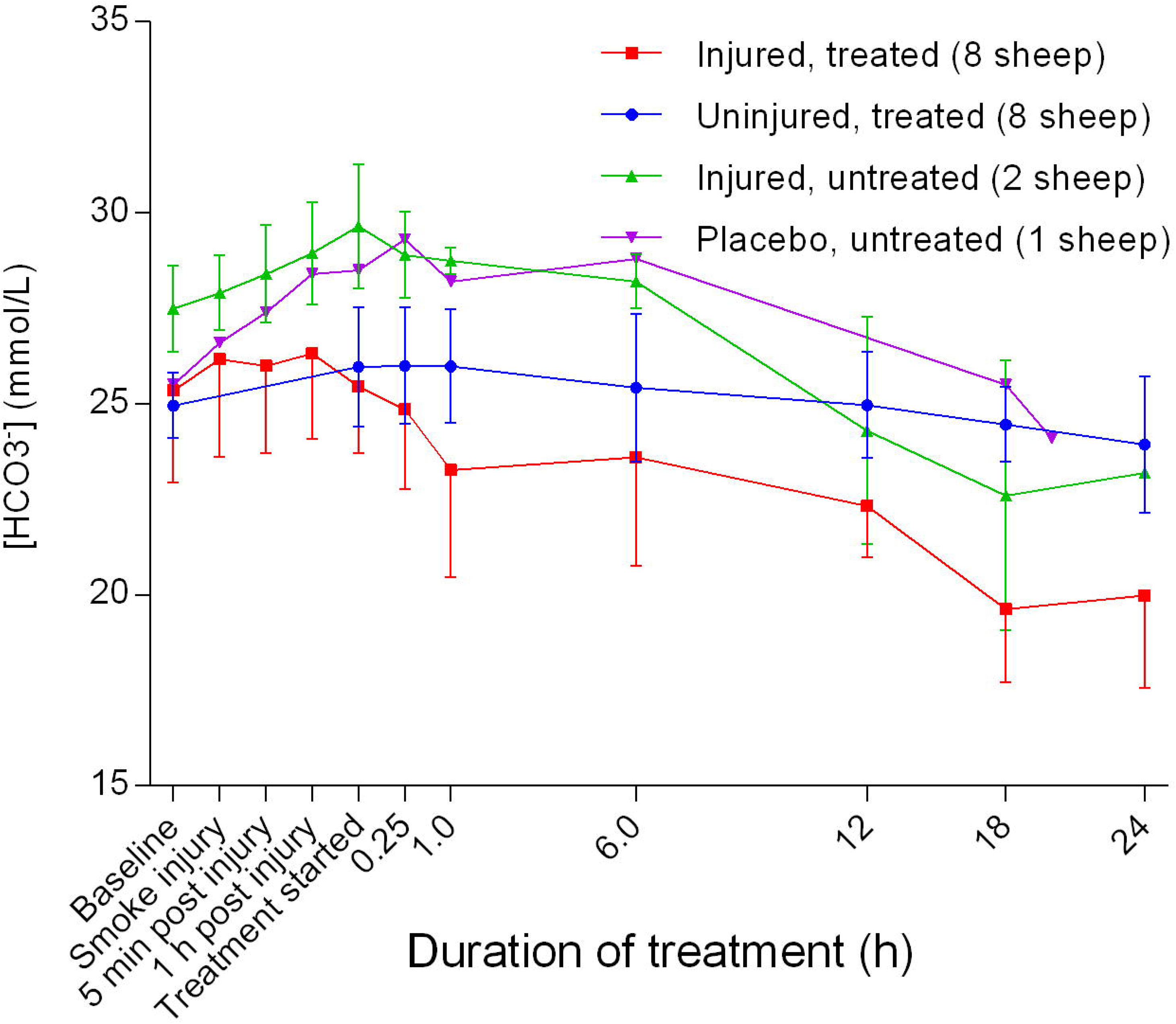
Blood bicarbonate concentration (Mean ± SD) for smoke and non-smoke injured sheep receiving extracorporeal life support alongside untreated controls.

### Haemodynamics

There was a gradual decrease in heart rate (HR) of the sheep during the course of the experiments, with the placebo/untreated groups maintaining a higher HR compared with the injured/untreated, injured/treated, and uninjured/treated groups early in the experiments (Figure 22). There was no significant difference in HR between the uninjured/treated and injured/treated groups. The mean arterial blood pressure (MAP) decreased in the early stages of the experiments before subsequently increasing gradually, peaking at approximately the time that the treatment begun before gradually decreasing again in all but the placebo/untreated sheep (Figure 23). The injured/treated groups had a consistently lower MAP compared with the other groups and there was a significant difference in MAP (*p* = 0.0058) between the uninjured/treated and injured/treated groups. The mean pulmonary artery pressure (MPAP) increased gradually, with the injured/treated group having a consistently higher MPAP (Figure 24). There was no significant difference in MPAP between the uninjured/treated and injured/treated groups. There was an initial, subtle increase in the central venous pressure (CVP) that peaked at approximately 1 hour post-injury followed by a decrease that stabilised at approximately 1 hour after the treatment was begun. Further, CVP levels in the injured/treated and placebo/untreated sheep were consistently higher and lower, respectively, in the course of the experiments.

**Figure 22.**
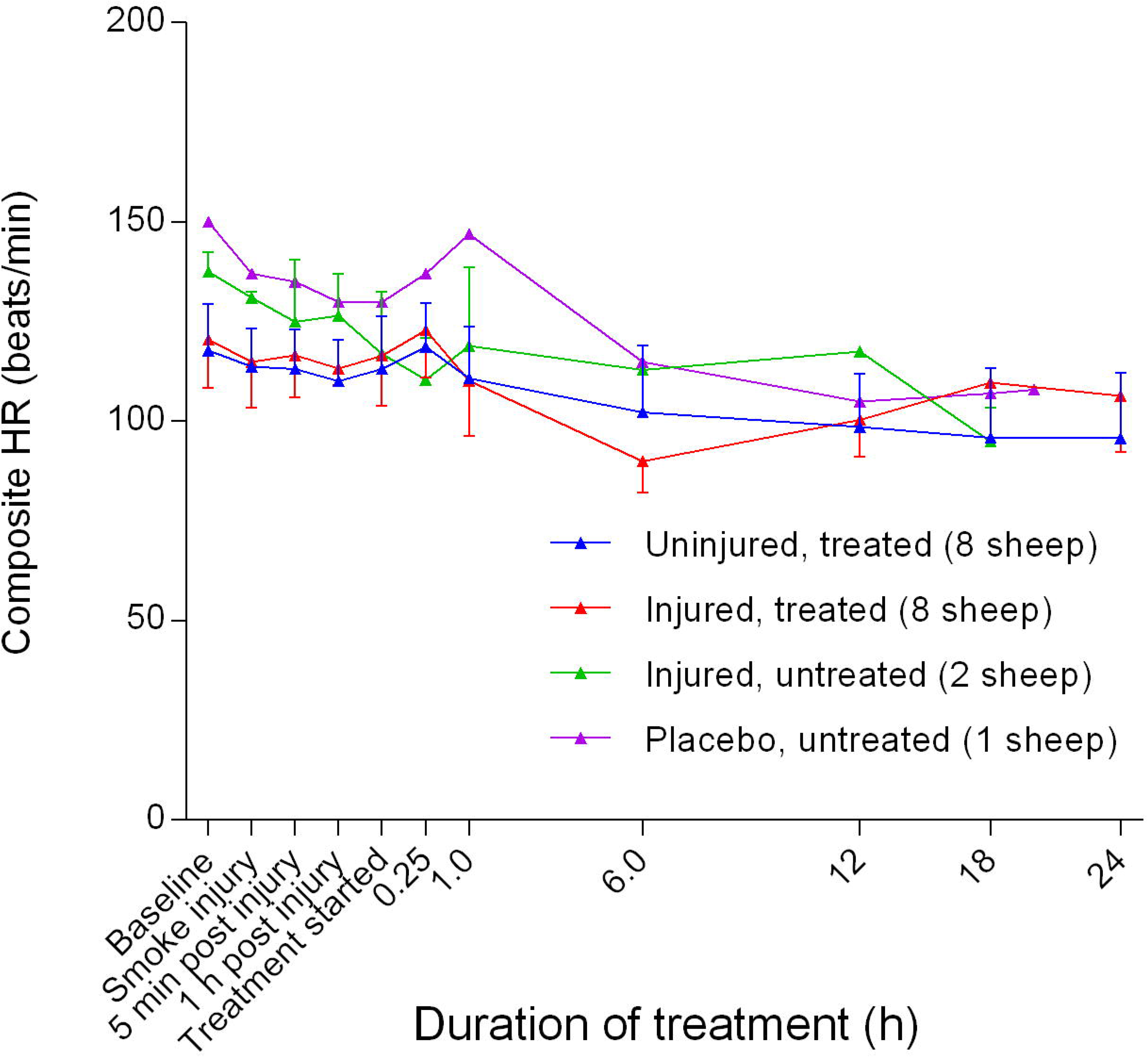
Composite heart rate (HR) (Mean ± SD) in smoke and non-smoke injured sheep that received extracorporeal life support as compared to that in untreated controls.

**Figure 23.**
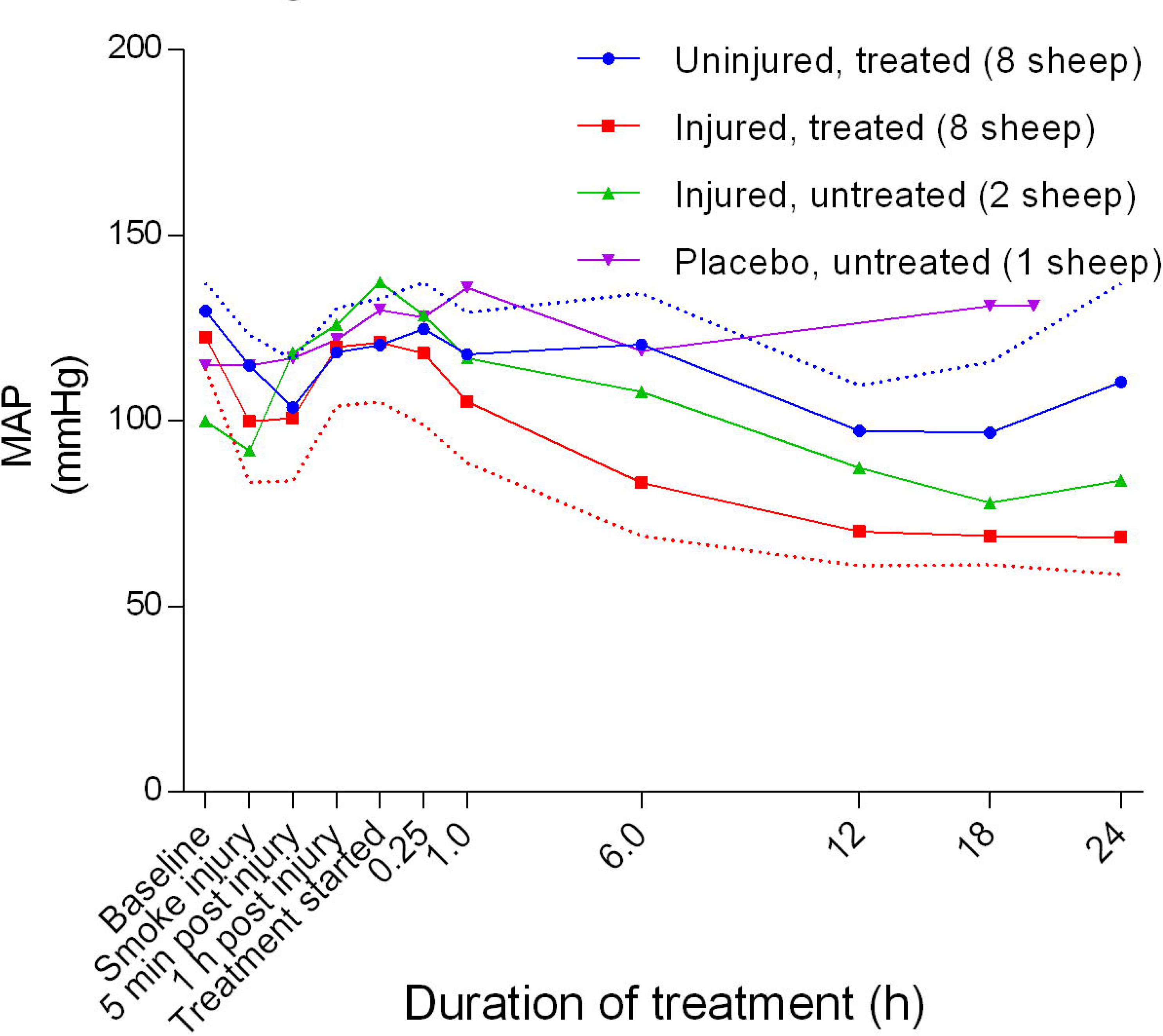
Mean arterial blood pressure (MAP) (Mean ± SD) in smoke and non-smoke injured sheep that received extracorporeal life support as compared to that in untreated controls. Dotted lines represent one-sided margins of error bars.

**Figure 24.**
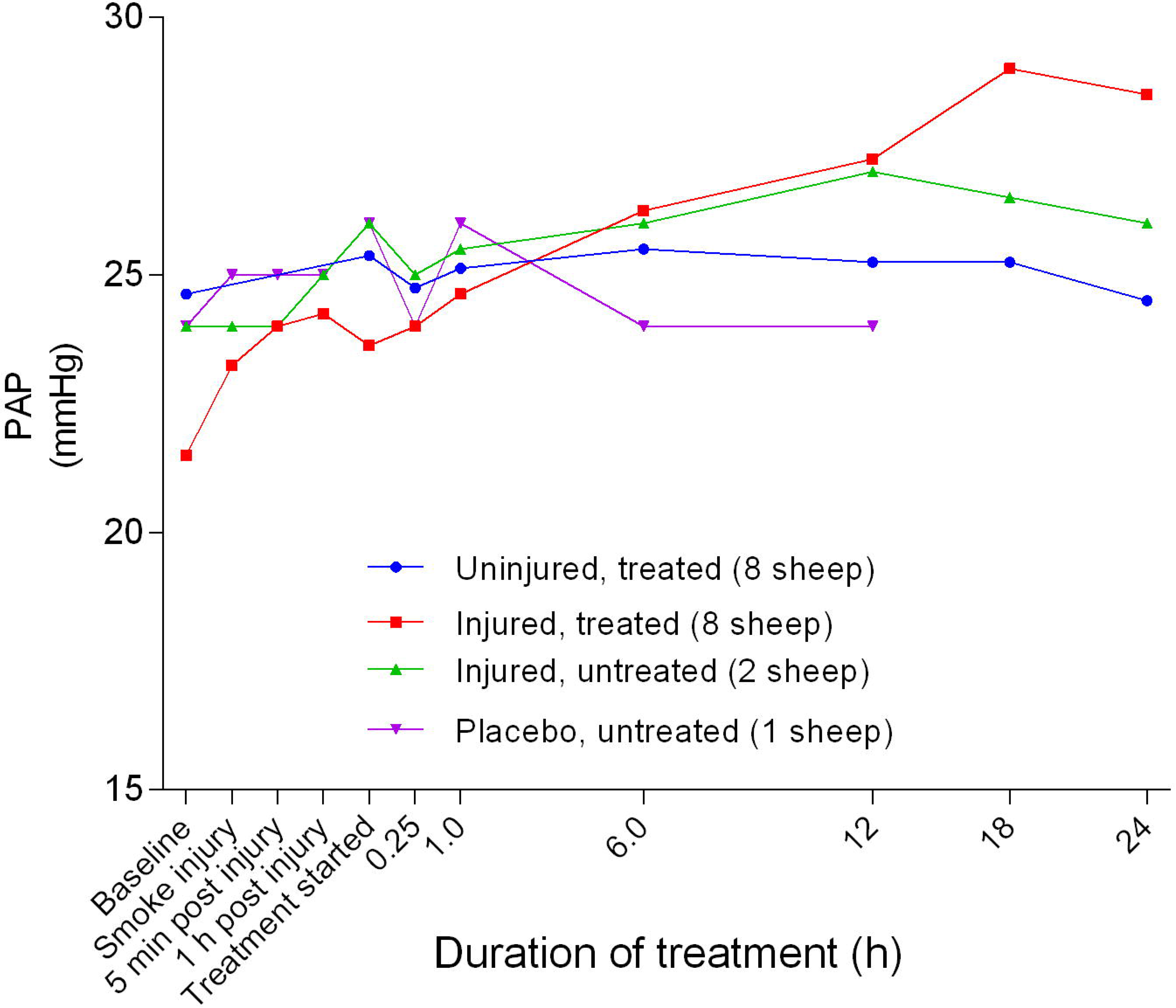
Pulmonary artery pressure (PAP) (Mean ± SD, no error bars shown) in smoke and non-smoke injured sheep that received extracorporeal life support as compared to that in untreated controls.

**Figure 25.**
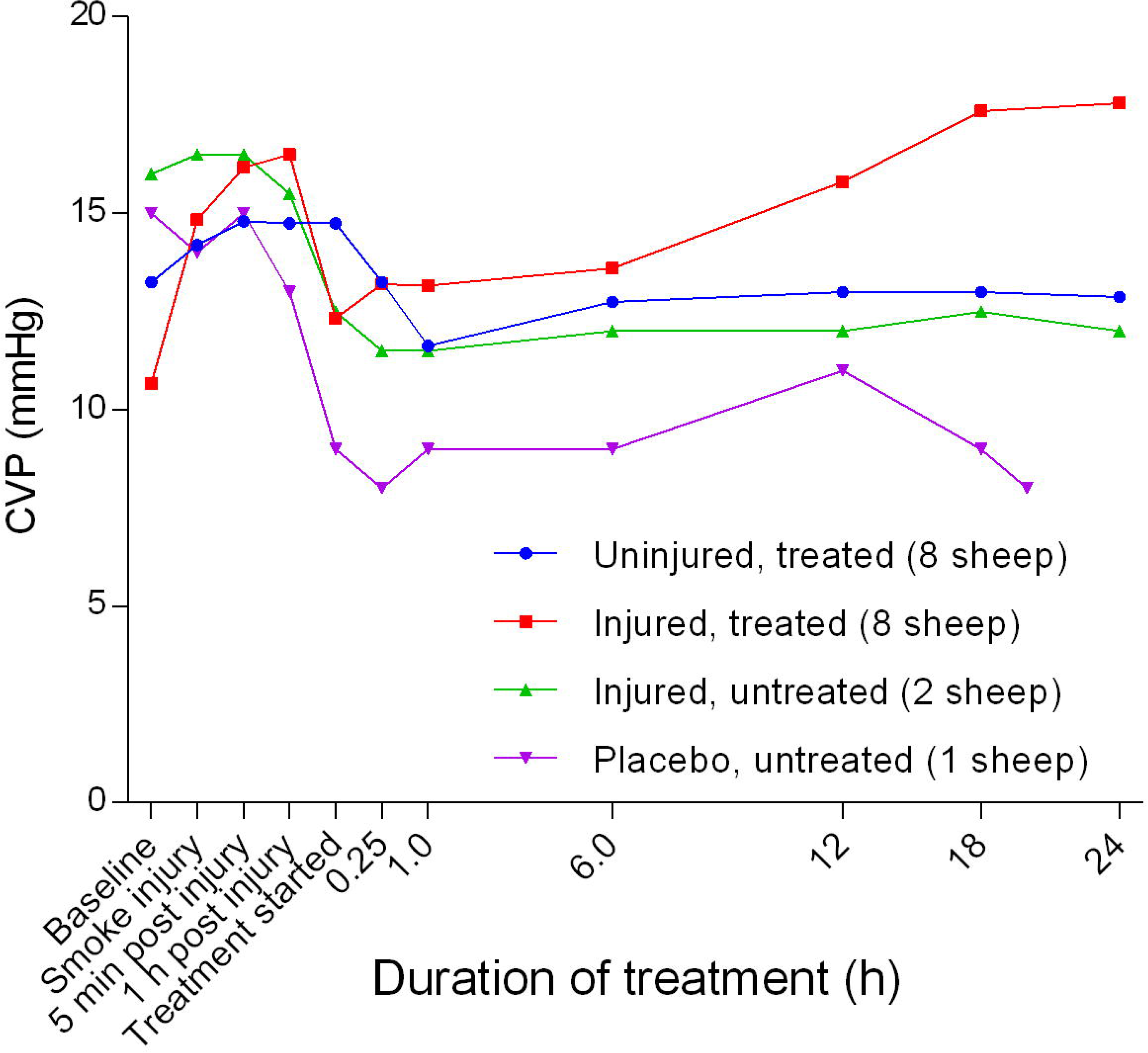
Mean central venous pressure (CVP) (Mean) in smoke and non-smoke injured sheep that received extracorporeal life support as compared to that in untreated controls.

Mixed venous oxygen saturation (SvO_2_) had a lower baseline before eventually rising to a relatively stable and higher level for the treated sheep, and a slightly lower level for the untreated sheep (Figure 26). The injured/untreated sheep maintained a consistently lower SvO_2_ compared with the other groups. Except for the placebo/untreated group, there was a decrease in continuous cardiac output (CCO) from baseline to approximately 1 hour after the treatment was begun (Figure 27). There was a significant difference (*p =* 0.0009) in CCO between the uninjured/treated and injured/treated groups with CCO in the treated groups increasing sharply before plateauing, particularly in the uninjured/treated group. There was also a subsequent gradual decrease in CCO in the injured/treated group. Stroke volume (SV) began to increase 1 hour after the treatment was begun for all groups, except for the injured/untreated group in which levels remained relatively constant (Figure 28). The SV in the injured/treated group began to decrease after 6 hours of treatment, while SV in the uninjured/treated and placebo/untreated sheep increased steadily before decreasing or levelling out after 12 hours or more of treatment. There were no significant differences in SV between the injured/untreated and placebo/untreated. The stroke volume index (SVI) began to increase 1 hour after the treatment was begun for all groups, except for the injured/untreated group, for which SVI remained relatively constant (Figure 29). The SVI in the injured/treated group began to decrease after 6 hours of treatment while SVI in the uninjured/treated and placebo/untreated groups increased before subsequently decreasing or levelling out after 12 hours or more of treatment. There were no significant differences in SVI between groups. While the cardiac index (CI) of the uninjured/treated and placebo/untreated groups remained relatively close to baseline levels (Figure 30), the CI of the injured/treated and injured/untreated groups declined gradually over the course of the experiments.

**Figure 26.**
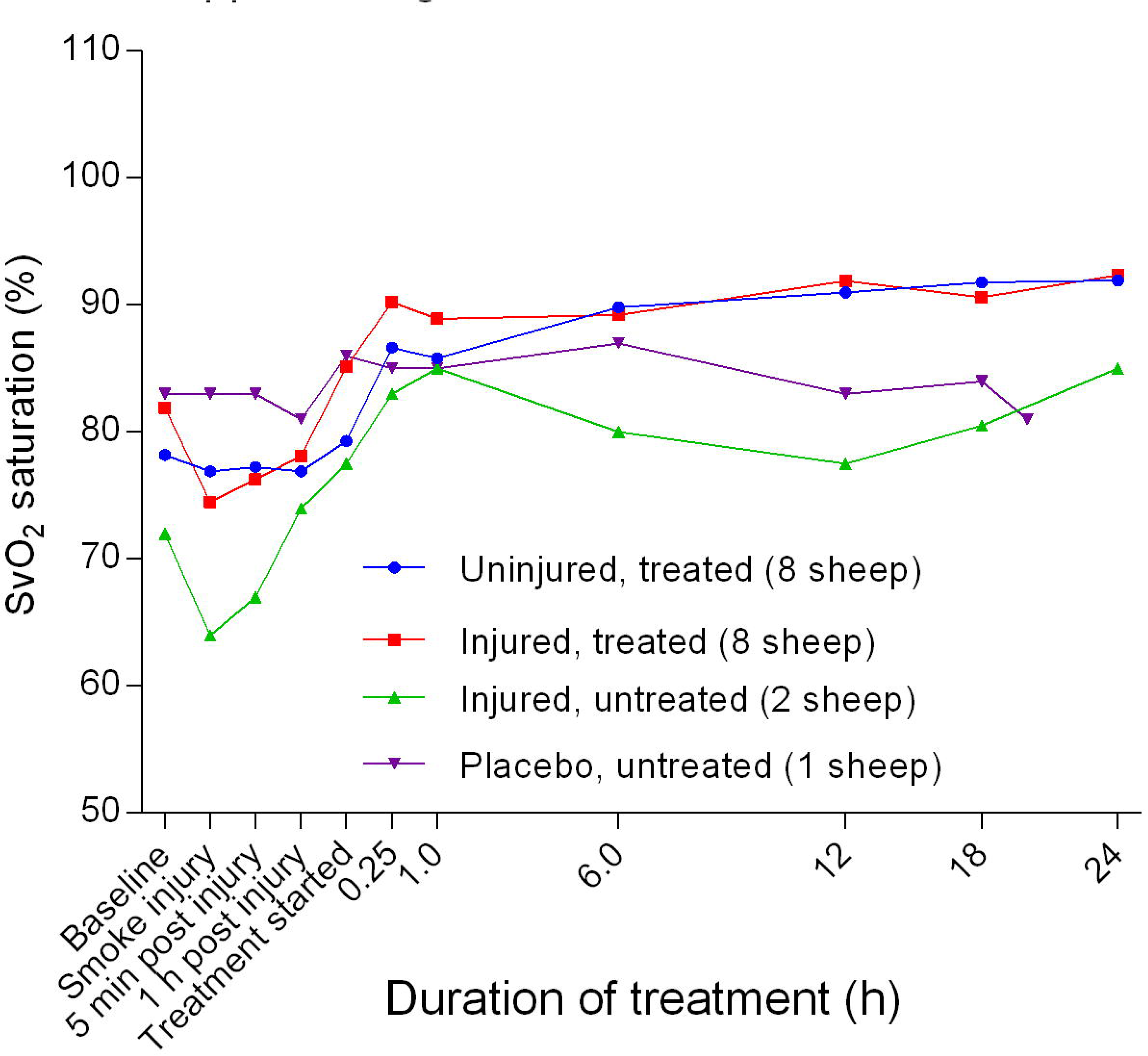
Mixed venous oxygen saturation (SvO_2_) (Mean ± SD, error bars not shown) in smoke and non-smoke injured sheep that received extracorporeal life support as compared to that in untreated controls.

**Figure 27.**
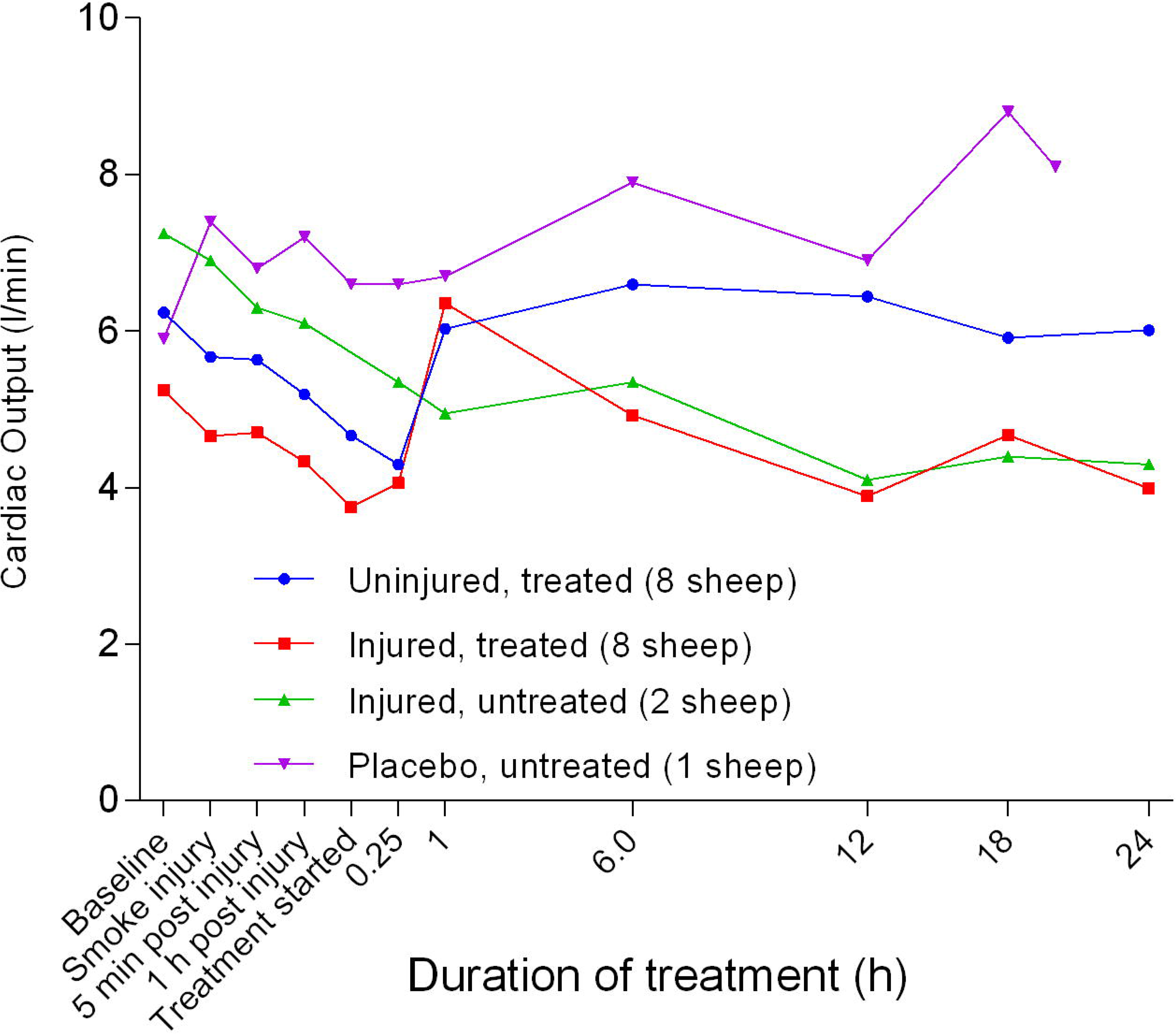
Continuous cardiac output (CCO) (Mean ± SD, error bars not shown) in smoke and non-smoke injured sheep that received extracorporeal life support as compared to that in untreated controls.

**Figure 28.**
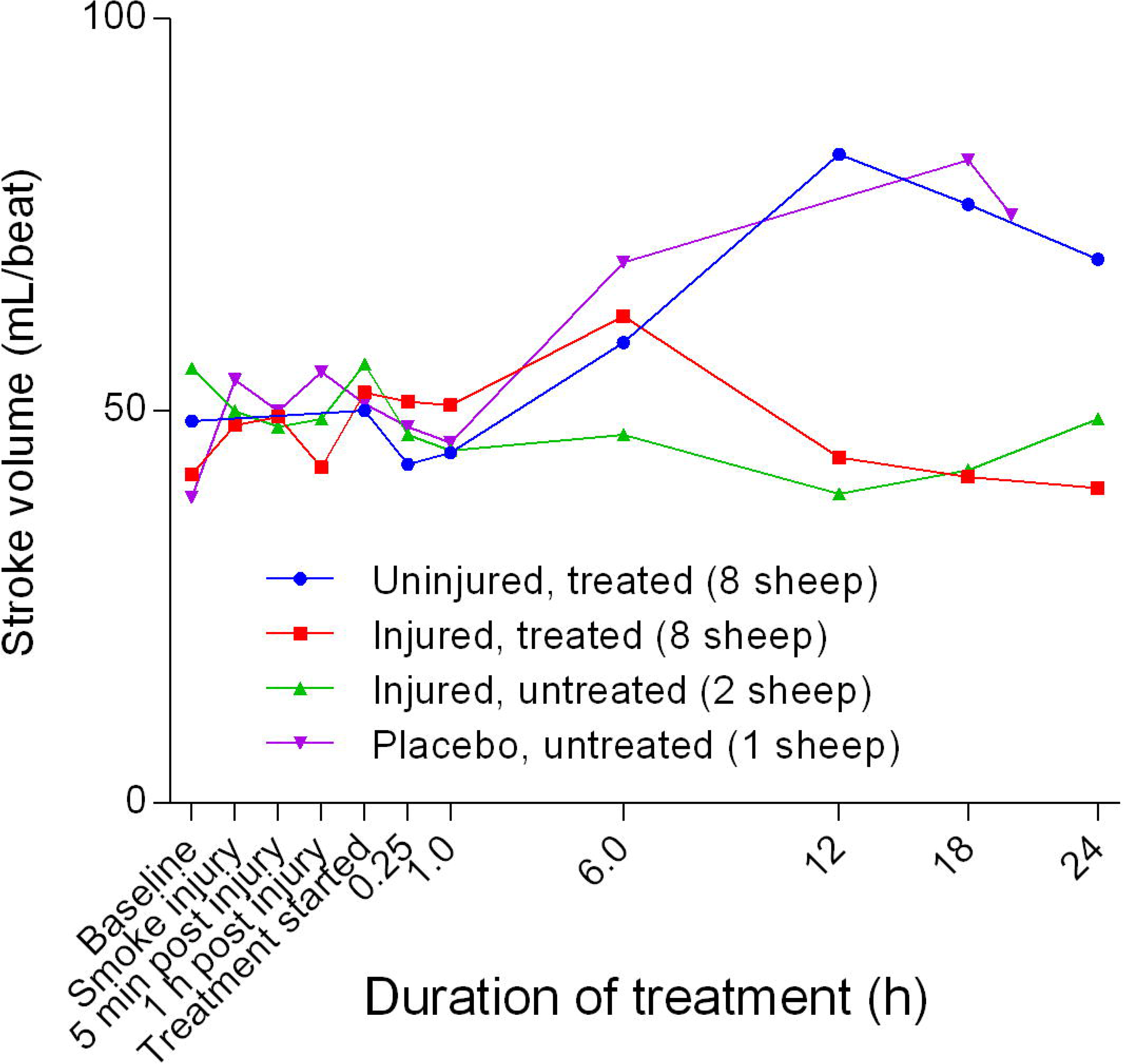
Stroke volume (Mean ± SD error bars not shown) in smoke and non-smoke injured sheep that received extracorporeal life support as compared to that in untreated controls.

**Figure 29.**
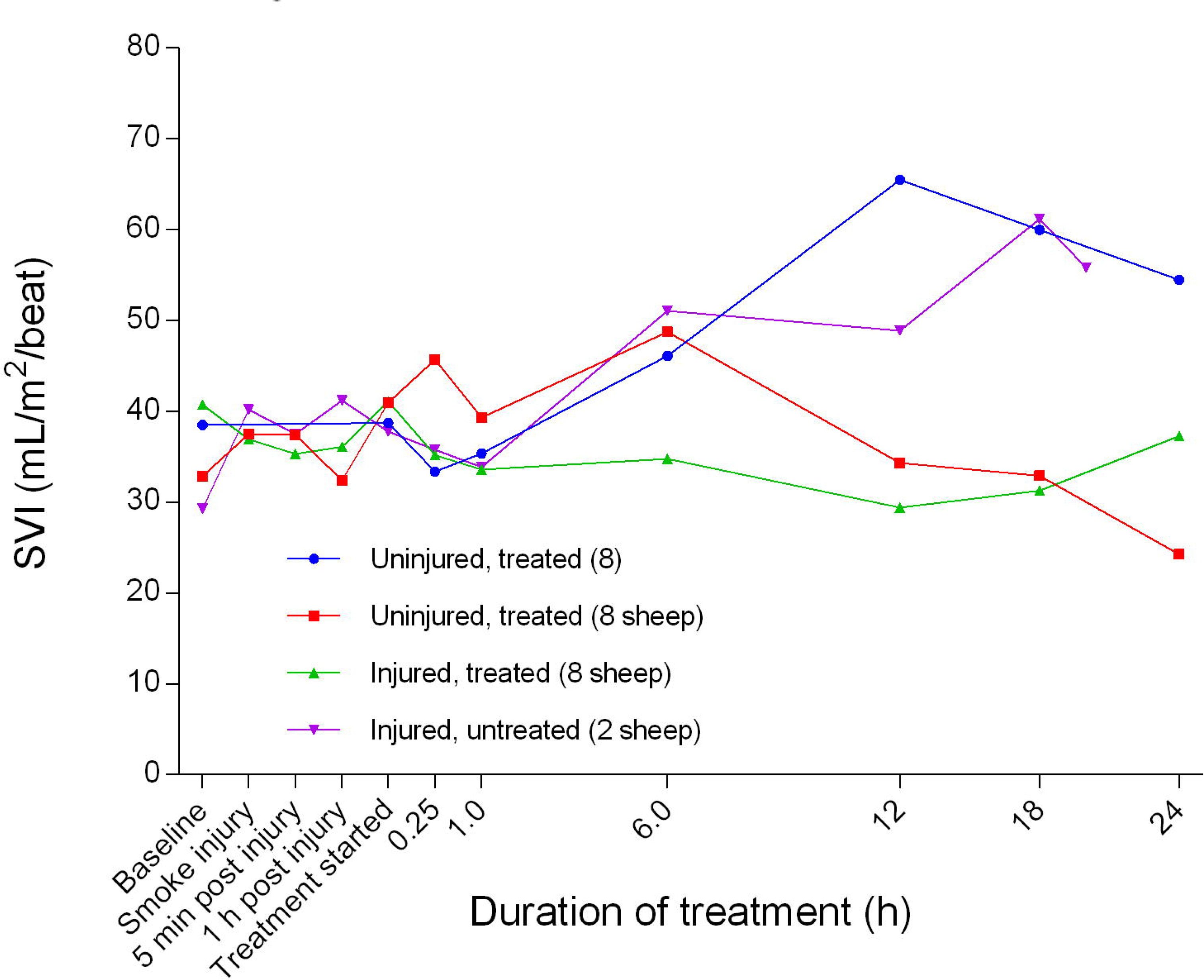
Stroke volume index (SVI) (Mean ± SD, error bars not shown) in smoke and non-smoke injured sheep that received extracorporeal life support as compared to that in untreated controls.

**Figure 30.**
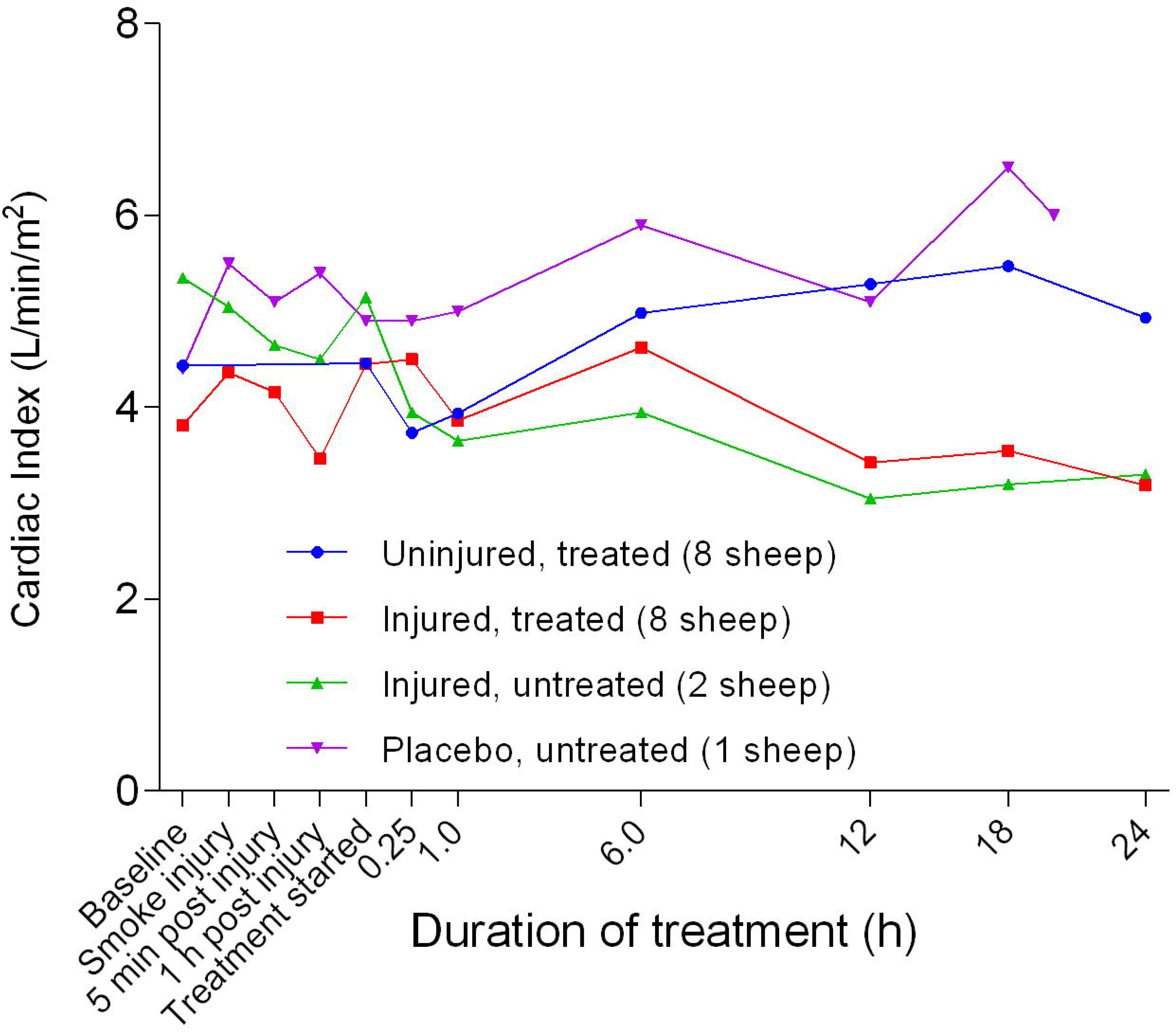
Cardiac index (Mean ± SD error bars not shown) in smoke and non-smoke injured sheep that received extracorporeal life support as compared to that in untreated controls.

After an initial increase in systemic vascular resistance index (SVRI) approximately 1 hour after treatment (Figure 31), SVRI began to decrease in all experimental groups before plateauing 12 hours after treatment, followed by a gentle increasing trend until the end of the experiments. The SVRI in the injured/treated group was consistently below that of the other groups during treatment while that of the injured/untreated group was correspondingly higher. There was no significant difference in SVRI between the groups. The pulmonary vascular resistance index (PVRI) remained close to baseline levels for all of the groups after 1 hour of treatment while that of the injured groups progressively increased and that of the uninjured groups remained lower with a subtle decrease after 6 hours of treatment (Figure 32). The PVRI in the placebo/untreated sheep remained close to baseline levels and the lowest throughout the course of the experiment. After a small peak attained at the beginning of the treatment, the right ventricular stroke work index (RVSWI) in the uninjured sheep gradually increased while that of the injured sheep decreased (Figure 33). There was a significant difference (*p* = 0.0196) in the RVSWI gap between the uninjured/treated and injured/treated groups. RVSWI in the placebo/untreated group remained high, while that of the injured/treated group was consistently the lowest. The left ventricular stroke work index (LVSWI) gradually increased in the uninjured/treated and placebo/untreated groups and plateaued 12 and 18 hours after treatment was begun, respectively, while LVSWI in the injured/untreated and injured/treated groups of sheep decreased and plateaued at 12 hours after treatment was begun and trended upward after 18 hours of treatment (Figure 34). LVSWI in the placebo/untreated group remained consistently higher than in the other groups while that of the injured/treated group was consistently the lowest.

**Figure 31.**
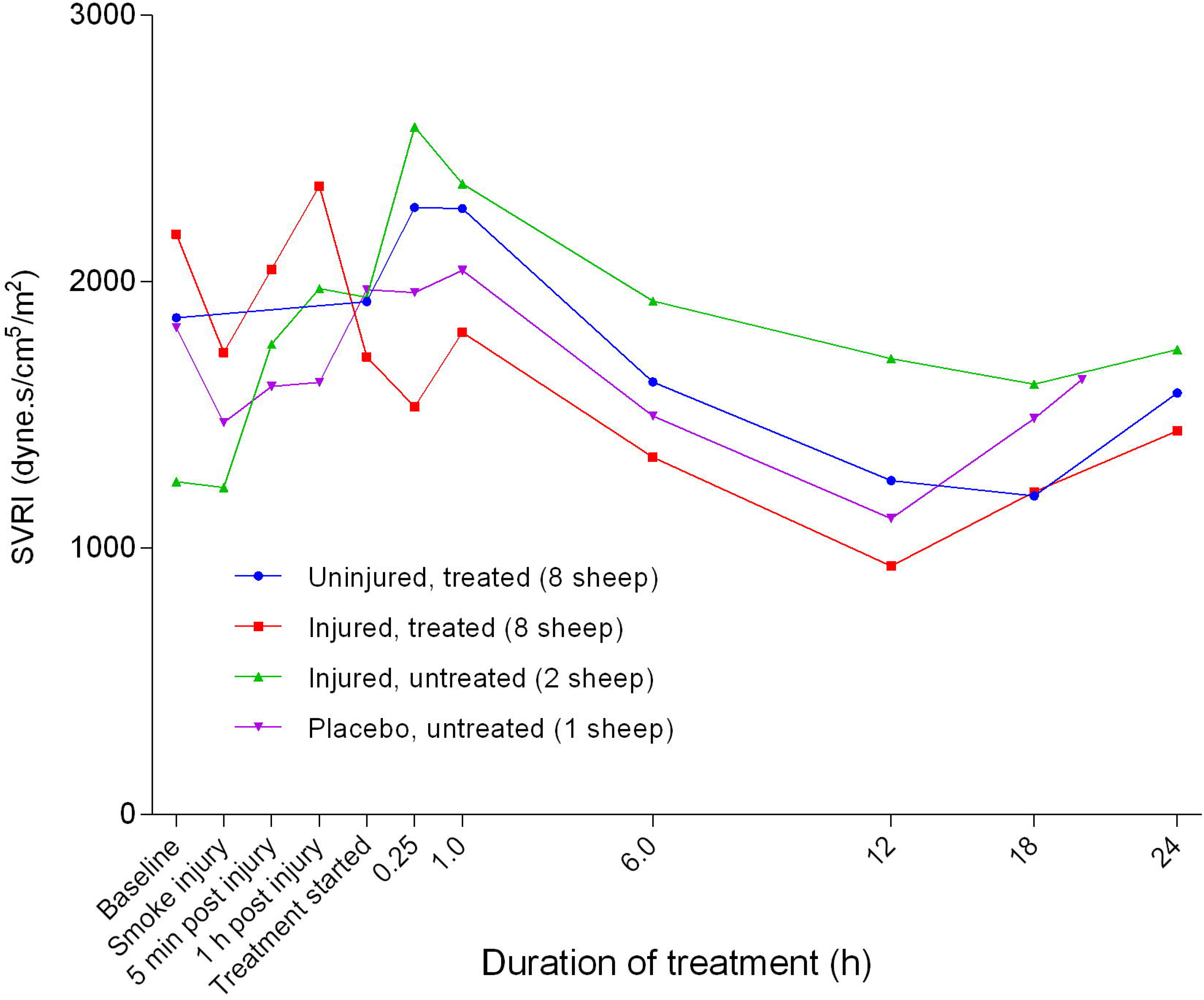
Systemic vascular resistance index (SVRI) (Mean ± SD error bars not shown) in smoke and non-smoke injured sheep that received extracorporeal life support as compared to that in untreated controls.

**Figure 32.**
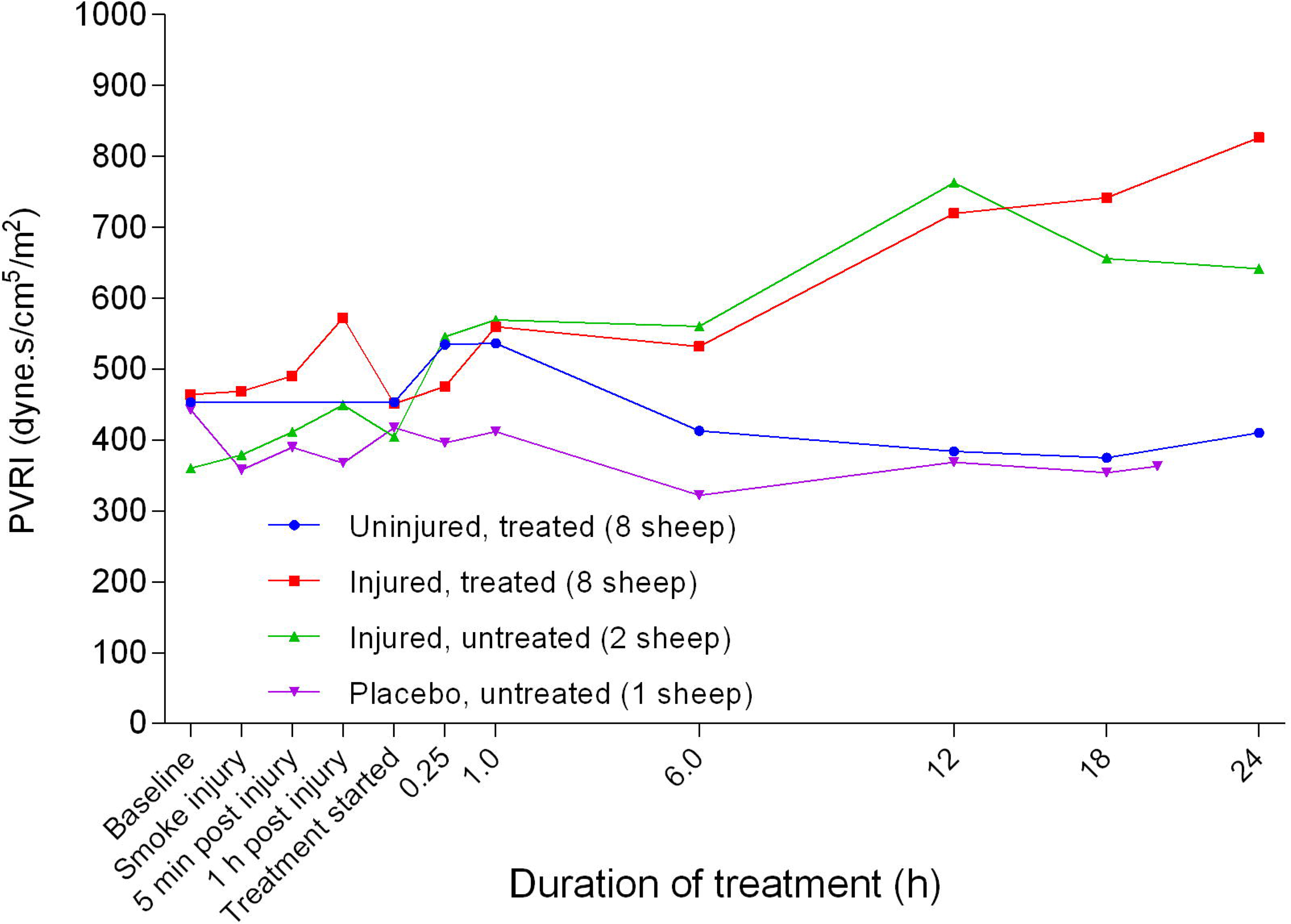
Pulmonary vascular resistance index (PVRI) (Mean ± SD, error bars not shown) in smoke and non-smoke injured sheep that received extracorporeal life support as compared to that in controls.

**Figure 33.**
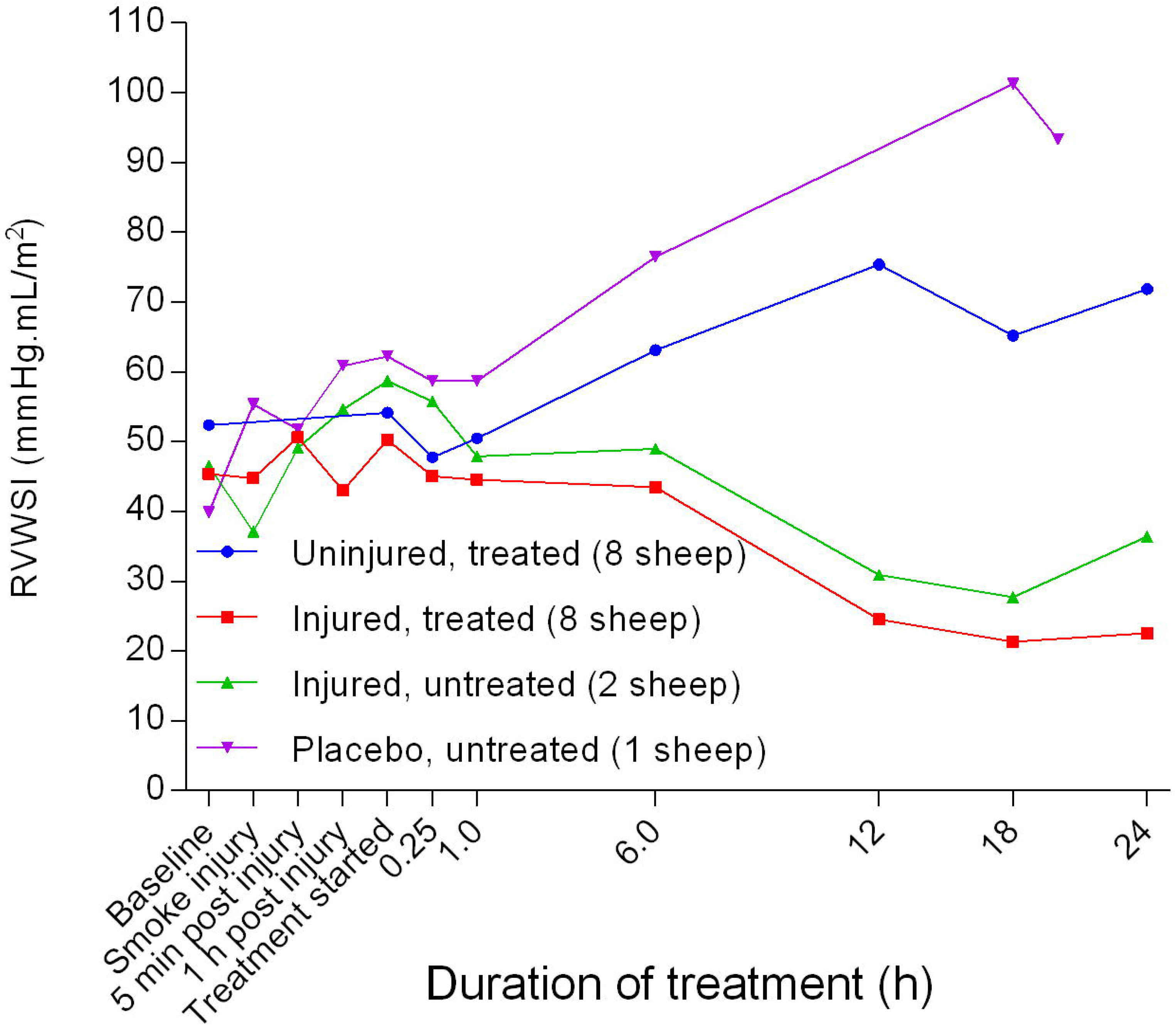
Right ventricular stroke work index (RVWSI) (Mean ± SD, error bars not shown) in smoke and non-smoke injured sheep that received extracorporeal life support as compared to that in untreated controls.

**Figure 34.**
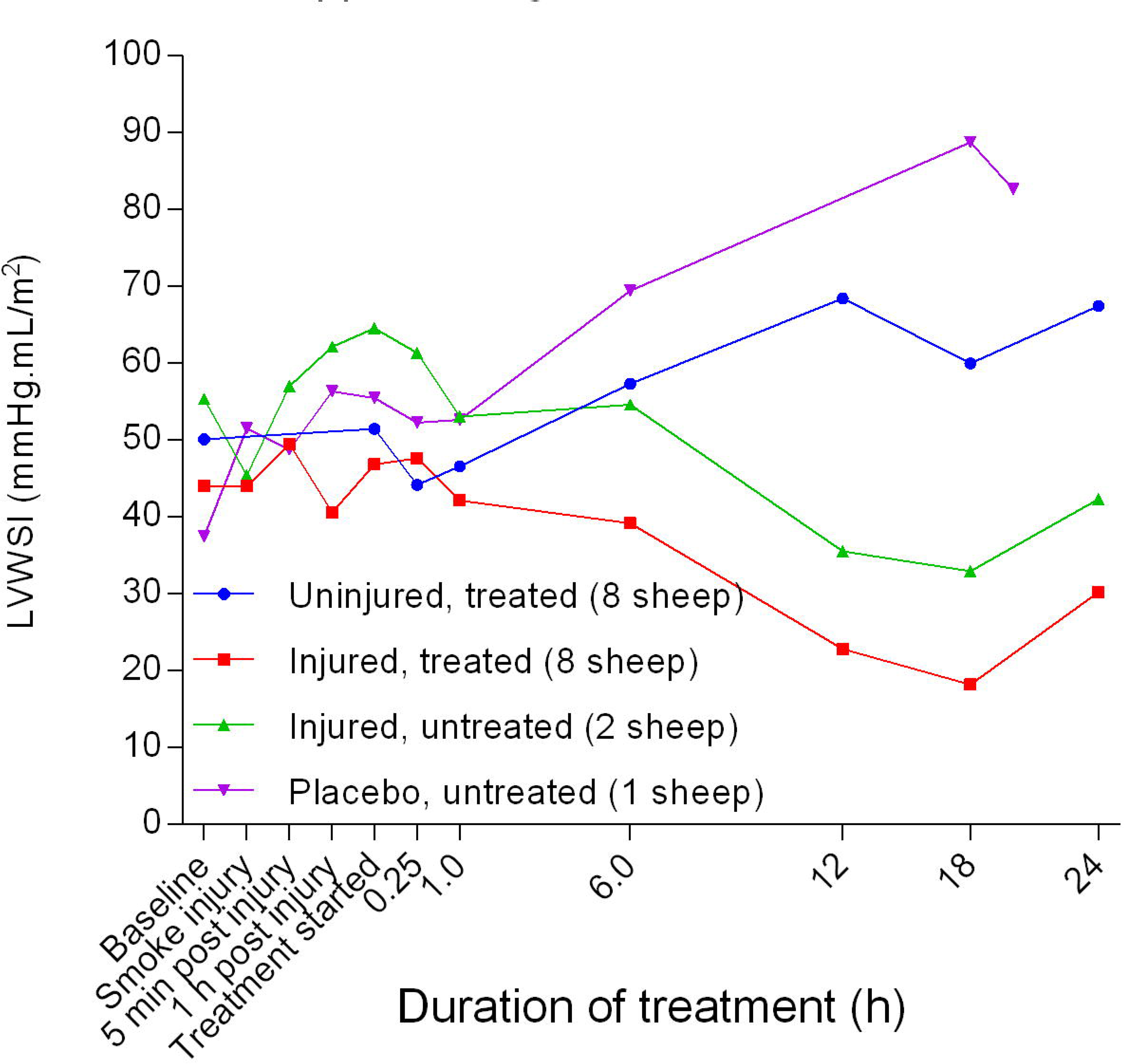
Left ventricular stroke work index (Mean ± SD error bars not shown) in smoke and non-smoke injured sheep that received extracorporeal life support as compared to that in untreated controls.

Following a decrease in the coronary perfusion pressure (CPP) from baseline in the smoke-injured sheep, there was a subsequent increase in this parameter within 5 minutes prior to a sustained decrease up to 18 hours of treatment, followed by another increase for the subsequent 6 hours (Figure 35). There was a significant difference in CPP (*p* = 0.0018) between the uninjured/treated and injured/treated groups and CCP in the placebo/untreated sheep remained relatively stable after an initial, subtle increase.

**Figure 35.**
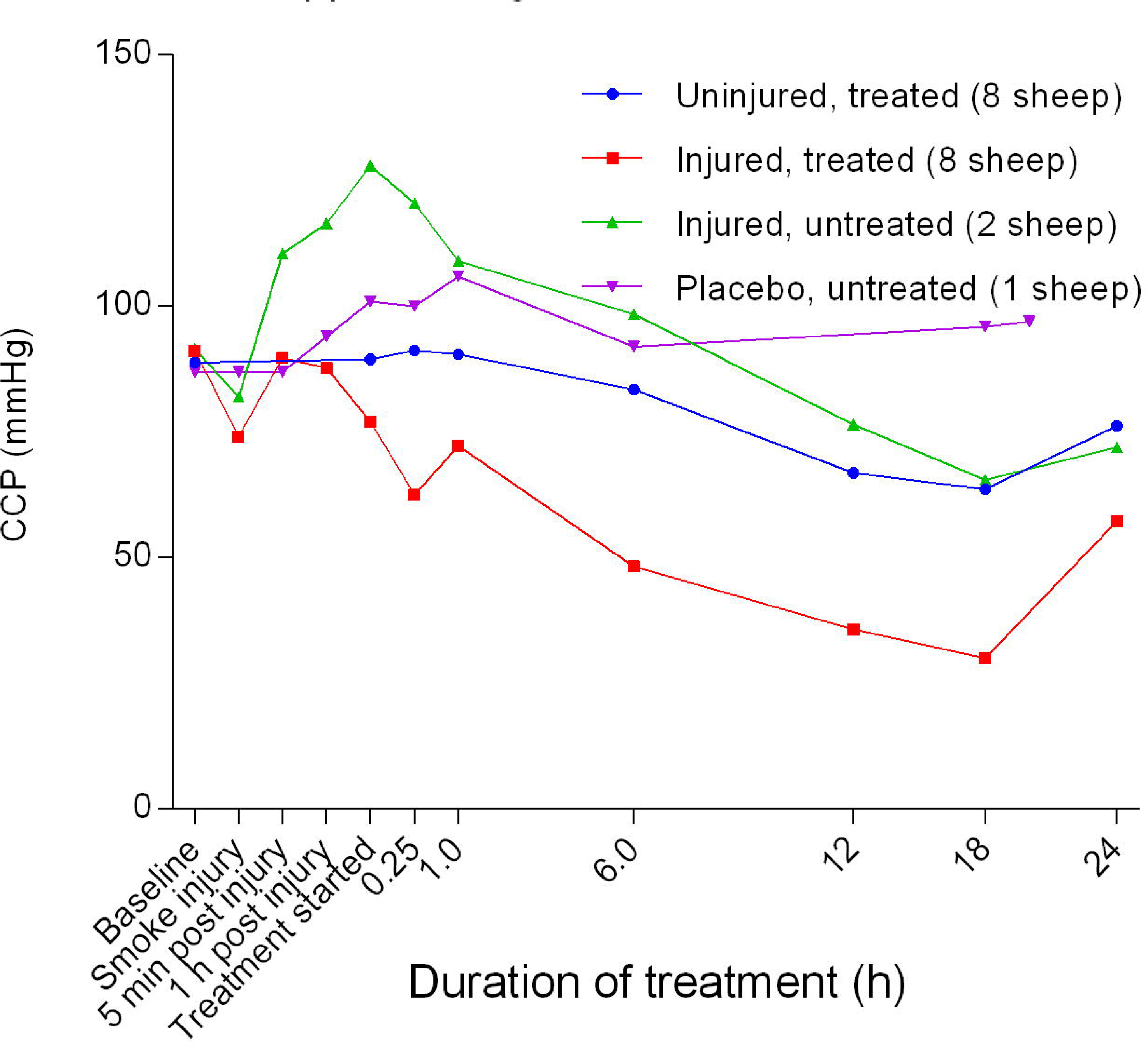
Coronary perfusion pressure (CPP) (Mean ± SD, error bars not shown) in smoke and non-smoke injured sheep that received extracorporeal life support alongside as compared to that in controls.

There was an initial subtle decrease in arterial oxygen content (C_a_O_2_) from baseline in all groups before a sustained increase in the injured/untreated group, a steady level in the placebo/untreated sheep, and a sharp trough in the injured/treated and uninjured/treated groups (Figure 36). Following the trough, the C_a_O_2_ of the injured/treated group gradually returned to baseline levels while that of the uninjured/treated group continued along a downward trend. There was a significant difference (*p* < 0.0085) in C_a_O_2_ between the uninjured/treated and injured/treated groups.

**Figure 36.**
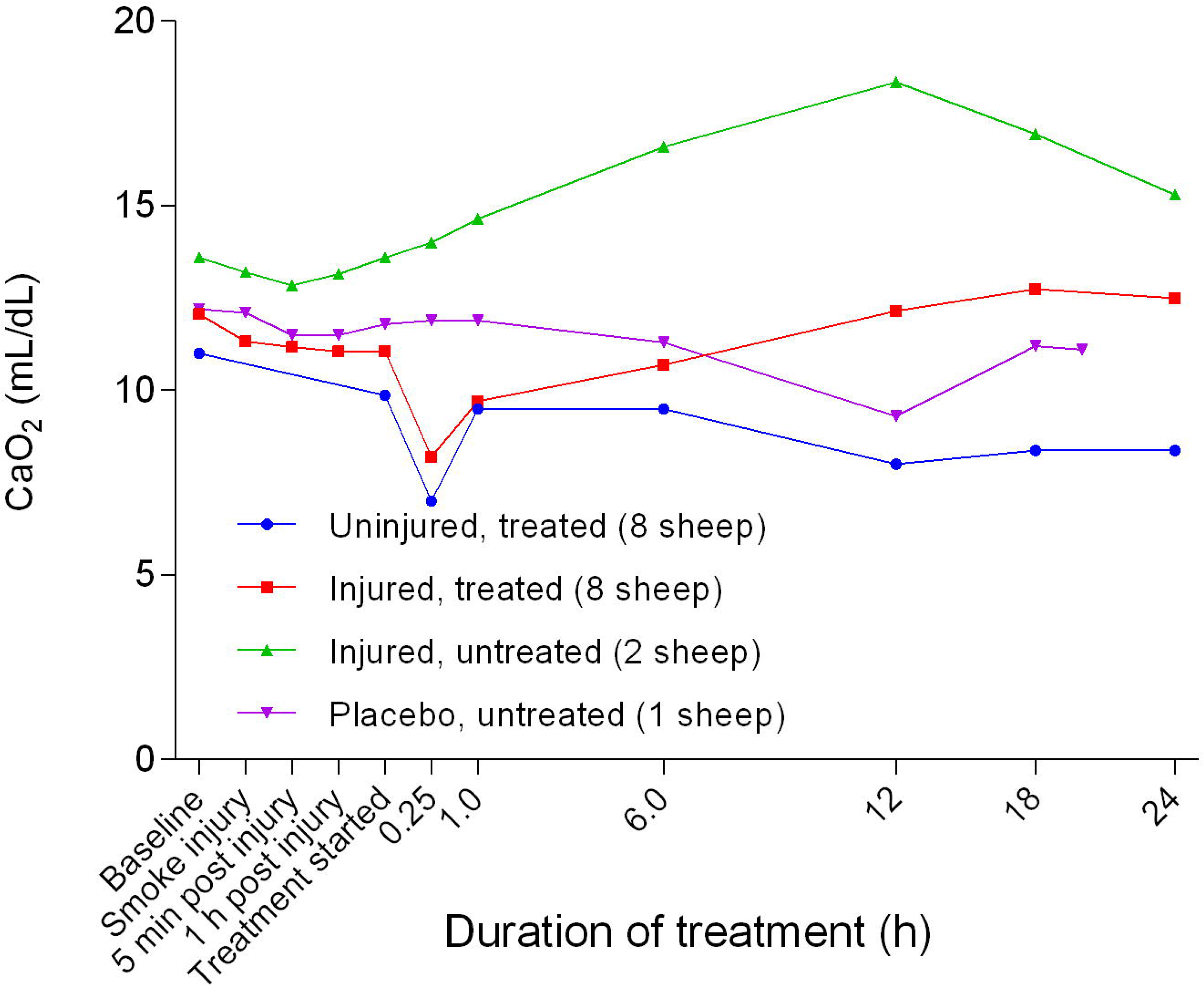
Arterial oxygen content (Mean ± SD error bars not shown) in smoke and non-smoke injured sheep that received extracorporeal life support as compared to that in untreated controls.

There was a slight decrease in the oxygen delivery index (DO_2_I) in all groups 1 hour after treatment before a further marked decrease, except for the placebo/untreated sheep (Figure 37). There was a significant difference (*p* = 0.0013) in DO_2_I between the uninjured/treated and injured/treated groups. The injured/treated group had the lowest DO_2_I compared with the other groups while the placebo/untreated sheep maintained the highest DO_2_I profile. The oxygen extraction index (O_2_EI) decreased in all groups before plateauing after approximately 6 hours of treatment. Further, there was a significant difference (*p* = 0.0247) in O_2_EI between the injured/treated and uninjured/treated groups (Figure 38). The O_2_EI in the injured/treated and injured/untreated groups was consistently lower and higher, respectively, compared with those of the other groups.

**Figure 37.**
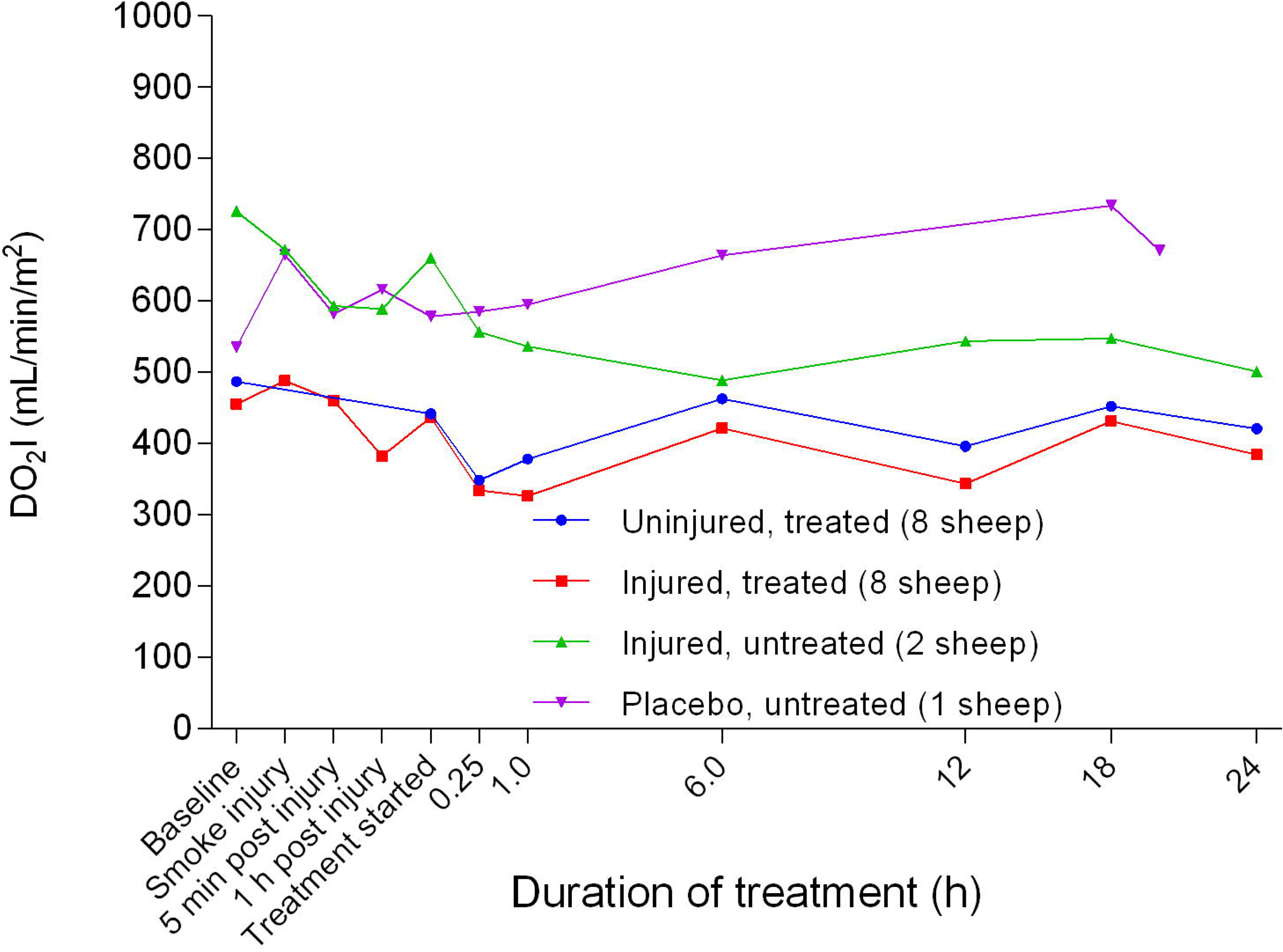
Oxygen delivery index (DO_2_I) (Mean ± SD error bars not shown) in smoke and non-smoke injured sheep that received extracorporeal life support as compared to that in untreated controls.

**Figure 38.**
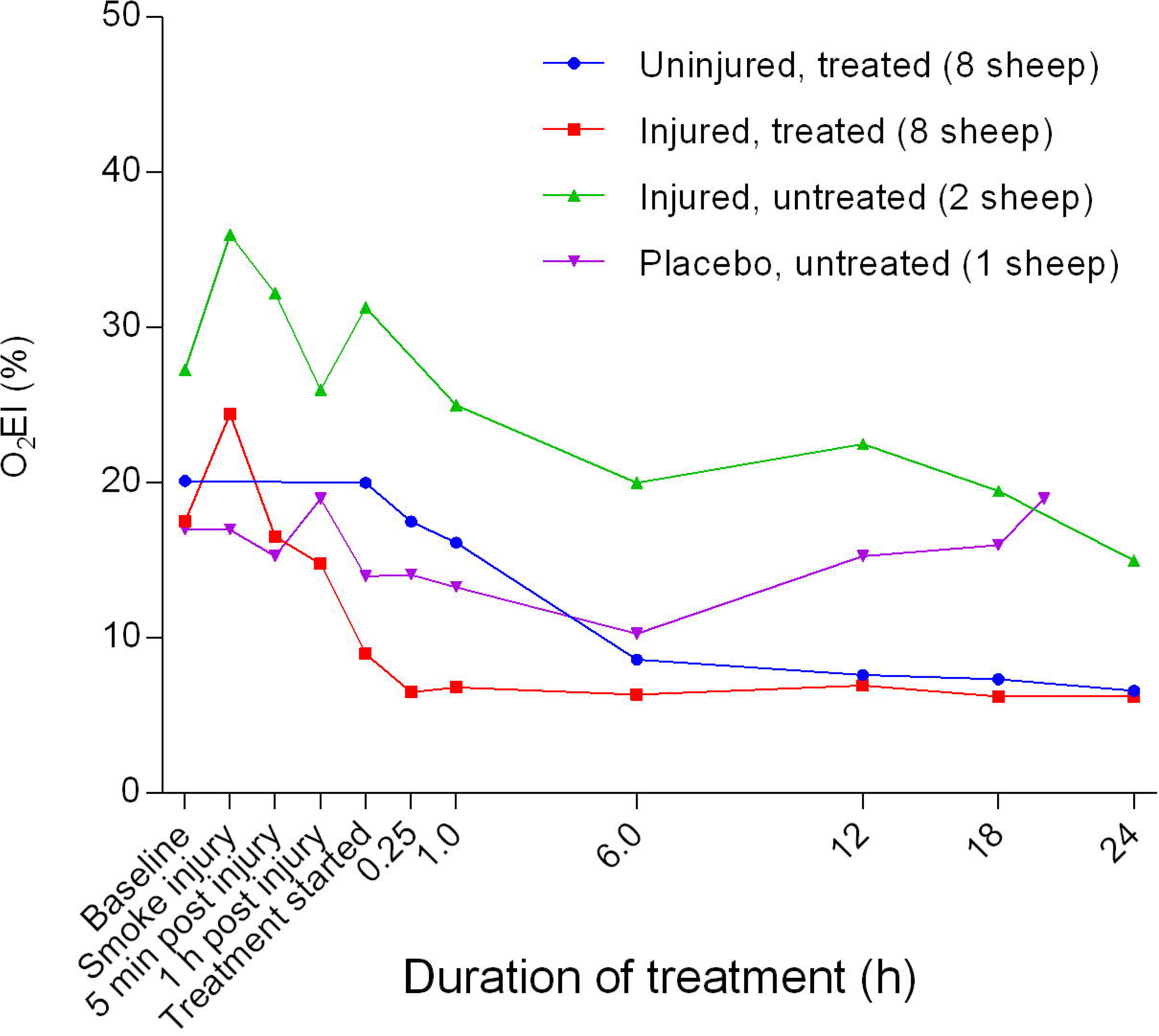
Oxygen extraction index (O_2_EI) (Mean ± SD error bars not shown) in smoke and non-smoke injured sheep that received extracorporeal life support as compared to that in untreated controls.

### Fluid input and urine output

There was a variation in the volume of intravenous fluids administered to sheep in the different experimental groups. The injured/treated sheep had the highest fluid requirements, while the placebo/untreated sheep required the least (Figure 39). There was a significant difference (*p* < 0.0001) in fluid requirements between uninjured/treated and injured/treated sheep. The injured/untreated and injured/treated groups produced the least and most urine on average, respectively (Figure 40). There was no significant difference in urine output between the uninjured/treated and injured/treated groups.

**Figure 39.**
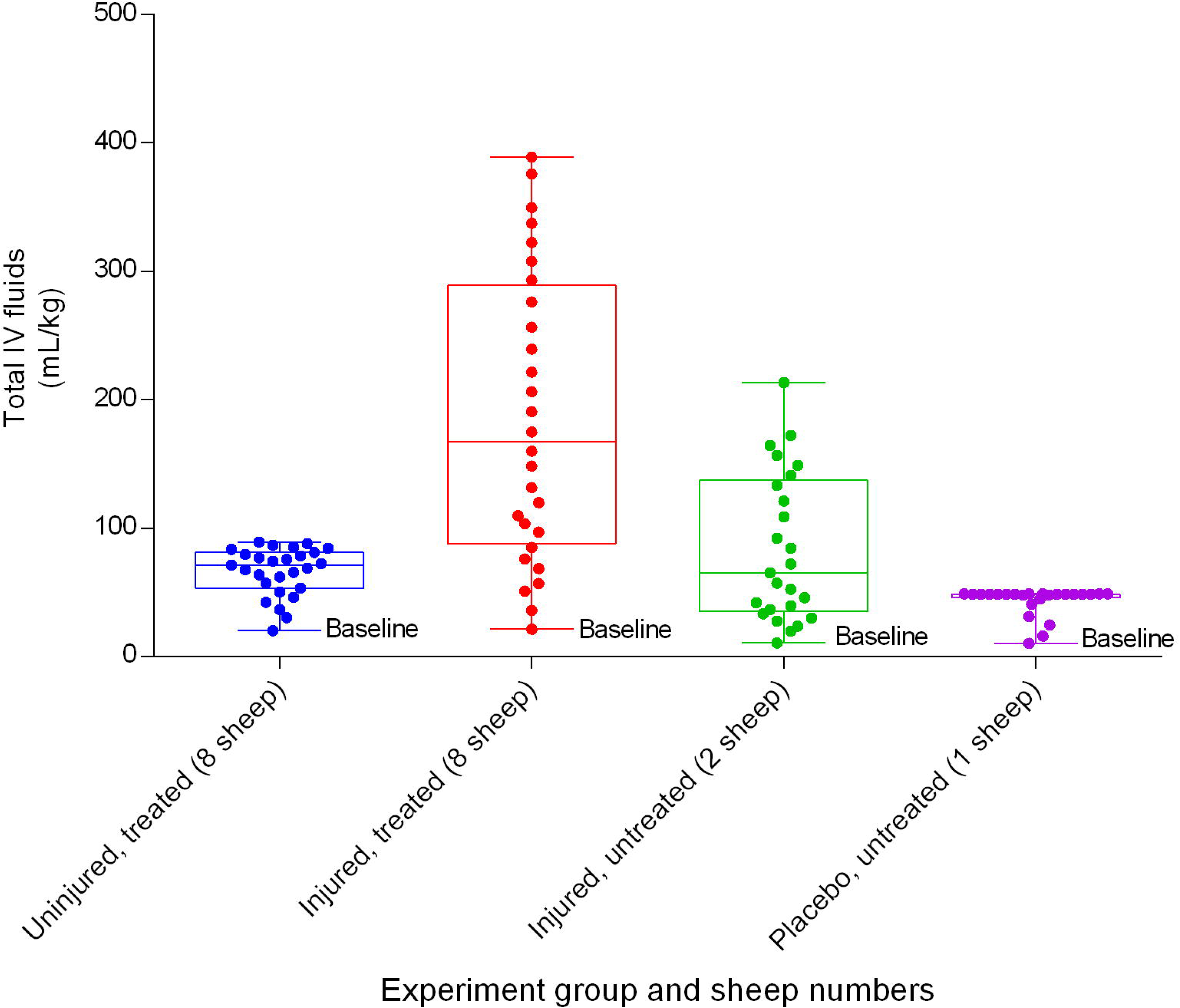
Total running intravenous fluids per unit body weight (Mean ± SD) in smoke and non-smoke injured sheep that received extracorporeal life support as compared to those in untreated controls over a 24h period.

**Figure 40.**
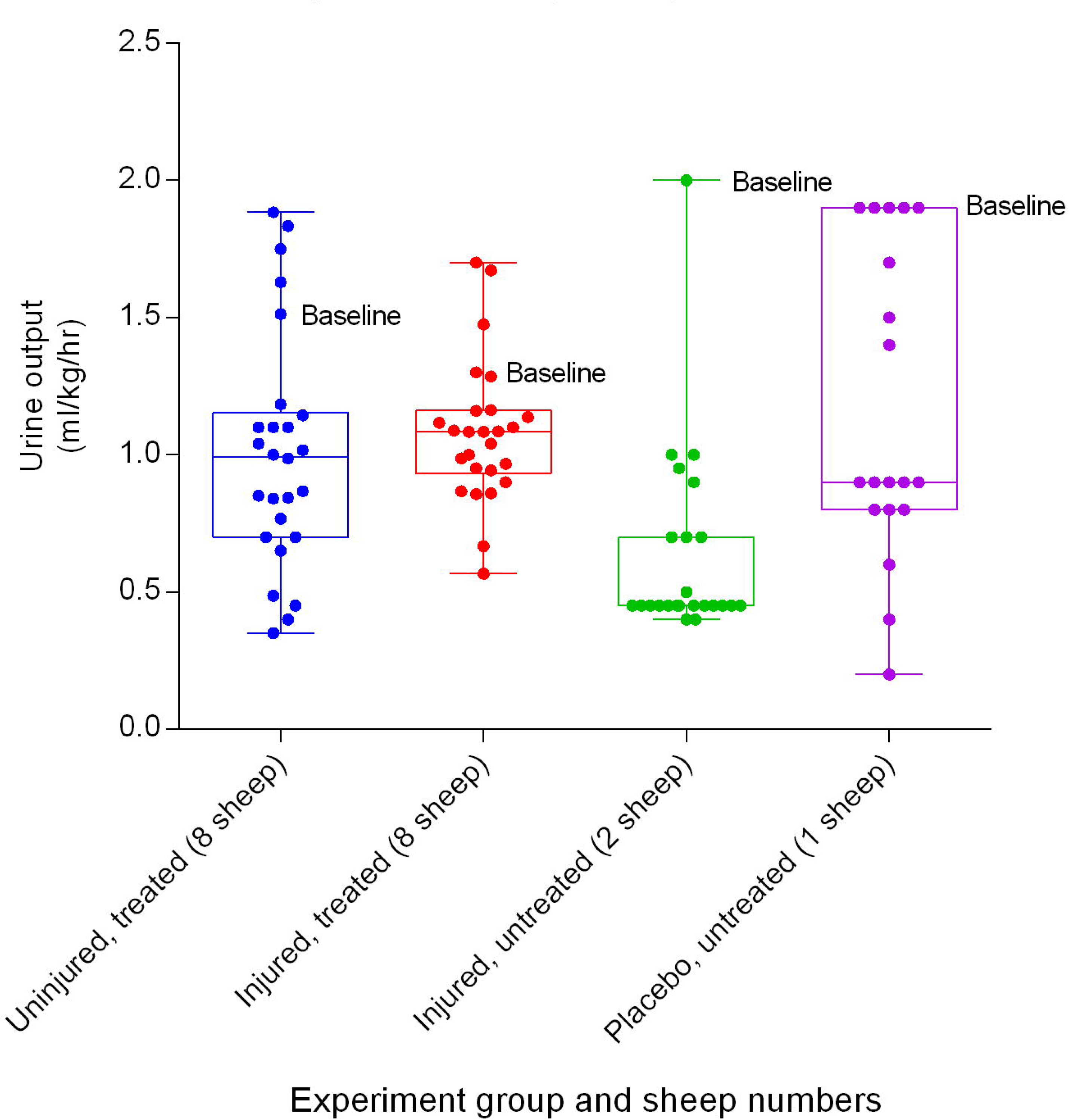
Urine output (Mean ± SD) in smoke and non-smoke injured sheep that received extracorporeal life support as compared to that in untreated controls over a 24h period. Lower and upper bar limits correspond to minimum and maximum urine output, respectively.

### Anaesthetics

There was a significant difference (*p* < 0.0001) in the amount of alfaxalone required between the uninjured/treated and injured/treated groups (Figure 41). The uninjured/treated group required more alfaxalone on average and the injured/untreated group required the least amount on average. Ketamine requirements differed between groups, with the injured/untreated group requiring the highest amount on average and the injured/treated group requiring the least (Figure 42). There was no significant difference in the quantities of ketamine required between the uninjured/treated and injured/treated groups, but there were significant differences in midazolam requirements (Figure 43), occurred between the uninjured/treated and injured/treated groups (*p =* 0.0067).

**Figure 41.**
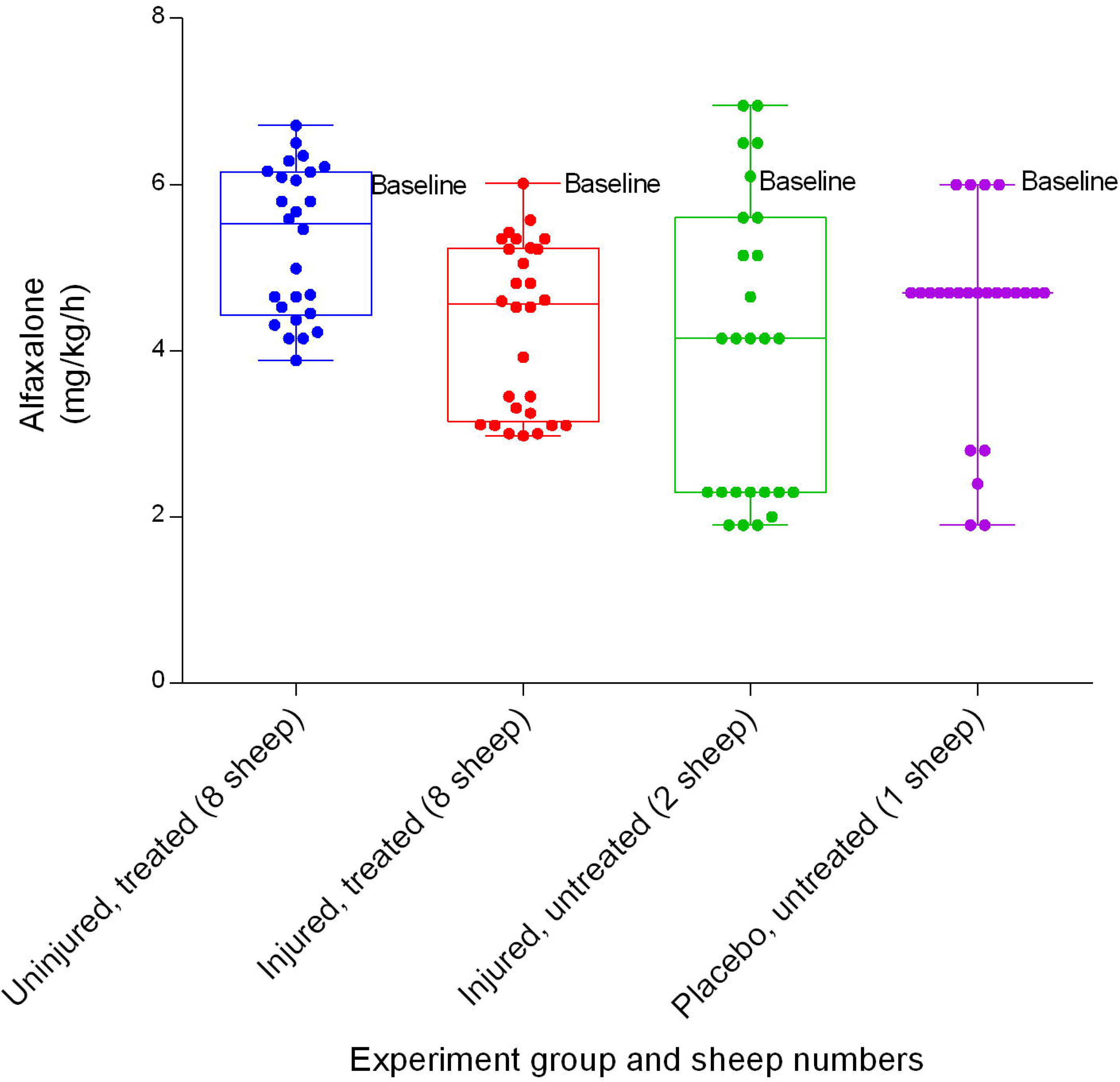
Alfaxalone infusion (Mean ± SD) indicating the titration range in smoke and non-smoke injured sheep that received extracorporeal life support as compared to that in untreated controls over a 24-hour period.

**Figure 42.**
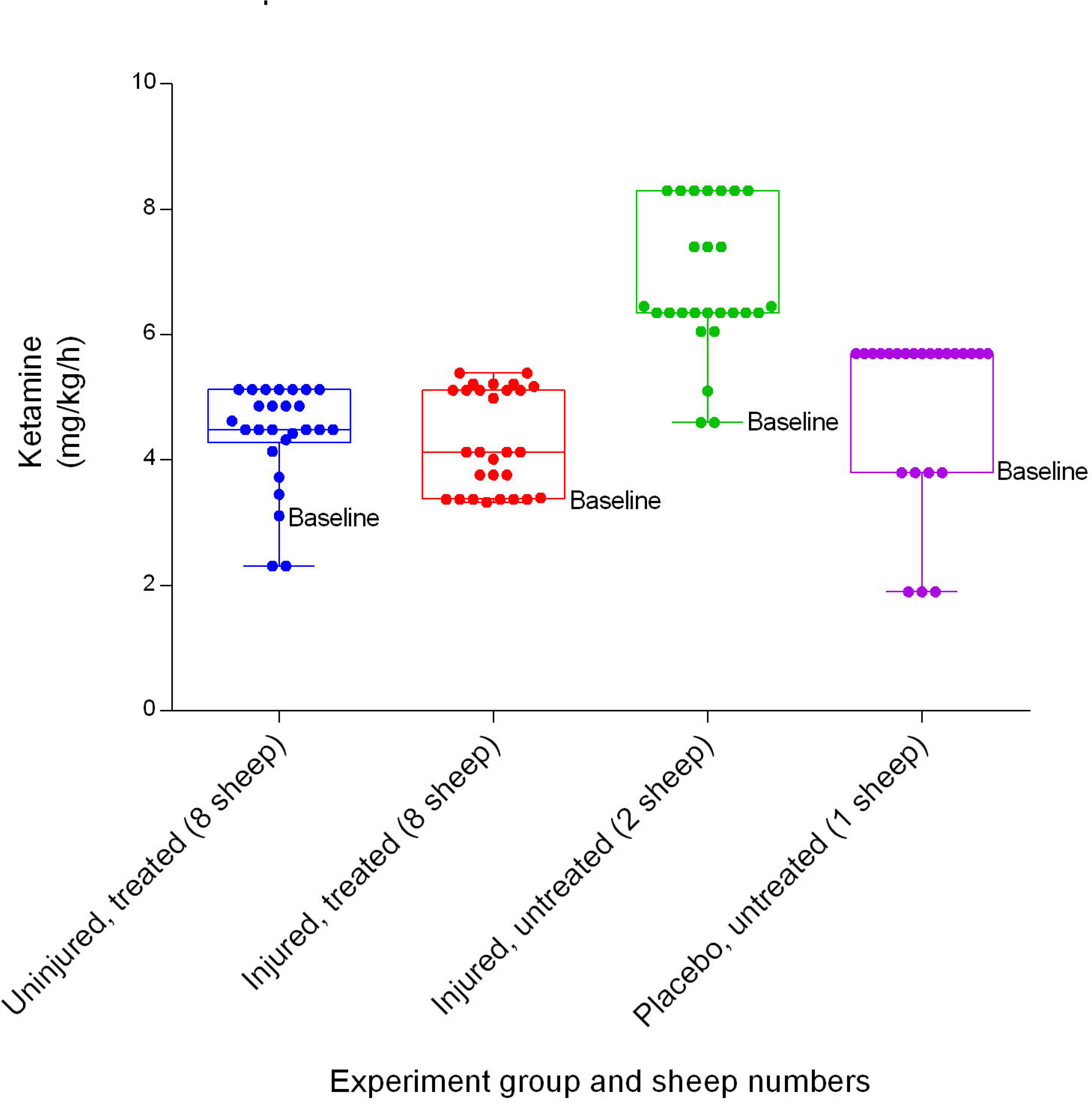
Ketamine IV infusion (Mean ± SD) indicating the titration range in smoke and non-smoke injured sheep that received extracorporeal life support as compared to that in untreated controls over a 24h period.

**Figure 43.**
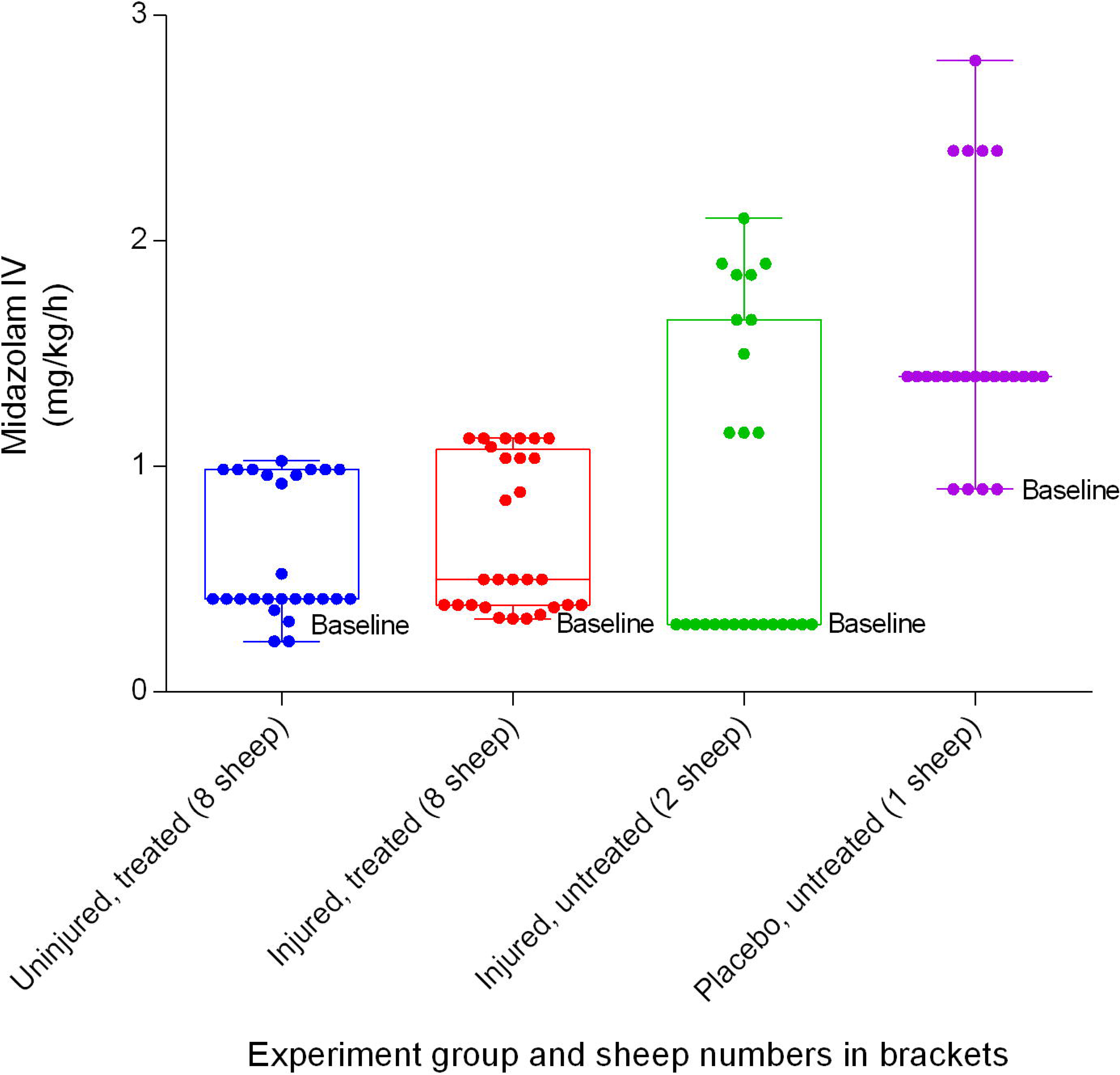
Midazolam IV infusion (Mean ± SD) indicating the titration range in smoke and non-smoke injured sheep receiving extracorporeal life support as compared to that in untreated controls over a 24-hour period.

### Anticoagulation

There were no significant differences in heparin infusion doses between the uninjured/treated and injured/treated sheep. Heparin requirements for the placebo/untreated group were the lowest (Figure 44). Activated clotting time increased sharply from baseline during pre-treatment and peaked after 1 hour of treatment, before decreasing sharply and plateauing (Figure 45). There were no significant differences in activated clotting time between groups.

**Figure 44.**
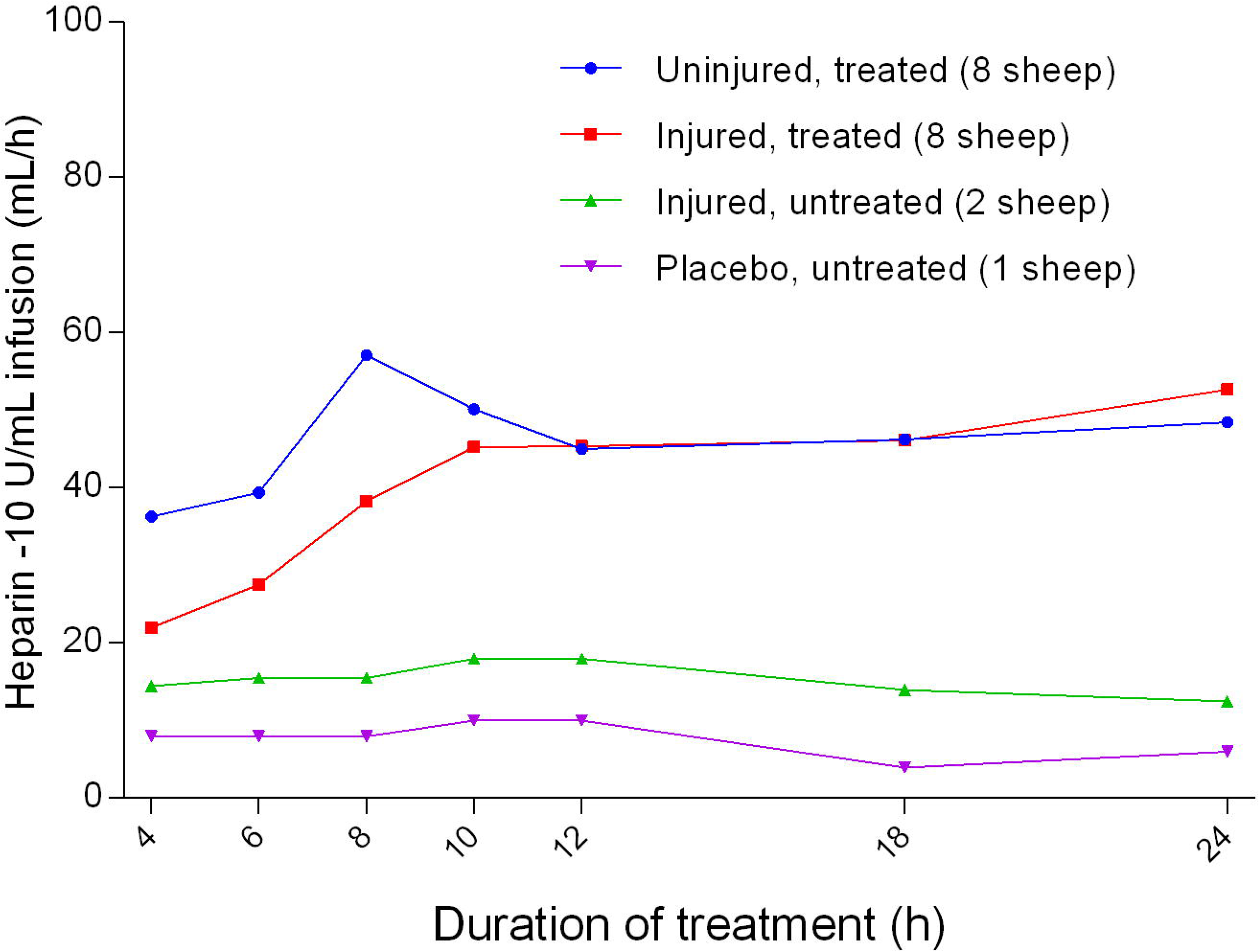
Heparin (10 U/mL) infusion rate (Mean ± SD, error bars not shown) in smoke and non-smoke injured sheep that received extracorporeal life support as compared to that in untreated controls.

**Figure 45.**
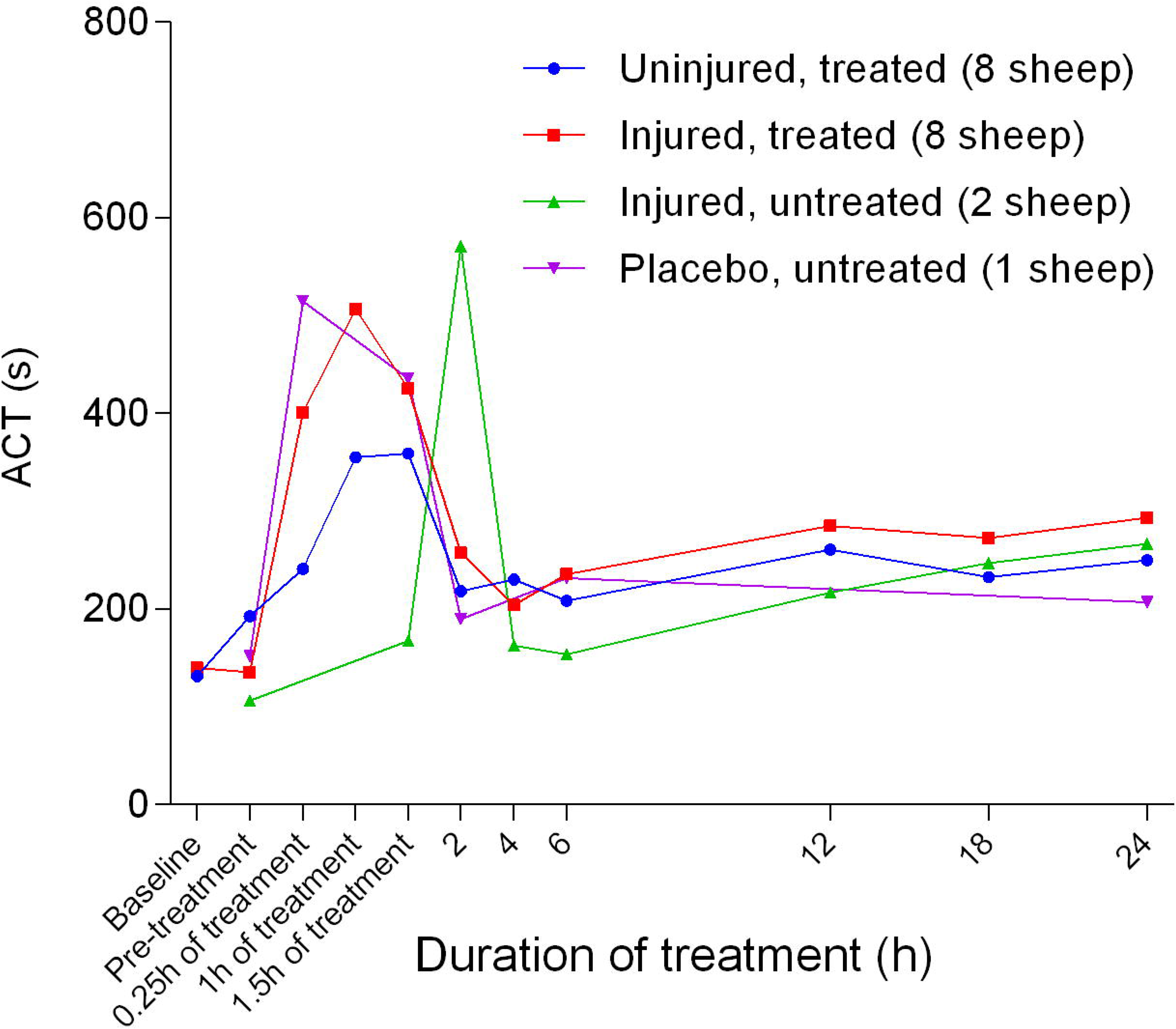
Activated clotting time (ACT) (Mean ± SD, error bars not shown) in smoke and non-smoke injured sheep that received extracorporeal life support as compared to that in untreated controls.

### ECLS circuit observations

There were significant differences in the ECLS pump speed (Figure 46), blood flow (Figure 47), and pressure differential (Figure 48) between the uninjured/treated and injured/treated groups. Further pump speed, blood flow, and pressure differential were significantly different (*p* = 0.0022, *p* = 0.0095 and *p* = 0.0041, respectively) between the two groups that received ECLS. These parameters in the uninjured/treated group were consistently higher than those of the injured/treated group.

**Figure 46.**
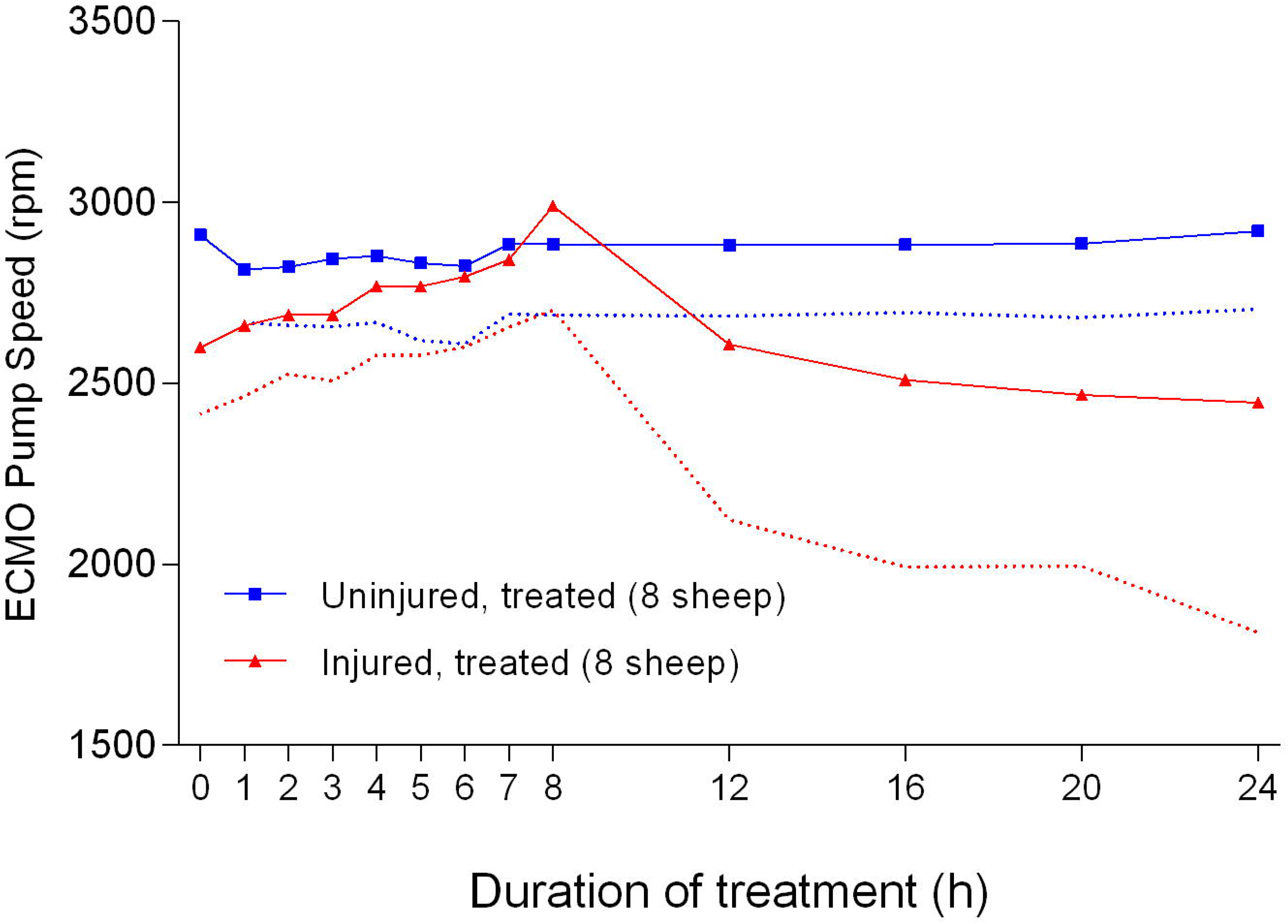
Extracorporeal life support pump speed (Mean ± SD) for smoke and non-smoke injured sheep that received treatment.

**Figure 47.**
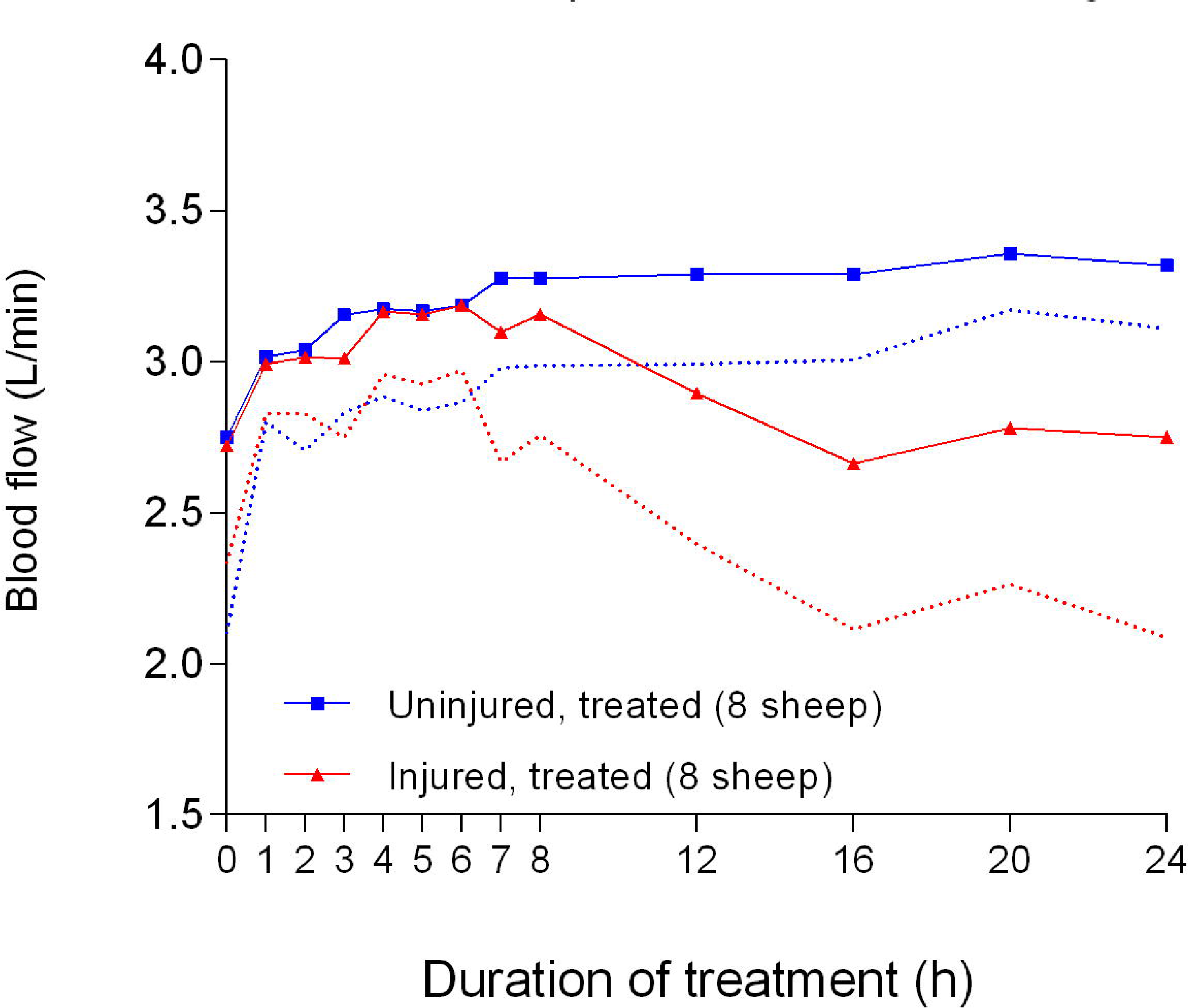
Blood flow (Mean ± SD) through an extracorporeal life support pump for smoke and non-smoke injured sheep that received treatment.

**Figure 48.**
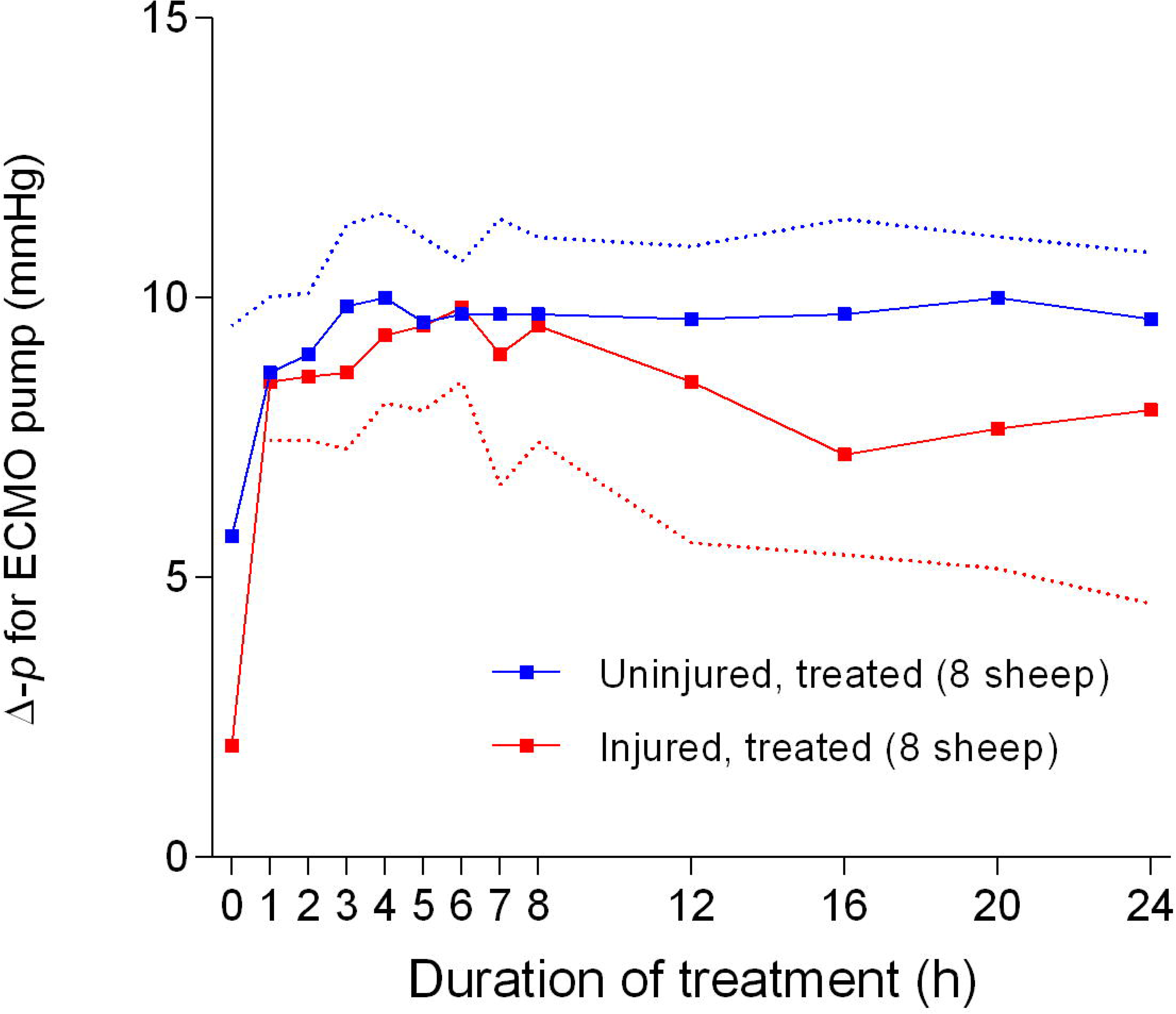
Pressure differential (Δ-p) (Mean ± SD) across the extracorporeal life support pump for smoke and non-smoke injured sheep that received treatment

### Body temperature

Body temperature in the untreated groups gradually increased from baseline levels and plateaued at approximately after 6 hours of treatment, and remained higher than that for the treated groups (Figure 49). There was no significant difference in body temperature between the treated groups.

**Figure 49.**
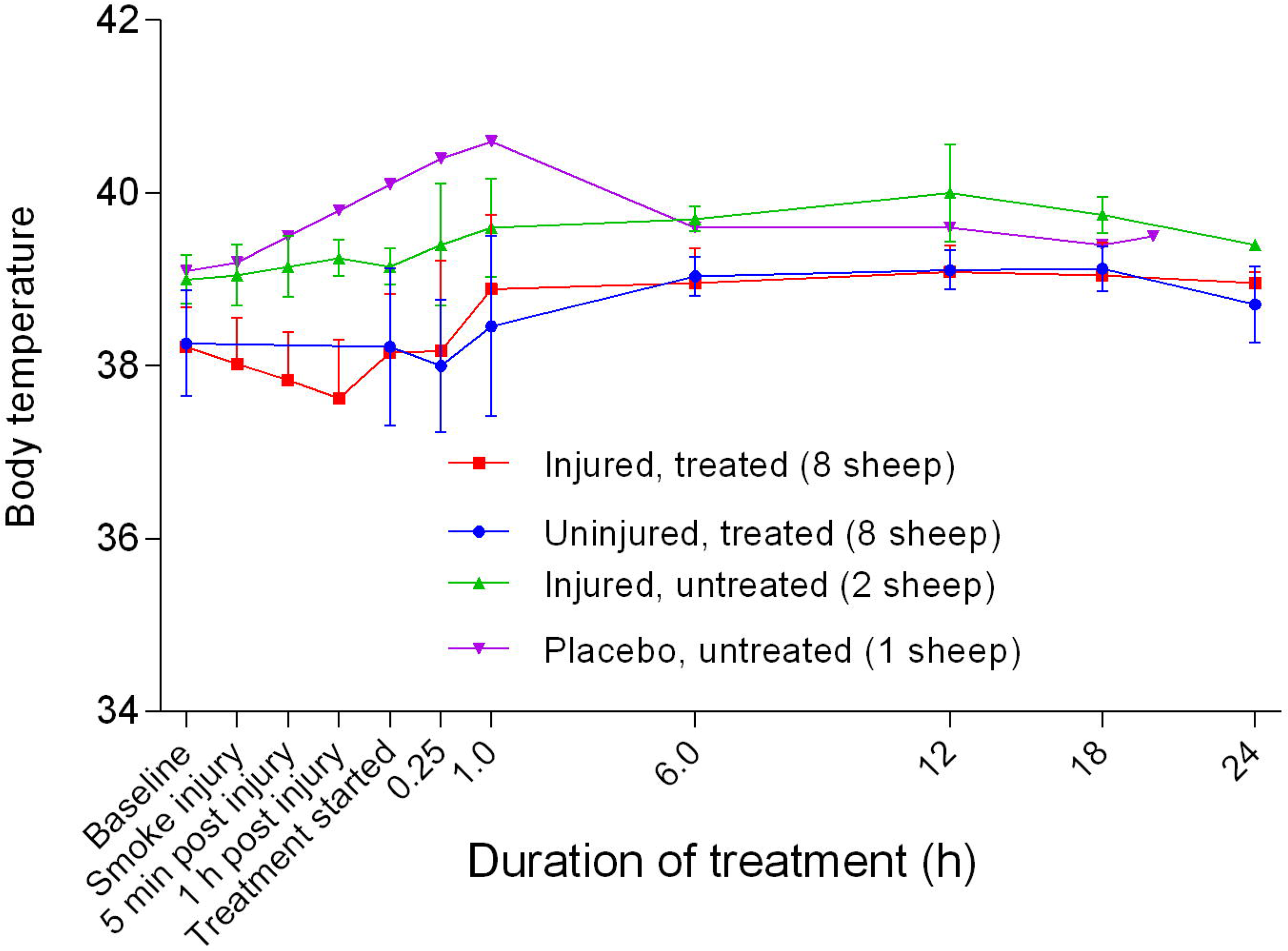
Body temperature (Mean ± SD) of smoke and non-smoke injured sheep that received extracorporeal life support as compared to that of untreated controls.

### Inflammatory cells and cytokines

Data were initially available in abstract form and subsequently as a full publication on inflammatory cell infiltration into the lung tissue with a trend toward increased lung injury in sheep that inhaled smoke, revealing damage to the bronchiolar lining and infiltration of inflammatory cells ^5, 21–23^.

## Discussion

The results of this study agree with and confirm earlier preliminary observations that ECLS causes a decrease in pulmonary compliance (Figure 2) over time^19^. It was expected that the injured sheep would have relatively lower SpO_2_ readings (Figure 3) compared with the other groups because of episodes of hypotension with hypoxemia, which can affect pulse oximeter function^24^. The relatively low etCO_2_ in the injured sheep suggested that the sheep may have hyperventilated (Figure 4), the causes of which were evaluated with respect to the reactive oxygen species or superoxide dismutase activity by a team from the source study ^25, 26^.

The relatively low blood pH in the injured/treated sheep as depicted in Figure 5, suggested that the sheep tended to have metabolic acidosis, as the same group of animals also had low etCO_2_. This also implies that there was no respiratory component that contributed to the observed acidosis. The low pCO_2_ in the uninjured/treated sheep could be a result of hyperventilation and the high pCO_2_ in the injured/untreated sheep suggested that CO_2_ clearance was curtailed by injury (Figure 6).

The treatment of the sheep contributed to lung injury by causing deterioration of pO_2_ (Figure 7). The low pO_2_ translated to a low partial arterial oxygen pressure/inspired oxygen proportion (PaO_2_/FiO_2_ ratio), which was much worse in the injured sheep. This finding showed that ECLS contributed to the deterioration of the PaO_2_/FiO_2_ ratio in the injured/treated group of sheep, a novel finding that was also unexpected in the primary study (this has since been replicated in a more recent study^9^). It could also be argued from the data that perhaps VV ECMO was performed in a suboptimal manner, considering that the sheep oxygenation appeared to be less effective on ECMO than on the ventilator alone, but this remains to be explored further in future.

Further, the relatively higher levels of [Hb] in the injured sheep suggested that these animals could have been dehydrated as a secondary consequence of excessive fluid loss due to inflammation and increased vascular permeability^27^, despite intravenous fluid replacement (Figure 8). However, blood total protein and albumin levels, which are better predictors of dehydration in sheep^28^, were not measured.

The inverse decrease in FO_2_Hb (Figure 9) relative to FCOHb (Figure 10) following smoke injury was expected and is a finding that is in agreement with other studies^27, 29, 30^. It has recently been demonstrated that FCOHb is not correlated to the extent of lung injury^27^. The gradual decrease in MetHb (Figure 11) was probably caused by the enzymatic activity of methaemoglobin reductase^31^ and the higher Hct observed in the injured sheep (Figure 12) could have been due to dehydration because Hct was measured by an automated method.

As presented in Figure 14, the [Ca^2+^] was lower than the published normal level of 2.4 mmol/L ^32^. Stress associated with yarding of the sheep and phosphorus imbalance in feed are the most likely suggested causes of low [Ca^2+^]^33^. Fasting the sheep for 24 hours prior to the experimental procedures could also have contributed to the relatively low [Ca^2+^]. The increase in Cl^-^ beyond the normal range of 105–110 mmol/L^32^ during the experiments (Figure 15) suggests that the sheep may have developed respiratory alkalosis. Hyperventilation or metabolic acidosis resulting from sustained salivary loss of sodium bicarbonate that was more severe in the injured/treated group may have played a role in hyperchloraemia, because Cl^-^ is known to replace HCO_3_^-^ when the latter is lost from the body^34, 35^. Baseline [K^+^] in all the sheep (Figure 16) was below the published normal range of 4–5 mmol/L^32^ and this relative hypokalaemia may have been related to low K^+^ in the diet^36–38^. The normal anion gap (Figure 17) with decreased HCO_3_^-^ (Figure 21) confirmed the presence of hyperchloraemic acidosis in all but the placebo/untreated sheep. The cause of the hyperchloraemia was likely the prolonged administration of 0.9% NaCl. Although normal [Glu] in ruminants is usually lower than that for other species, its relative progressive increase in the injured sheep (Figure 18) may have been related to stress and severe pain associated with injury or the development of enterotoxaemia^39, 40^. The relative increase in [Lac] beyond the reported normal range of 1–2 mmol/L in the injured sheep (Figure 19) suggested dehydration, trauma, and sepsis^41^. In particular, sepsis is a concern related to the sub-optimal rumen function, leading to loss of its buffer effect and an increase in the number of anaerobic bacteria with prolonged hypomotility, such as that which occurs during long-duration anaesthesia. Therefore, the increases in both [Glu] and [Lac] are consistent with severe injury.

The elevated Base (ecf) above +2 mmol/L for most of the first 12 hours in the placebo/untreated and injured/untreated sheep suggested that the sheep were metabolically alkalotic ^39^, before returning to normal levels (Figure 20). The relatively low Base (ecf) (less than −2 mmol/L) was consistent with loss of HCO_3_^-^ and the tendency of developing metabolic acidosis ^39^ in the injured/treated sheep. The marked decrease in [HCO_3_^-^] in the injured sheep was consistent with metabolic acidosis and was more severe in the injured/treated group, as illustrated in Figure 21, thereby suggesting that ECLS was a contributing factor.

The resting HR of sheep is 50–80 beats/min^32^. In a study that instrumented conscious sheep, the baseline heart rate was registered as 106 ± 9 beats/min^42^. In the present report, all of the sheep had a relatively high HR, thereby suggesting that stress and pain were contributing factors (Figure 22). The gradual decrease in HR during the course of the experiments was consistent with the effects of anaesthesia^32^.

In sheep, a mean arterial pressure below 60 mmHg indicates inadequate tissue perfusion^32^. Although the MAP values in the injured sheep were lower than for the uninjured sheep (Figure 23), MAP values were still within the published normal value of 70 mmHg^32^; the magnitude of injury was again a predictor of how low the MAP was. Another predictor for the severity of the injury was the mean pulmonary artery pressure, which was highest for the injured/treated sheep as illustrated in Figure 24. The baseline values for MPAP were higher than the 17 ± 1 mmHg reported in another study that used sheep for experiments^42^. The baseline CVP in all of the sheep in the present report was > 10 mmHg (Figure 25), which was much higher than the 5.5 ± 1.2 mmHg reported elsewhere^42^ in instrumented conscious sheep and a novel finding in this study. Thus, in this study, the severity of injury and treatment contributed to the CVP elevations found among the sheep.

There was a benefit of ECLS treatment for SvO_2_, as it remained high for both the injured/treated and uninjured treated groups (Figure 26). The consistently low SvO_2_ in the injured/untreated group was expected because of the slightly reduced cardiac output in this group; however, this level of SvO_2_ was still higher than that reported in other studies^42^. Smoke injury was associated with a sustained decrease in cardiac output in all the sheep that were exposed to smoke. As in CCO changes (Figure 27), the SV (Figure 28), SVI (Figure 29) and CI (Figure 30) all had similar profiles for different groups, with the injured sheep having lower values. The decrease in SVRI in all of the sheep at a later stage in the experiments suggested that there was systemic vasodilation (Figure 31). In contrast, the increase in PVRI in the injured sheep suggested that vasoconstriction was caused by exposure to smoke injury (Figure 32). The exposure to smoke injury worsened both RVSWI (Figure 33) and LVSWI (Figure 34) while there was an increase in both parameters in the uninjured sheep. Reduced RVSWI is associated with poor functioning of the right ventricle^43, 44^ and LVSWI is a reliable parameter for left ventricular function^45^.

The reduction in coronary perfusion pressure in the injured/treated—and to a certain extent the uninjured/ treated sheep—suggested that ECLS contributed to the decrease in CPP, in addition to smoke injury (Figure 35). The CPP is an indicator of myocardial perfusion and has been proposed as a drug target during resuscitation^46^. The observations in the present study support the suggestion that CPP could be used to predict the severity of injury in sheep.

Further, the apparent increase in CaO_2_ in the injured sheep (Figure 36) could have been due to the relative increase in [Hb] secondary to dehydration as illustrated in Figure 8. The low DO_2_I in the injured/treated and uninjured/treated groups suggested that ECLS also contributed to this, in addition to smoke, based on the relatively higher DO_2_I in the injured/untreated sheep (Figure 37). Interestingly, the O_2_EI (Figure 38) had a comparable profile to that of the PaO_2_/FiO_2_ ratio and could also be used to predict the contribution of ECLS to smoke-related injury.

The smoke-injured sheep required considerable amounts of intravenous fluids (Figure 39) to compensate for the losses from pulmonary exudation and inflammation^27, 29^. The mean urine production in all groups (Figure 40) was marginally lower than the published normal of 1.2 mL/kg/h^32^ but still considered to be within the acceptable range for this cohort of sheep. Moreover, the dosage of anaesthetic drugs used was considered adequate for the experiments (Figures 41, 42 and 43). In addition, heparin infusion (Figure 44) was indicated to prolong the activated clotting time (Figure 45) in order to minimise the risk of thrombosis during intravascular procedures^47^.

The reduction in the ECLS pump speed (Figure 46), flow (Figure 47), and pressure differential (Figure 48) could have resulted from systemic hypotension contributing to low amounts of blood to the pump. The ECLS was configured such that the centrifugal pump pulled blood from the inferior vena cava and returned it into the right atrium; therefore, if the circulating volume was low, the flow would decrease for a given pump speed and in this case, both rpm and flow would reduce. Centrifugal ECLS pumps are known to be preload dependent and afterload sensitive^48^, thereby making rpm and flows directly proportional to each other. The reason for the systemic hypotension remains undetermined. It is possible that an unknown pulmonary component or product produced in the smoke-damaged lungs of the sheep played a role. It must be noted that the body temperature of the sheep was generally within the physiological range (Figure 49).

Certain observations regarding this study could affect the interpretation of red blood cell indices and their derivatives. For example, animals differ from humans in that estimated changes in plasma volume is preferably determined by changes in packed cell volume (PCV) or haemoglobin concentration and total plasma protein (TPP)^49–51^. Moreover, in animals, there is a wider range of normal PCV than TPP ^52^. In critical care for domestic animals, the change in both PCV and TPP is most useful as a crude index of change in plasma volume^53^. A centrifuge that spins minute amounts of blood for rapid, cost-effective determination of PCV and TPP permits instant adjustments in an animal’s fluid needs. However, measurements of PCV and TPP were not conducted in the primary study. As with all data that are collected with different objectives, it was tedious to align certain time points with real-time observations made in the laboratory, particularly for data that was manually input. There was also no information regarding pre-anaesthetic blood tests.

An additional limitation of this study is related to the overall objective of providing useful information relevant to the sheep ECMO model in particular, and to the scientific community interested in large animal experimentation and veterinary medicine in general. Because the method of data collection method has not been validated across several laboratories or research groups, it is considered relatively preliminary and further validation studies are required. Moreover, the number of sheep (Figure 1) was low and this was particularly so in the injured/untreated and placebo/untreated groups, thereby preventing comparisons between the treated and untreated sheep. A further limitation is that cytokine levels, as predictors of lung injury, were not quantified. Using ELISA assays to quantify cytokine levels proved difficult and the cost was prohibitive in the present study, although subsequent efforts were made by a few members of the original research group in this regard^5^. It is partially for this reason that pioneering studies^16^ have been proposed for the development of proteogenomic assays as an alternative to ELISA—to learn from circulating markers of acute inflammation in injured sheep used as models of intensive care, to understand critical illness. On another note, it can be argued that sheep ECMO data is already out there however, cardiac function and haemodynamic data has not been reported in sufficient detail in the manner that this study has. Also, this paper has many figures which some readers may see as being unfocused. The upside is that in recent times, academic journals now have sufficient capacity for lengthy, more informative and detailed reports by preferably having all data in one publication, rather than chopping it up into tight ‘salami’ publications. In the literature related to this burn and smoke inhalation model lung, function has probably been reasonably well reported, but cardiac function is probably less so – which makes this report to stand out strongly. Lastly, it can be construed that observational control groups are too small (n=1 or 2). Putting out data from small groups and placing it in graphs with experimental groups with n=8 may be seen as misleading, but cutting these observations from the other two groups completely would be a disservice to science, considering that few animals as possible are used for research purposes nowadays to prove a point.

Nevertheless, although ECMO treatment has been demonstrated to contribute to, and exacerbate the deterioration of pulmonary pathology by reducing lung compliance and PaO_2_/FiO_2_ ratio in sheep studies^5, 9, 22^, the understanding of ECMO in respiratory life-support in human medicine continues to grow in a positive direction overall; moreover, there is evidence that it is useful as a life-saving treatment. These novel observations from sheep could help in understanding similar pathology such as that which occurs in animal victims of smoke inhalation from house or bush fires, aspiration pneumonia secondary to tick paralysis, and in the management of COVID-19 in humans^54–56^.

The World Health Organization (WHO) has recently recognised and classified COVID-19, caused by the novel coronavirus SARS-CoV-2, as a global pandemic and public health emergency ^57, 58^. The National Institute of Allergy and Infectious Disease (NIAID) of the United States of America recognises that coronaviruses constitute a large group of viruses known to cause respiratory diseases, including the common cold^59^. However, in recent years, three novel members of this family of viruses have arisen from animals to cause severe and extensive infection and death in humans^60^. In addition to bats, a large number of coronaviruses are known to circulate in certain domestic animals like cats and occasionally spill over to humans and cause serious illnesses, such as the SARS coronavirus (SARS-CoV) that emerged in Southern China in 2002^61^. As a future perspective, the outcomes of the present study could be used to guide additional studies to enable the mortality indicators and prognostic indicators associated with ECMO and allied technology to be further evaluated and well understood in sheep and other experimental animals.

## Conclusions

The results of this study demonstrated that ECLS contributed to the worsening of pulmonary pathology by reducing lung compliance and PaO_2_/FiO_2_ ratio. The O_2_EI changes mirrored those of the PaO_2_/FiO_2_ ratio and decreasing CPP was a predictor of a greater magnitude of cardiopulmonary injury in sheep. These novel observations could help in further understanding similar pathology in other patients; for example, in the resuscitation of animals injured from in house or bush fires. A similar data acquisition approach could be used in evaluating the effectiveness of a given experimental or clinical intervention to further the understanding of the clinical condition being 25 studied and to aid in the formulation of treatments aimed at improving the survival of animal patients. In veterinary medicine, albeit now a considerably expensive and remote option, ECLS knowledge could complement the treatment of potentially reversible aspiration pneumonia, a secondary complication associated with both *Ixodes holocyclus* toxicity and laryngeal paralysis, in valuable companion animals and in critically ill humans who require respiratory support, like COVID-19 patients.

## Data availability

### Underlying data

Chemonges, Saul (2020), Cardiorespiratory physiological perturbations after acute smoke-induced lung injury and during extracorporeal membrane oxygenation support in sheep, v4, Dryad, Dataset, https://doi.org/10.5061/dryad.3r2280gd5.

This project contains the following underlying data:

C24H-01.xls

E24H-01 Saul.xls

E24H-02 Saul.xls

E24H-03 Saul.xls

E24H-04 Saul.xls

E24H-05 Saul.xls

E24H-06 - 4146 Saul.xls

E24H-07 Saul.xls

E24H-E24H-Activated Clotting Time (Saul).xls

E24H-Activated Clotting Time (Saul).xls

E24H-Anaesthetics, Inotropes & Anticoagulants (Saul).xls

E24H-Arterial Blood Gas Values (Saul).xls

E24H-BP, ventilation & haemodynamic data (Saul).xls

E24H-Calculated Resp+Haemodynamic Variables (Saul).xls

E24H-Fluids and Urine Production (Saul).xls

Fig1.tif, Fig2.tif, Fig3.tif, Fig4.tif, Fig5.tif, Fig6.tif, Fig7.tif, Fig8.tif, Fig9.tif, Fig10.tif, Fig11.tif, Fig12.tif, Fig13.tif, Fig14.tif, Fig15.tif, Fig16.tif, Fig17.tif, Fig18.tif, Fig19.tif, Fig20.tif, Fig21.tif, Fig22.tif, Fig23.tif, Fig24.tif, Fig25.tif, Fig26.tif, Fig27.tif, Fig28.tif, Fig29.tif, Fig30.tif, Fig31.tif, Fig32.tif, Fig33.tif, Fig34.tif, Fig35.tif, Fig36.tif, Fig37.tif, Fig38tif, Fig39.tif, Fig40.tif, Fig41.tif, Fig42.tif, Fig43.tif, Fig44.tif, Fig45.tif, Fig46.tif, Fig47.tif, Fig48.tif, Fig49.tif

SC24H 1+2 PF and Carboxy.xlsx SC24H-01 Saul.xls

SC24H-02 Saul.xls

SE24H-01 Saul.xls SE24H-02 Saul.xls

SE24H-03 Saul.xls

SE24H-04 Saul.xls

SE24H-05 Saul.xls

SE24H-06 Saul.xls

SE24H-07 Saul.xls

SE24H-08 Saul.xls

SE24H-Activated Clotting Time (Saul).xls

SE24H-ALL SO FAR (6) Saul.xls

SE24H-Anaesthetics, Inotropes & Anticoagulants (Saul).xls

SE24H-Arterial Blood Gases (Saul).xls

SE24H-Calculated Resp + Haemodynamic Variables (Saul).xls

SE24H-Fluids and Urine Production (Saul).xls

SE24H-Physiologic monitoring data (Saul).xls

Data are available under the terms of the Creative Commons Attribution 4.0 International license (CC-BY 4.0). (https://creativecommons.org/licenses/by/4.0/legalcode).

## Author Contributions

The author (SC) was solely responsible for the study design, writing the manuscript, analysing and interpreting the data, and final approval of the manuscript. In addition, SC is fully accountable for the work.

This study was funded by an Australian Postgraduate Award (APA) scholarship through the University of Queensland for studies for higher-degree research and The Prince Charles Hospital Foundation 2013 Equipment Grants Program. Even though the original data collection was funded by the National Health and Medical Research Council (NHMRC), this research was not part of the NHMRC-funded project. It was conducted at Queensland University of Technology Medical Engineering Research Facility (QUT-MERF).

## Acknowledgements

Gratitude is extended to members of the sheep ECMO research team at the Critical Care Research Group laboratory that is affiliated with The University of Queensland for their various roles during this research and the staff at QUT-MERF for assistance in the care of the sheep. The author is also thankful to Dr. Jane Charbonneau and SCRIBENDI for editorial support.

